# Identification of transcriptional network disruptions in drug-resistant prostate cancer with TraRe

**DOI:** 10.1101/2022.05.10.491360

**Authors:** Charles Blatti, Jesús de la Fuente, Huanyao Gao, Irene Marín, Zikun Chen, Sihai. D. Zhao, Winston Tan, Richard Weinshilbaum, Krishna R. Kalari, Liewei Wang, Mikel Hernaez

## Abstract

Metastatic castration-resistant prostate cancer (mCRPC) presents very low survival rates due to lack of response or acquired resistance to the available therapies. To date no molecular mechanisms of resistance have been identified, pointing out their complex dynamics. To identify key genes and processes associated with phenotypically-driven regulatory differences, we developed TraRe, a computational method that provides a three-tier analysis: i) at the network level, inferring differentially regulated modules; ii) at the regulon level, identifying regulatory relationships linked to phenotypic differences; and iii) at the single gene level, identifying TFs consistently linked to rewired modules. We applied TraRe (available in Bioconductor with full documentation) to transcriptomic data from 46 mCRPC patients with Abiraterone-response clinical data and uncovered abrogated immune response regulatory modules that showed strong differential regulation in Abi-resistant patients. These modules were replicated in an independent mCRPC study. Further, we experimentally validated key rewiring predictions and their associated transcription factors. Among them, ELK3, MXD1, and MYB were found to have a differential role in cell survival for Abi-response-specific settings. Moreover, we identified the role of ELK3 in cell migration capacity, which could have direct impact on mCRPC. Collectively, these findings shed light on the underlying regulatory mechanisms driving abiraterone response.

## INTRODUCTION

Prostate cancer is among the most frequently diagnosed male cancers in the world^1^. Primary treatments, including surgery or radiation, can achieve a cure for many of the afflicted men^2^. However, a substantial portion is diagnosed with advanced-stage disease or experience disease recurrence. While the mainstay of treatment for these cases has been androgen depletion therapy (ADT), data from clinical trials in patients newly diagnosed with metastatic prostate cancer (mPC)^3, 4^ has demonstrated a survival advantage of 37% for those receiving ADT in combination with other drugs (e.g., enzalutamide, abiraterone acetate, apalutamide and docetaxel) over ADT alone. Additionally, even though ADT is initially effective, > 95% of mPC patients on this treatment relapse, indicative of Castration-Resistant Prostate Cancer (CRPC) development^5^, which has a poor 5-year survival. Due to significantly improved survival rates, the first-line therapy for CRPC patients then becomes either the CYP17A1 inhibitor abiraterone (Abi)^6^ combined with prednisone (AA/P), or the androgen receptor (AR) inhibitor enzalutamide^7^. While results are encouraging, trials with abiraterone or enzalutamide treatment have highlighted two major challenges: 1) pre-existing mechanisms of resistance (primary resistance) preclude responses for nearly half of CRPC patients, and 2) resistance can develop rapidly in initial responders (acquired resistance)^8^.

Together with many other cofactors, the AR transcription factor regulates a series of downstream genes upon ligand binding, followed by translocation into the nucleus. Recently, it was also noted that there are specific co-factors in CRPC tumors not present in normal tissue, which can guide AR to specific gene regions to regulate gene transcription^9, 10^. Additionally, mutations or tumor-specific alterations in pathways could also significantly affect transcription networks and downstream signaling. For example, the *wnt* pathway, which is involved in the regulation of transcription networks, has recently been associated with Abi resistance^11^. Therefore, it is critical to take a comprehensive approach to study the regulation of transcription networks in CRPC and to understand how this regulation might contribute to Abi response.

The regulatory interactions between genes and their corresponding pathways drive various cellular functions that are critical in tumor development and response to therapy^12^. These regulatory relationships, termed Gene Regulatory Networks (GRNs), provide a concise representation of the transcriptional regulatory landscape of the cell^13, 14^. It is well-established that functional GRNs and their products change in response to different conditions, such as cellular DNA damage or environmental stress^15–17^. Hence, the construction and exploration of the topology of GRNs and their constituents are compelling approaches for developing and understanding biological mechanisms.

In the CRPC context, one of the main questions is how the cell changes its behavior in response to drugs, as evidenced by signatures of differentially expressed genes that are a downstream effect of global cell de-regulation in different response groups. Drug treatments can activate different functional pathways in patients based on differences in their underlying GRN^18, 19^. Identifying significant changes among networks from different response groups, also referred to as network *rewirings*, can help discover novel molecular diagnostics and prognostic signatures. While differential gene expression analysis evaluates changes in gene expression under different conditions, the incorporation of network structures and differential network analysis can provide insights that are mechanistically grounded^20^. As an example resulted from differential network analysis, RNA levels of the prostate cancer biomarker AMACR were discovered to have a positive correlation with the tumor suppressor gene PTEN in adjacent normal tissue that was no longer present in prostate tumor samples^21^.

To mechanistically understand how the regulatory differences in transcription networks may contribute to Abi response in metastatic castration-resistant prostate cancer (mCRPC) patients we built a computational framework, termed TraRe, that: i) provides a robust and efficient methodology to uncover gene regulatory modules from high throughput sequencing data; and ii) given those modules, develops an efficient differential network analysis that highlights the transcriptional rewiring associated with a particular phenotype. Specifically, we applied TraRe to the gene expression profiles of 46 mCRPC samples collected before initiation of AA/P treatment from the Prostate Cancer Medically Optimized Genome-Enhanced Therapy (PROMOTE) study that was conducted in a prospective fashion (NCT: 01953640)^11^. First, we showed that TraRe was able to uncover regulatory modules that captured key cellular processes and whose mechanistic rewiring related to Abi response. We matched these findings in a separate cohort of mCRPC patients. Importantly, we were able to uncover transcription factors (ELK3, MYB and MXD1) whose regulatory patterns were significantly associated with response-disrupted regulatory modules. Finally, we experimentally validated in two different cell lines the key rewiring predictions identified from the response-specific regulatory modules.

## MATERIAL AND METHODS

### TraRe: Phenotype-associated transcriptional rewiring

We present TraRe, a computational method to elucidate transcriptional rewiring across phenotypes from RNA sequencing data.

Given a gene expression matrix for *n* samples, the phenotype class of each sample, and a list of transcription factor (TF) regulators, TraRe infers regulatory modules and their phenotype-specific network disruptions (rewiring) by measuring changes in driver-target co-expression among phenotypes (Supplementary Figure S1). In order to infer such rewired networks, TraRe first infers the overall regulatory landscape of the samples, and then uncovers those modules whose topology is rewired between the phenotypes. In addition, TraRe also identifies TFs and regulons (a TF and its corresponding target genes) that are significantly prevalent in the rewired modules.

### Module-based inference of GRNs

TraRe uncovers the GRNs governing the transcriptomic activity of all samples via a module-based approach, where a module represents one or several related biological functions^22^. The module-based approach for inferring GRNs has been shown to be more accurate than inferring a unique global GRN and then isolating individual communities within it^22, 23^.

First, the gene expression data is divided into a matrix *Z* for the expression of target genes and a matrix *X* for the expression of the driver regulators (Supplementary Figure S1A). *Z* is then iteratively partitioned into *K* gene sub-modules where each sub-module’s expression pattern is approximated by a sparse combination of driver gene expression profiles. Specifically, two steps are taken per iteration. In the first step, for each of the *K* inferred sub-modules, the expression profile that is most correlated, on average, with all the sub-module genes is selected as its representative. We denoted this representative profile as the *eigengene y*, as it is the first eigenvector of the sample covariance matrix of the sub-module^22^. Then, the regulatory program of each sub-module (i.e., the set of driver genes regulating it, represented as the sparse vector *β*) is inferred by approximating the expression profile of the eigengene *y* with a sparse linear combination of driver genes. That is,

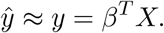

This inference is done via Variational Bayes Spike regression (VBSR)^24^. Within the VBSR, *β*s are computed via a coordinate ascent algorithm which iteratively minimizes the Kullback–Leibler (KL) divergence among the approximate posterior distributions *q̂*_*βj*_ and the complete posterior distribution of the VBSR model *p*(*θ|W*) (see ref.^24^).

The inferred approximation of the eigengene *ŷ* becomes the new sub-module representative profile. In the second step of each iteration, each target gene is reassigned to the sub-module whose representative it is most correlated with. Target and driver genes are iteratively reassigned to sub-modules as aforementioned until no further reassignments occur or when a specified number of iterations is reached.

Note that all target genes get assigned to a particular sub-module. As such, there may be outliers that are not well fit by the inferred regulatory program *β*. Thus, as a final refinement step for each inferred sub-module, the expression of each target gene is modeled from the driver genes in the sub-module via a new VBSR model. This process yields a weighted, undirected bipartite graph for each sub-module representing its individual GRN drawing specific connections between target genes and their inferred regulators. During this step, target genes within sub-modules can be rejected during the fitting. Therefore, outlier target genes will be dropped out due to lack of good model fit, leading to refined sub-modules with consistent GRNs (Supplementary Figure S1B).

At the end of this stage, TraRe has generated a set of refined sub-modules depicting the regulatory landscape of the provided input gene expression data *Z*. It is worth mentioning that TraRe’s package includes two additional regression models to link targets and drivers. These models are Linear Regression Model (LM) and Least Absolute Shrinkage and Selection Operator (LASSO).

### Uncovering of rewired GRNs

After inferring the refined sub-modules as described above, TraRe assesses the refined sub-modules independently for evidence of network rewiring between phenotypes. Hence, the description below focuses on a single sub-module.

Given the gene expression matrix associated with each sub-module (Supplementary Figure S1C), the gene-gene covariance matrix associated to each binary-labeled phenotype is computed separately, namely Σ^0^ and Σ^1^. Then, the distance between the two covariance matrices is computed as the Frobenius norm of the difference between both of them:

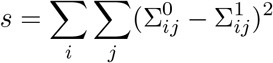

This distance, which we term the *rewiring score* is used as a statistic to measure the dissimilarity between the covariance matrices, and thus, the differential regulatory effect of the sub-modules across phenotypes. In order to assess the statistical significance of the rewiring score *s* a permutation test is performed, where the rewiring score is evaluated against a null distribution H_0_ generated via random permutations of the phenotype labeling (Supplementary Figure S1D). Thus, a sub-module is said to be rewired if its rewiring score *s* satisfies

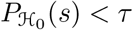

where *τ* is a user-defined threshold, set by default to 0.05. Note that a significant dissimilarity over the covariance matrices may indicate fundamental underlying disruptions in gene co-expression between phenotypes.

### Uncovering robust rewired GRNs across multiple runs

Unraveling GRNs from transcriptomic data, also known as reverse engineering the transcriptome, is a complex problem that generally requires heuristic algorithms. Thus, every run of TraRe (with a unique random seed) generates slightly different results. Hence, towards accounting for the implicit variability of TraRe and increasing its generalization capabilities, the module generation process is repeated *B*times using different subsets of 80% of the samples. Refined sub-modules that generalize well across different runs and different samples should be similarly captured in each run. Hence, similar sub-modules across runs will cluster together, giving rise to consistent regulatory modules in which spurious patterns coming from a specific group of samples are dropped out.

After inferring the rewired sub-modules from all runs as mentioned above, an similarity matrix is built so that similar rewired sub-modules are grouped together through hierarchical clustering. Specifically, the *ij* similarity value of this matrix is defined based on the log significance (using the hypergeometric test) of the overlap of the pair of submodules:

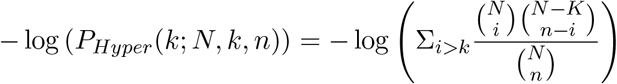

where *N* is the total number of genes in the dataset, *K* and *n* are the number of genes in the rewired sub-modules *j* and *i*, respectively, and *k* is the number of common genes between both rewired sub-modules. Then, hierarchical clustering is performed to group together very similar rewired sub-modules that have spawned across different runs, yielding robust rewired regulatory modules (Supplementary Figure S1F).

### Uncovering rewiring-specific driver genes

We classify transcription factors (TFs) as rewired-specific if they appear statistically significantly more times in rewired sub-modules than in non-rewired ones. We set the re-runs of TraRe to *B* = 50, and computed an enrichment p-value of the over-presence of a given TF in rewired sub-modules using a hypergeometric test with

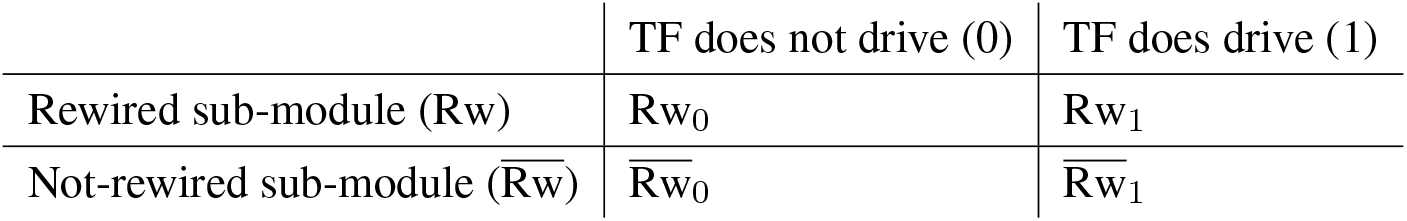

as the associated contingency table. Here, Rw and Rw are the non-rewired and rewired sub-modules, respectively, and their sub-indexes point to whether the transcription factor is (1) or is not (0) a driver in the sub-module, respectively.

In addition, the Odds Ratio test is used to further assess the *rewiring specificity* of the driver genes. Specifically, the odds ratio is computed as

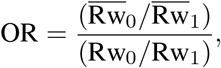

where the Haldane^25^-Anscombe^26^ correction, is applied due to the sparsity of the contingency table. Finally, rewired-specific TFs are filtered by a threshold (set to 0.05 for the hypergeometric test and to 1 for the OR test by default) and sorted according to the associated p-value.

### Uncovering rewiring-specific regulons

As opposed to the previous analysis where we evaluated the specificity of driver genes within rewired sub-modules, target genes can not be evaluated using the aforementioned approach, as they are assigned to a unique sub-module in each run. Therefore, we chose to assess target genes via the regulons of rewired sub-modules that they are associated with.

From the inferred rewired sub-modules, we first extract all regulons, which are composed by a driver gene and its associated target genes. Note that these regulons form simple networks and hence can be tested for their potential rewiring applying the same test as we did for the inferred sub-modules (see above).

Therefore, each regulon will have an associated p-value (adjusted with the Benjamini-Hochberg multiple hypothesis correction to compensate for spurious significant p-values), which is informative about the network’s disruption significance within that set of targets associated with a single driver gene. Only significant rewired regulons (corrected p-values below 0.05) are taken henceforward.

Along the 50 runs, several regulons may be led by the same TF and contain repeated associated targets; thus, we merged those regulons under the same driver gene. Then, we computed the multiplicity for each target, defined as the number of times they appear within the unified regulons, and its significance, defined as the Fisher’s combined probability test over the p-values associated with the regulons it appears in. Targets whose p-values are below a threshold (set to 0.05 by default) are dropped out and the remaining targets are sorted by the multiplicity. The filtering process can drop out every target within a regulon. If this is the case, the associated regulon is removed.

Additionally, we computed the overall multiplicity of the targets across all the rewired regulons (not only in those under the same driver gene) and calculated the associated p-values with the aforementioned Fisher’s exact test. A list of rewired targets, those whose p-value is below the threshold (set to 0.05 by default) is generated.

### Visualization of the results in TraRe

TraRe includes a set of functions that allow the visualization of the results obtained in the stages of the method in an html report. Moreover it outputs different summary files and figures that include the sub-module similarity matrix in the form of a clustered heatmap with the corresponding dendrogram and the modeled bipartite graphs of the different robust, rewired regulatory modules that the clustering algorithm identifies.

### Datasets

We have tested our method using clinical and molecular data from two metastatic castration-resistant prostate cancer (mCRPC) datasets: 1) Mayo Clinic clinical trial (NCT: 01953640), Prostate Cancer Medically Optimized Genome-Enhanced Therapy (PROMOTE) and 2) the Abida *et al.*^27^ study from the work of Stand Up 2 Cancer-Prostate Cancer Foundation Prostate Dream Team (SU2C).

### PROMOTE study

The PROMOTE study aimed to identify the genomic alterations associated with resistance to abiraterone acetate/prednisone (AA/P) treatment^11^. The primary resistance was determined at 12 weeks of therapy using several criteria for progression such as serum prostate-specific antigen (PSA) measurement, bone, and computerized tomography imaging and symptom assessments. In the original PROMOTE study^11^, whole transcriptome sequencing (RNAseq) was performed using the TruSeq poly-A selection library and Illumina HiSeq 2000 instruments. For this manuscript, we obtained the RNA sequencing from 64 tumor samples and normalized the gene counts using conditional quantile normalization. This gene expression data came from 46 bone and 18 soft tissue metastatic sites before the AA/P treatment began. We performed our TraRe analysis on only the 46 bone samples, from which 29 and 17 were Abi responders (R) and non-responders (NR), respectively.

### Stand Up 2 Cancer Study

In addition to the PROMOTE study, we ran TraRe on the ‘Stand Up 2 Cancer’ study data, which also consists of patients with mCRPC disease. We downloaded the clinical data and sequencing data from the database of Genotypes and Phenotypes (dbGaP) (accession code,phs000915.v2.p2) and the cBioPortal^28^ Public Datahub (https://github.com/cBioPortal/datahub/tree/master/public/prad_su2c_2019). Since the PROMOTE RNA sequencing study data was generated using TruSeq poly-A selection library, in the SU2C study, we filtered out the data generated using a hybridization capture-based library. We obtained the RNAseq gene expression data (FPKM) from 270 SU2C samples generated using the TruSeq poly-A selection library, and removed genes with no expression in more than 90% of the samples. The FPKM data was then quantile normalized and log2 transformed.

### Study harmonization

To harmonize the information of both studies, only those patients treated with Abi in the SU2C study were selected. Specifically, 121 patients whose transcriptomic data was available prior to Abi treatment were used to infer the regulatory modules.

Note that the number of genes examined from the two matrices differ. Therefore, only common genes (12791) have been selected when evaluating both datasets jointly. In all other analyses, the original features (genes) have been maintained. In order to partition the genes in this analysis between regulators and targets, we defined the regulators as all genes which match the list of transcription factors selected from the Human Protein Atlas 2020 available from v19.3^29^, and the targets as all of the remaining genes.

### Enrichment analysis

We used the CommunityAMARETTO algorithm^30^ to cluster together similar inferred sub-modules and to assign them a biological meaning. Using an hypergeometric test with a FDR < 1*e^−^*^6^, a module-based network of overlapping sub-modules is created and then significantly connected subnetworks are identified using the Girvan-Newman algorithm^31^ and clustered into regulatory modules, which are assessed for enrichment (hypergeometric test) with known functional categories in the curated (C2) and Hallmark (H) gene sets of Molecular Signatures Database (MSigDB^32^, v5.2) For the biological processes (BP) Ontology (C5, MSigDB) enrichment analysis of the genes included in the rewiring-specific driver list, regulons and rewired regulatory modules, the following R packages were used: clusterProfiler (v4.0.5), org.Hs.eg.db (v3.13.0), AnnotationDbi (v1.54.1) and DOSE (v3.18.3).

### Simulated data generation

To evaluate our methods, we generated a simulated dataset with underlying regulatory modules, some of which were rewired, and its corresponding simulated gene expression data. For our simulated evaluations, 10 sub-modules were generated through the following process. Driver genes from the PROMOTE’s dataset were selected and sampled such that a linear combination of them defined each sub-module mean expression profile *µ*, with an average number of drivers within each sub-module set to 5. Target gene expression profiles were generated by sampling 200 times from a Multivariate Gaussian distribution *N*(*µ, σI*). As *σ* was selected to be much lower than the variance of *µ*, sub-module’s genes were strongly correlated (*>* 0.7) to each other. In order to simulate rewiring within sub-modules, targets genes and TFs within samples from one phenotype class were decorrelated (whitening), imposing a clear disruption among phenotype’s covariance sub-matrices. From the 10 generated sub-modules, sub-module 1, 4 and 7 were randomly chosen to be rewired sub-modules and the seed was fixed to maintain consistency across scenarios.

TraRe’s inference method was applied to the simulated sub-modules selecting VBSR, LM and LASSO as the fitting models. This process was repeated 5 times with random 80% of the samples each, to increase the inference generalization. Finally, we clustered the inferred sub-modules yielding the final inferred regulatory modules. To that end we developed a decoder-encoder-based pipeline where two graphs were built.

First, simulated and inferred sub-modules were paired and used to build a bipartite graph where the Jaccard index was used as the similarity metric to relate the simulated and inferred sub-modules. From these sub-module pairs, an alluvial plot was drawn showing the flow of simulated-to-inferred sub-modules across the 5 runs (*decoding*). A second graph was built over inferred sub-modules using the aforementioned similarity metric, where *edge betweenness* clustering (from R’s igraph library) was applied to group similar inferred sub-modules (nodes) from each run (*encoding*). A second alluvial plot was drawn to show connections from inferred to clustered regulatory modules. These two alluvial plots were connected to visualize the complete flow from simulated sub-modules to clustered inferred regulatory modules. The intuition behind the sub-module’s clustering is that, if simulated sub-modules are correctly uncovered on every run, a graph clustering technique should group them together across runs, recovering the original regulatory modules.

In order to assess TraRe’s inferring capability within the simulated data, three scenarios were considered, where the three fitted models were benchmarked using classification metrics. On each scenario, ROC curves were calculated when varying: i) the noise variance *σ* = [0.01, 0.4, 0.8, 1.2, 1.6, 2, 2.4, 2.8, 3.2, 3.6, 4, 4.4, 4.8, 10, 25, 50, 100], ii) the p-value threshold prior to Bonferroni correction (0.02 to 0.14 for VBSR and LM, 1 to 0 for LASSO), and iii) the number of samples (10, 16, 28, 38, 46) in the generated bulk RNAseq expression matrix. Note that, in the absence of p-value for the LASSO model, the Lambda parameter has been used.

Finally, we evaluated VBSR, LASSO and LM for a fixed *σ* (1.3), p-value threshold (0.05) and number of samples (46), which are the default parameters henceforth. For each driver gene, simulated-inferred sub-modules pairs were evaluated by measuring true positives (TP), if a driver is contained in both the simulated and the inferred sub-module, false positives (FP), if there is a driver in the inferred but not in the simulated and false negative (FN), if there is a driver in the inferred but not in the simulated sub-module. Moreover, a linear model was used to fit the inferred regulatory program of TFs based on the simulated regulatory programs. *R*^2^ adjusted, Precision and Recall parameters were used for model assessment.

### Cell culture

22Rv1 and LNCaP cells purchased from ATCC were cultured in RPMI1640 medium (Gibco, Grand Island, NY) supplemented with 10% FBS (Sigma) and 1% Pen-Strep. To develop Abi-resistant cell lines (LNCaP-AR or 22Rv1-AR), cells were maintained in the medium supplemented with 5 µM of abiraterone (Selleck Chemicals, Houston, TX) until viability reached over 95%. Abi-resistance was validated by proliferation assay. For knockdown experiments, cells were seeded in regular RPMI1640 in 6-well plates, and were transfected using siRNA against ELK3, MXD1, MYB, ZNF3, ZNF91 or non-targeting siRNA (Horizon Discovery) by RNAiMax (Thermo Fisher Scientific) according to manufacturer’s protocol. siRNA sequences can be found in Table S10

### Proliferation assay

36 hours before Abi treatment, cells were seeded in phenol red-free RPMI1640 (Gibco, Grand Island, NY) supplemented with 10% charcoal stripped FBS (Sigma) in 96-well plates, and 50 nM pregnenolone was added after 16 hours. Cells were then treated with Abi or vehicle and monitored for proliferation at day 0, 2, 4 and 6 after initiation of treatment by Cyquant direct assay (Thermo). For proliferation after knockdown, cells were trypsinized 24 hours after transfection and reseeded in 96-wells plates. Proliferation was monitored at day 1, 3, 5 and 7 after reseeding.

### qRT-PCR

Total RNA for qRT-PCR was extracted from cell lines 48 hours after knockdown using Quick-RNA MiniPrep Kit (Zymo Research, Irvine, CA) according to the manufacturer’s instructions. qRT-PCR was performed using the *Power* SYBR® Green RNA-to-C_T_™ 1-Step Kit (Life Technologies, Grand Island, NY) and QuantiTect® (QIAGEN, Germantown, MD) or PrimeTime® (IDT, Inc., Coralville, Iowa) pre-designed qPCR primers (IDT Coralville, IA). Gene expression analyses were performed using the ΔΔCt method, and GAPDH was used as the internal reference. Two independent experiments were performed. Primer sequences are in Supplementary Table S9.

### Wound-healing assay

LNCaP, 22Rv1 or their Abi-resistant cells were seeded in 6-well plates at 6 x 10^5^ density, and transfected with siRNA targeting ELK3 or scrambled siRNA control. 24 hours after transfection, a scratch wound was made by pipet tip. Images were then taken immediately and after 24 hours.

## RESULTS

### Overview of the study

We applied the developed TraRe framework (Figure 1) to the gene expression profiles of 46 mCRPC baseline pretreatment samples from the PROMOTE study. Abi response annotation for all patients was measured at three months after treatment initiation and patients were separated into two groups: treatment responders (R) and non-responders (NR)^11^. Given the availability of pre-Abi treatment gene expression of the PROMOTE samples and a set of known transcription factors (TFs), TraRe first performed an iterative clustering process, which partitions target genes into different sub-modules, whose expression can be modeled by the expression of a sparse set of TF regulators (Supplementary Figure S1B, see Methods). In order to overcome sensitivity to outliers in the data, TraRe was run several times, each run randomly selecting a different subset of the patient samples (80-20 split), as well as a random initialization of the parameters. We then created our regulatory modules by clustering the similar inferred sub-modules across the multiple runs. We reasoned that regulatory modules sharing common sets of genes across subsampled runs would uncover a regulatory landscape of biological processes that are not a consequence of overfitting to a specific set of patients, and therefore, better generalize to the disease population. To validate this hypothesis, regulatory modules were independently inferred on the SU2C^27^ dataset consisting of 121 mCRPC patients (see Methods). Module-oriented community detection and functional annotation methods^30^ showed that indeed our method found shared regulatory modules between PROMOTE and SU2C.

**Figure 1:**
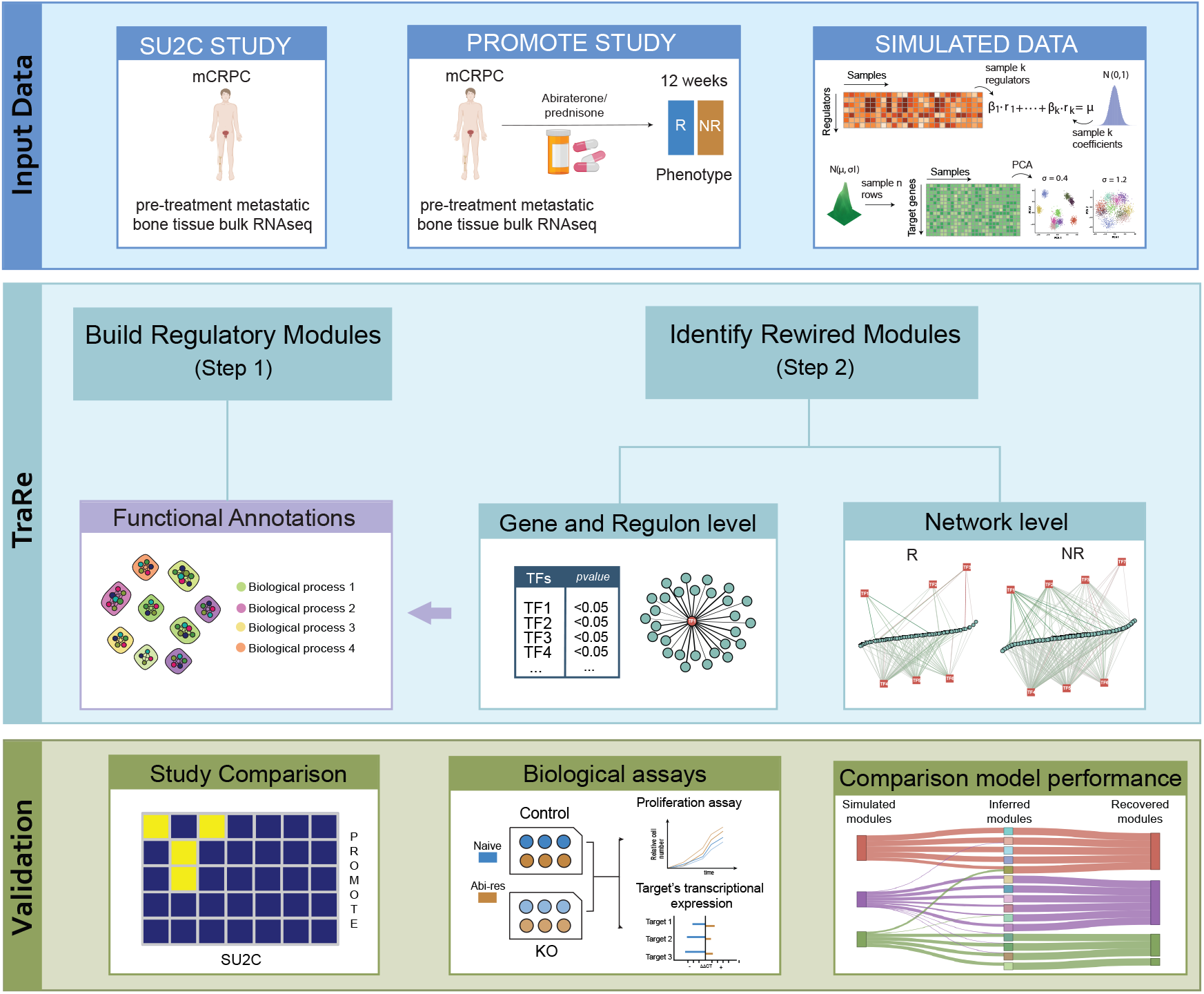
TraRe’s workflow overview. **Input Data:** Datasets used in the study: bulk RNAseq data from Stand Up 2 Cancer (SU2C) and Promote studies of mCRPC cohorts previous to abiraterone treatment; Simulated Data: Generation and visualization, via PCA, of the simulated modules. Driver genes from the PROMOTE’s dataset were selected and sampled such that a linear combination of them defined each simulated module mean *µ*. For each module, target genes were generated by sampling from a multivariate Gaussian distribution. **TraRe:** Build Regulatory Modules (Step 1); Functional annotations: over-representation analysis of Regulatory Modules, regulons and TFs with curated genesets associated with biological functions.Identify Rewired Modules (Step 2); Gene and Regulon level: ranked TFs associated with response and TF-target association in regulons; Network level: phenotype specific regulatory module analysis. **Validation:** Study comparison: Comparative analysis of regulatory modules of PROMOTE and SUC2 study. Biological assays: *in vitro* TF knock-down studies with naïve and Abi-resistant cell lines to assess proliferation and target mRNA expression; Comparison model performance: Alluvial plot built to compare fitting models across simulated GRNs.

We next aimed at unraveling those regulatory modules from PROMOTE showing differential mechanistic regulation between Abi R and NR. Specifically, for each sub-module generated across runs, we performed a statistical *rewiring* test based on the differences between Frobenious norms of the gene-gene covariance matrices to identify those whose co-expression patterns were significantly different between the two response groups (Supplementary Figure S1C-D). We considered a robust regulatory module to be rewired if at least 40% of its sub-modules showed significant rewiring by our test.

We then sought more insights into the key TFs potentially driving the response-specific rewiring process. We used Bayesian sparse regression^13^ models in each sub-module to associate each TF to a specific set of target genes, known as regulons (see Methods). We, thus, were able to rank TFs and their target genes by their associations with the response-specific rewiring process. Finally, we closely investigated the regulons of key rewired regulatory modules in order to uncover specific regulatory relationships between TFs and targets that behaved distinctly between response groups.

Therefore, in this study we developed TraRe, a computational method that provides a three-tier analysis to identify key biological processes and genes associated with phenotypically-driven regulatory differences: i) at the module level, by inferring differentially regulated modules; ii) at the regulon level, by identifying specific regulatory relationships that may be associated with phenotypic differences; and iii) at the single gene level by identifying TFs consistently associated with rewired modules. We concluded this study by experimentally validating a subset of our findings with TF knockdown in Abi-resistant prostate cancer cell lines. We also shared our findings on the ideal selection of regression models for uncovering phenotype-associated regulatory rewiring disruptions, as well as the success of the proposed rewiring statistical tests on varied simulated benchmark datasets.

### TraRe recapitulates regulatory programs associated with metastatic Castration-Resistant Prostate Cancer

We first focused on uncovering the regulatory landscape and associated key biological processes of mCRPC. To that end, TraRe was ran on the 46 PROMOTE samples from pre AA/P-treatment bone biopsies. TraRe was run 10 times with each time sampling without replacement 80% of the samples, and each run set to infer up to 100 sub-modules (*K* = 100). This number of sub-modules was chosen following previous works^22, 33, 34^. Nevertheless, we tested different values of *K* and lower numbers yielded sub-modules with impractical sizes (*K* = 50), whereas larger numbers (*K* = {200, 300, 500}) generated sub-modules of sizes unfit for gene set enrichment analysis (Supplementary Figure S2). Across all runs, TraRe inferred a total of 835 sub-modules, and identified 83 distinct communities of overlapping sub-modules (hypergeometric test, FDR < 10*^−^*^5^) (see Methods). Of the 83, 38 were identified as (robust) regulatory modules as they contained sub-modules shared among at least half of the 10 runs (See Supplementary Tables S1 to S3 on pages 5–8). These 38 regulatory modules were functionally annotated with cAMARETTO software^30^ using curated C2 and hallmark gene set enrichment analysis (Figure 2A, see Methods, Supplementary zip file). Some regulatory modules, such as P-M15 and P-M18, contained genes enriched in gene sets related to cell constitutive and housekeeping functions such as G protein-coupled receptors (GPCR) and peptide-ligand binding receptors (P-M15 and P-M18) or cell-cycle and transcription (P-M15), and are thus not explicitly associated to Abi response (Figure 2C).

**Figure 2:**
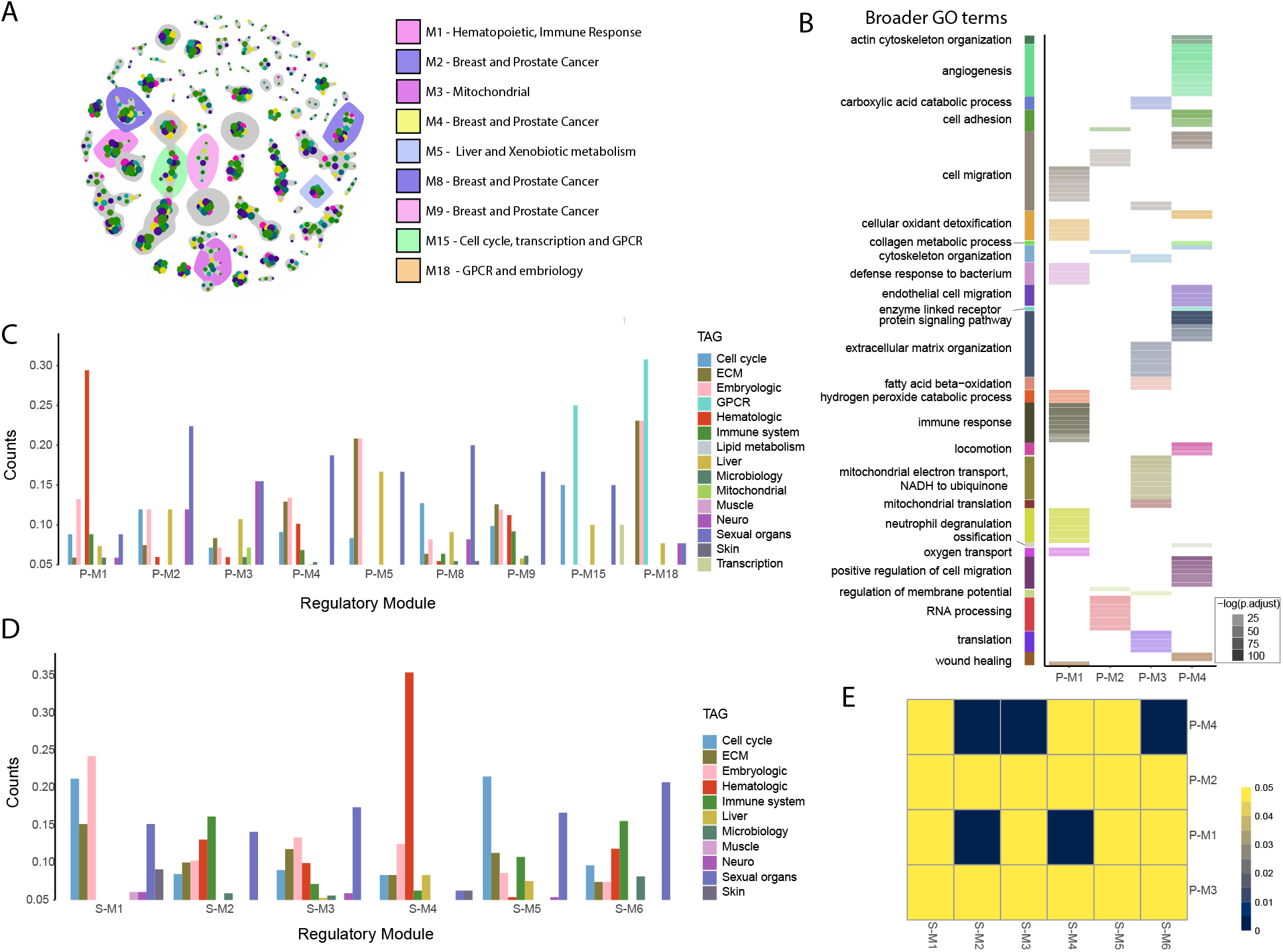
TraRe recapitulates complex regulatory interactions on metastatic Castration-Resistant Prostate Cancer. **A:** Some regulatory modules identified with cAMARETTO software on inferred GRNs with TraRe. **B:** Gene Ontology gene sets of biological processes terms in which the rewired regulatory modules are enriched. **C-D:** Categorization of functional annotation enrichment found with cAMARETTO software in selected PROMOTE’s (**C**) and SU2C’s (**D**) regulatory modules. **E**: Similarity heatmap (Jaccard index) between rewired regulatory modules in PROMOTE study and overlapped SU2C study regulatory modules. ECM: Extracellular matrix.

We then sought to test our regulatory modules for differential mechanistic regulation between 29 Abi R and 17 NR in the PROMOTE dataset. To achieve this goal, we applied our rewiring statistical test (see Methods) to all sub-modules separately. We found that 4 out of the 38 regulatory modules (P-M1 to P-M4) were enriched with rewired sub-modules. These rewired regulatory modules were characteristically enriched for: embryological and hematopoietic-related gene sets (P-M1), mitochondrial gene sets (P-M3) and different gene sets related to cancer and reproductive system organs (P-M2 and P-M4) (Figure 2C). In accordance with these findings, these regulatory modules were also enriched in GO Biological Process terms related with the functions mentioned above: mitochondrial electron transport in P-M3 and activation of different hematologic cells and immune response in P-M1 (neutrophil, leukocytes, etc.). In the case of the other two regulatory modules, the most important GO Biological Processes associated functions were related to RNA processing (P-M2) and cell migration, angiogenesis and extracellular matrix organization (P-M4) (Figure 2B, Supplementary Excel).

We then examined the TFs driving the global expression patterns of these rewired regulatory modules (Supplementary Table S1). We found that TFs CEBPE, GATA1, KLF1 and MYB, which drove regulatory module P-M1, are the master regulators of granulocytes differentiation^35, 36^, erythroid development^37^ and other hematopoietic and immune pathways^38, 39^. In regulatory module P-M2, the key driver gene ELK1 (a nuclear target of the MAPK signaling cascade) has a potential role in the activation of AR signaling of growth genes in prostate cancer^40^, while SREBF2 is overexpressed in more aggressive prostate cancer cell lines and metastatic prostate cancer^41^. The important driver gene from regulatory module P-M3, SMAD7 was shown to also promote migration and invasion in prostate cancer cells^42^, and SOX8 is known to be involved in cisplatin-chemoresistance in different cancers^43, 44^. Finally, in regulatory module P-M4, SNAI2, is involved in epithelial-mesenchymal transitions in different cancer types^45, 46^, has anti-apoptotic activity^47^ and it is a metastasis-promoting TF in breast cancer progression^48^.

Other regulatory modules containing rewired sub-modules (P-M5, P-M8 and P-M9) were found to be enriched with genes down-regulated in prostate cancer samples (as well as different cancer types) (Figure 2A). The driver gene ETV5 from P-M9 is involved in the AR signaling pathway associated with invasion and it has been identified to participate in rare gene fusion permutations in prostate cancer^49^. Interestingly, P-M5, with two rewired sub-modules, was found to be enriched with liver and xenobiotic metabolism-related pathways such as HNF1A, cytochrome P450 and glucuronidation, affecting drug response and efficacy.

To further validate the regulatory modules uncovered by TraRe in the PROMOTE dataset we ran TraRe on the 121 mCRPC samples of the SU2C dataset (see Methods) with the same run settings. We found 49 robust regulatory modules in the SU2C dataset. Six of these SU2C robust regulatory modules were highly overlapping with our regulatory modules identified from the PROMOTE dataset. These six modules were enriched with cell cycle related genes that were also found to be differentially regulated in different cancer cells (SU2C S-M21), haematologic (S-M9), embryological and cell cycle (S-M4), breast and prostate cancer gene sets (S-M2, S-M3 and S-M6) (Figure 2D). We were especially interested in the significant overlaps of the six SU2C modules with our rewired regulatory modules from PROMOTE (Figure 2E). P-M1 from PROMOTE was closely related with S-M4 from SU2C, as expected by their common gene set enrichment analysis (hematologic gene sets, Figure 2D), and also with S-M2, a module enriched with immune gene sets. Similarly, P-M4 (related with breast and prostate cancer) from PROMOTE was related to the modules S-M2, S-M3, and S-M6 from SU2C, primarily enriched with the same cancer gene sets (Figure 2D). As a final note, we were unable to compare if the SU2C modules have similar associations with response to that of PROMOTE due to the lack of an analogous clinical response measurement after three months of treatment. Thus, TraRe was able to generate biologically meaningful robust regulatory modules from PROMOTE data that were consistent across multiple runs (Figure 2A). Specifically, the highlighted PROMOTE regulatory modules P-M1, P-M2, P-M3 and P-M4, which were enriched with differentially rewired sub-modules between Abi R and NR, captured novel transcription regulation that may help elucidate the foundations underpinning abiraterone sensitivity and resistance. Similar modules were also observed in an independent mCRPC study (SU2C).

### TraRe identifies transcription factors playing a key role driving abiraterone response

Next, we investigated individual TFs that may be driving significant differences in the regulatory networks between the Abi treatment responder (R) and non-responder (NR) groups. We hypothesized that the rewired regulatory modules inferred in the previous section are likely to be regulated by TFs that themselves might not be rewired between R and NR as the conserved behaviors of these TFs could underpin the module that they regulate (as we show in the following section). To increase the statistical power required when working at single-gene resolution, we ran TraRe 50 times on the 46 PROMOTE samples with the usual settings, yielding 4088 sub-modules, from which 225 were marked as significantly rewired by TraRe. We then selected those TFs that were identified as the “drivers” driving the core sub-module expression and that were highly enriched in rewired sub-modules (see Methods). The resulting ranking contained 33 TFs that likely played a key role in the driving rewiring of the GRNs associated with the different Abi treatment outcomes (Table 1).

**Table 1:**
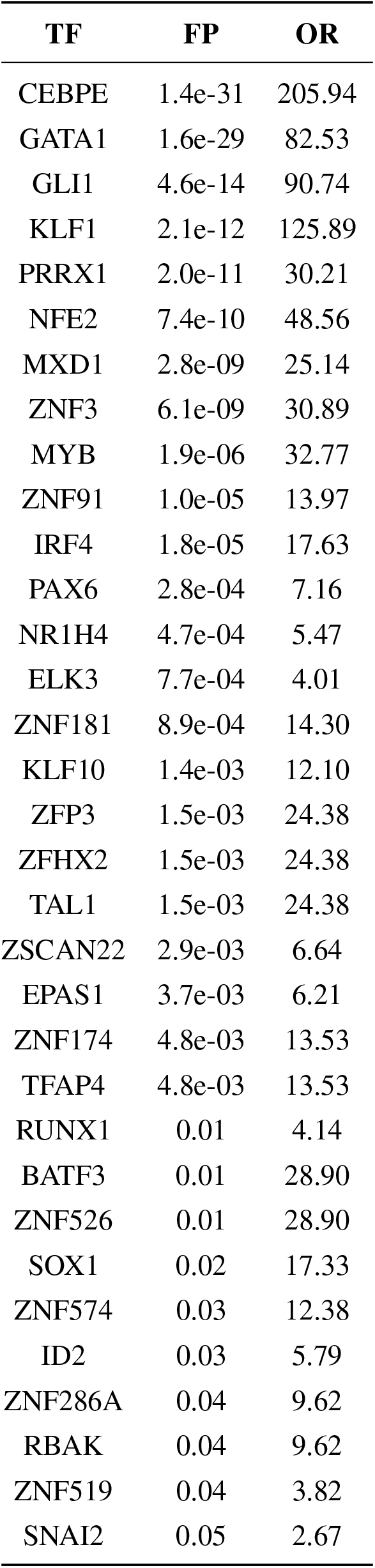
Rewired TFs. TF: Transcription Factors; FP: Fisher’s p-value; OR: Odds Ratio.

| **TF** | **FP** | **OR** |
| --- | --- | --- |
| CEBPE | 1.4e-31 | 205.94 |
| GATA1 | 1.6e-29 | 82.53 |
| GLI1 | 4.6e-14 | 90.74 |
| KLF1 | 2.1e-12 | 125.89 |
| PRRX1 | 2.0e-11 | 30.21 |
| NFE2 | 7.4e-10 | 48.56 |
| MXD1 | 2.8e-09 | 25.14 |
| ZNF3 | 6.1e-09 | 30.89 |
| MYB | 1.9e-06 | 32.77 |
| ZNF91 | 1.0e-05 | 13.97 |
| IRF4 | 1.8e-05 | 17.63 |
| PAX6 | 2.8e-04 | 7.16 |
| NR1H4 | 4.7e-04 | 5.47 |
| ELK3 | 7.7e-04 | 4.01 |
| ZNF181 | 8.9e-04 | 14.30 |
| KLF10 | 1.4e-03 | 12.10 |
| ZFP3 | 1.5e-03 | 24.38 |
| ZFHX2 | 1.5e-03 | 24.38 |
| TAL1 | 1.5e-03 | 24.38 |
| ZSCAN22 | 2.9e-03 | 6.64 |
| EPAS1 | 3.7e-03 | 6.21 |
| ZNF174 | 4.8e-03 | 13.53 |
| TFAP4 | 4.8e-03 | 13.53 |
| RUNX1 | 0.01 | 4.14 |
| BATF3 | 0.01 | 28.90 |
| ZNF526 | 0.01 | 28.90 |
| SOX1 | 0.02 | 17.33 |
| ZNF574 | 0.03 | 12.38 |
| ID2 | 0.03 | 5.79 |
| ZNF286A | 0.04 | 9.62 |
| RBAK | 0.04 | 9.62 |
| ZNF519 | 0.04 | 3.82 |
| SNAI2 | 0.05 | 2.67 |

Interestingly, the highest ranked TFs were those also found in the most significantly rewired regulatory module (P-M1), such as CEBPE, GATA1 or KLF1. Furthermore, using the STRING database^50^, we found that the interaction network between the top ranked TFs (Figure 3A) showed multiple connections between those TFs identified in P-M1. The top enriched pathways associated with those TFs included regulation of hemopoiesis and different blood cell differentiation pathways, specifically myeloid cells, indicating a strong functional relation among these TFs (Figure 3B, Supplementary Table Excel). These results showed the robustness of the rewiring analysis of the P-M1 since its drivers were also among the top significant rewired TFs, highlighting the capability of TraRe to infer GRNs with drivers and target genes that are associated with the same pathways and biological functions.

**Figure 3:**
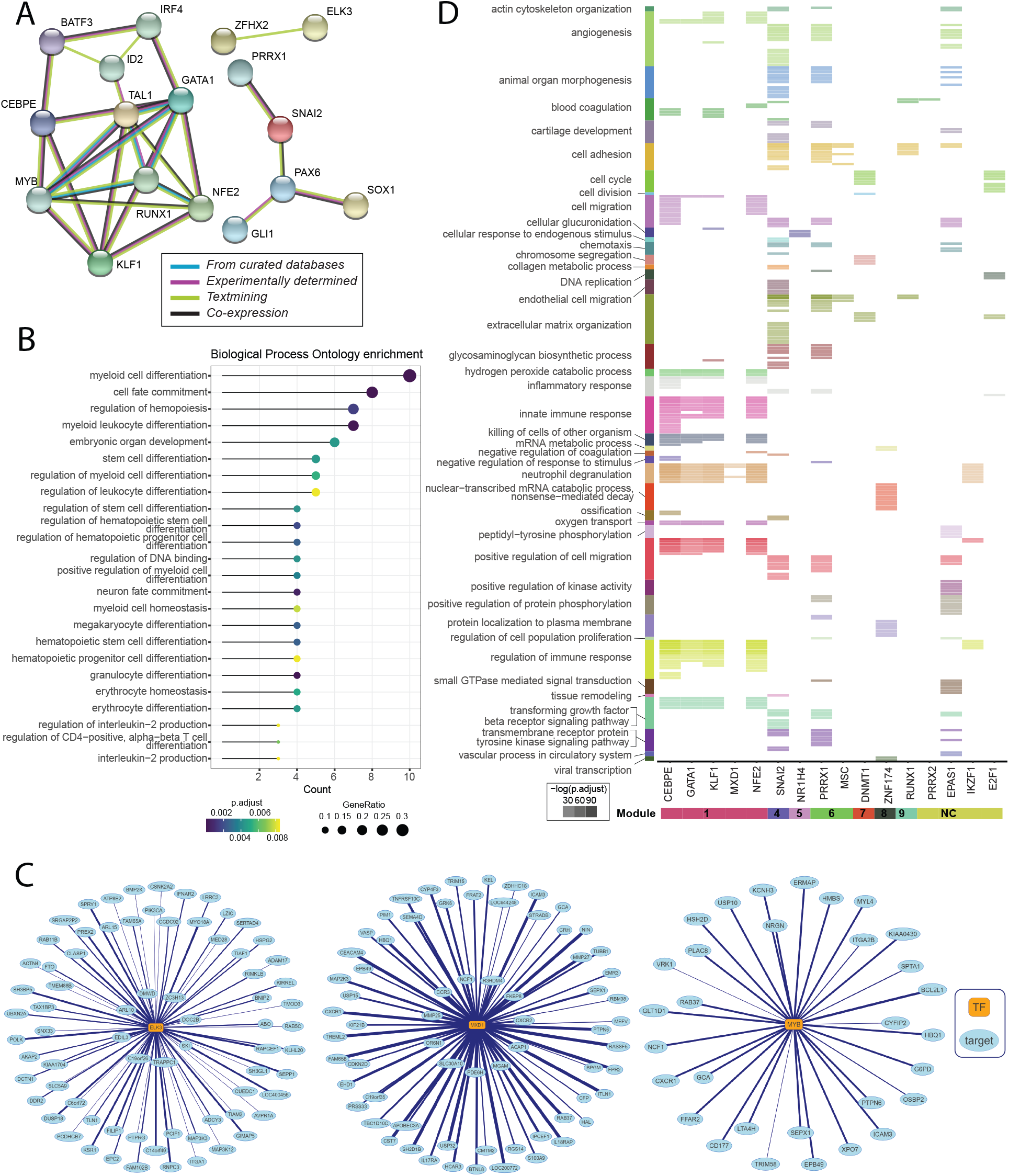
TraRe unraveled Transcription Factors playing a key role in Abiraterone Response of mCRPC patients. **A:** Protein-protein association network of top rewired TFs. Relationships are based on experimental data (purple), curated databases (blue), co-expression (black) and text mining (yellow). 21 out of 33 nodes contain an edge. Pprotein-Protein Interaction enrichment p-value: < 1.0e-16. Figure generated with STRING Database^50^ web tool v.11.0 (Permanent hyperlink). **B:** Enrichment analysis of Gene Ontology Biological Process top terms of the rewired TFs. **C:** Enrichment analysis on Gene Ontology Biological Process terms of the targets in the regulons. Gene Ontology analysis was performed by using R package clusterProfiler (v3.18.1) and org.Hs.eg.db. **D:** Graphical representation of three regulons (edges filtered for better visualization)

In order to provide a better understanding of the rewired regulatory modules that were generated by TraRe, we explored the relationships between TFs and target genes at the regulon level. A regulon is defined as a set of target genes associated with one driver gene, in this case, a TF. Therefore, we identified robust rewired regulons across the rewired modules (see Methods, Supplementary Excel). A visual example of these is shown in Figure 3C for three TFs selected for further biological validation, ELK3, MYB and MXD1. In total, there were 50 significant rewired regulons and almost all of the top TFs described in the previous section, such as MYB or MXD1, were master regulators of these regulons. Functional enrichment analysis with GO terms of the regulons shows that the most frequent terms outputted from the over-representation analysis with biological processes were under broader terms related with regulation of immune response and cell migration, extracellular matrix organization, cell adhesion and angiogenesis as well as different receptor signaling pathways. Consistent with the results obtained in the functional annotation of the rewired regulatory modules, the regulons of the master regulators presented in P-M1, such as CEBPE, GATA or NFE2, were enriched in gene sets related with neutrophils and innate immune response (Figure 3D, Supplementary Excel).

In summary, we showed that TraRe was able to uncover key TFs and regulatory programs (regulons) related to ABI response and associate them with functional pathways. Furthermore, these analyses at both single-gene (TF) level and regulon level complemented the findings at module level (previous section).

### TraRe uncovers regulatory changes between response-specific GRNs

In previous sections, we showed that TraRe was able to recapitulate known biological processes in mCRPC through the inference of regulatory modules. Some of those modules showed significant evidence of containing Abi response-specific gene regulation patterns, indicating that potential perturbations in the GRNs underlying their biological processes could relate to the success of Abi treatment. To further delve into these altered regulatory processes, we used TraRe to find regulatory modules formed by consistently rewired GRNs whose rewiring was associated with Abi response.

To better understand how the interactions between TFs and their target genes differ between the samples from Abi R and NR, we focused on the significantly rewired and highly robust regulatory module PROMOTE module P-M1 (see Supplementary Table S1). P-M1 was composed of 11 sub-modules from the 10 original runs of TraRe. Note that each run would ideally contribute at least one sub-module to the regulatory module P-M1, indicating that P-M1 is consistently discovered across different runs. Indeed most runs contributed with one sub-module to P-M1, with the first and seventh runs splitting the co-expressed genes into two separate sub-modules (Supplementary Figure S9). All 11 of the individual sub-modules were statistically significantly rewired with respect to Abi response, with a total of 9 driving regulators (TFs) found (Supplementary Figure S12); GATA1, CEBPE, FOXN1, MYB, TAL1, GLI1, KLF1, MXD1, and NFE2. We merged the 619 unique genes of the 11 individual sub-modules and used the Variational Bayes Spike Regression (VBSR, see Methods) model to redraw the combined GRN using all 46 bone samples (Figure 4A, right). The resulting network successfully modeled the expression of 222 of the target genes using the 9 different regulators. We also created VBSR-based GRNs for the 17 NR and 29 R patients separately (Figure 4A left and center) to examine the differences in the underlying regulatory networks for the two groups (similar results were observed using all patient samples and alternative regression methods, see Supplementary Note S1.1 and Supplementary Table S12).

**Figure 4:**
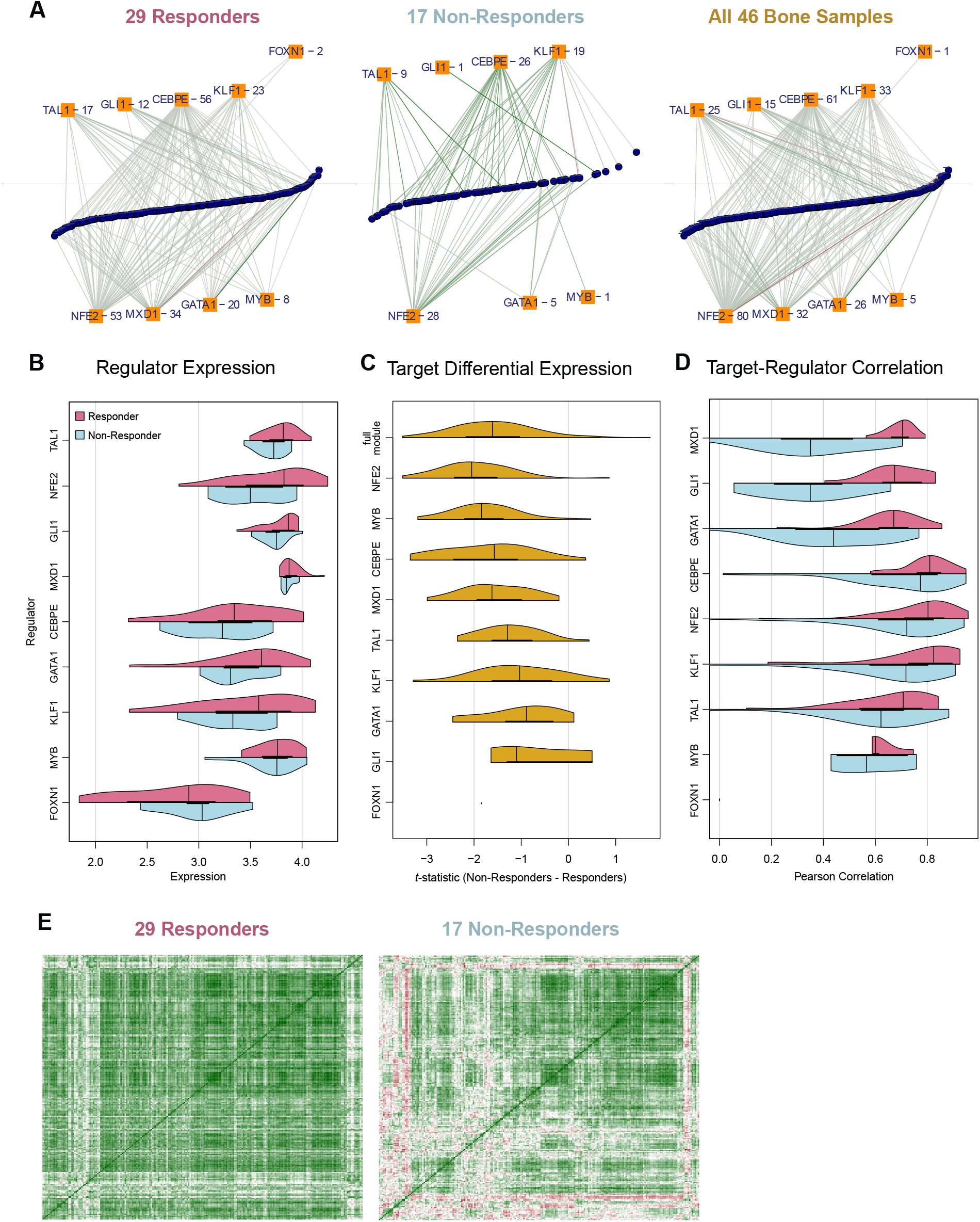
Regulatory Signals of P-M1. **A:** Sparse VBSR regulatory networks for P-M1 (Hematopoietic and immune response) built from expression data from only responders, only non-responders, and all bone samples. Regulators (orange squares) are connected to their target genes (blue circles) by edges colored by their regulatory relationships (green positive, red negative). **B:** For each regulator of the regulatory module community, a violin plot showing the regulator expression among responders (pink) and non-responders (blue). **C:** Changes in the expression (t-statistic) of the global all-bone sample targets of each regulator between responders and non-responders. **D:** Correlation relationships between regulators and their all bone sample targets among samples of the different response classes. **E:** Pairwise correlation among P-M1 genes within responders and non-responders.

Interestingly, the expression levels of the nine driving TFs were, generally, higher and less variable in R than NR (Figure 4B). Our method was also able to identify important module regulators that themselves were not differentially expressed (GATA1, KLF1, MYB) between the two patient groups. TAL1, a BHLH DNA binding transcription factor essential for maintaining the mutlipotency and quiencese of hemopoietic stem cells^51^ and implicated in the development of hemopoietic malignancies, was significantly differentially expressed (p-value 0.036) with decreased expression levels in NR. When considering the target genes of the module, we found regulators such as NFE2 and CEBPE, each regulated expression of more than 20% of the 222 targets in the bone samples, while others like MYB and FOXN1 only associated with the expression of a few targets. We found that across all regulators, the targets of those regulators were more highly expressed in R (Figure 4C), with NFE2 and CEBPE again regulating targets that were most strongly differentially expressed.

When we turned our attention to the types of relationships between regulators and their targets in P-M1, we observed that there was generally a positive correlation representing an activatory role of the regulators. In the patients who respond to abiraterone treatment, the positive correlations between the regulators and targets were much stronger (Figure 4D). For the 25 targets of TAL1 and the 33 targets of KLF1, the strong positive correlations were mostly preserved in R, while for other regulators like MXD1 (32 targets) and GLI1 (15 targets), their activation of the targets in NR seemed to be largely lost. The loss of the regulatory cohesion in NR became even more pronounced in the context of correlations between all genes in P-M1 (Figure 4E). A large percentage of gene pairs tightly co-expressed in the R showed weak or negative correlations in NR represented by three-month response phenotype.

In summary, TraRe identified important regulators of P-M1 that played distinct roles in phenotype-specific network rewiring, with target genes in P-M1 appearing to be much more strongly regulated in patients that responded to abiraterone treatment and looser cohesion of the regulatory module was found in the non-responders.

### The role of ELK3, MYB and MXD1 in abiraterone response

Finally, we validated our findings based on identified regulons with significantly different regulatory program between Abi R and NR using the prostate cancer cell lines LNCaP and 22Rv1. In order to examine the potential differential-regulation between Abi-sensitive and Abi-resistant settings, we developed LNCaP and 22Rv1 Abi-resistant cell lines, namely LNCaP-AR and 22Rv1-AR (Figure 5B-C). We started by selecting five regulons (ELK3, MXD1, MYB, ZNF3 and ZNF91) based on the following criteria: i) regulon significantly rewired between responder and non-responder (combined Fisher p-value<0.01); ii) the TF genes were expressed in both LNCaP and 22Rv1 using DepMap database (https://depmap.org/) (logTPM>1); and iii) TF regulated a series of highly repeatable targets for at least one of the response groups (Figure 5A, see supplementary S4 to S8 on pages 9–11) which were also expressed based on DepMap database (logTPM>1). All except ZNF91 were highly coexpressed with their targets in R only (Figure 5A). We performed siRNA knockdown of each TF (Figure 5D), and examined the top-ranked predicted targets based on multiplicity (Figure 5E-F). We found that targets in three regulons, regulated by ELK3, MXD1 and MYB, exhibited significantly differential regulation of downstream targets between Abi sensitive parental cells and Abi-resistant cells. Specifically, among the 30 tested targets in these three regulons, 19 were significantly decreased in at least one parental cell lines but not in the Abi-resistant line, 10 were significantly decreased in both parental cell lines but not in either Abi-resistant line after siRNA knockdown (Figure 5E-F). Therefore, we further tested whether these three regulons might differentially affect the proliferation in parental vs. Abi-resistant prostate cancer cell lines. Intriguingly, we found that knockdown of ELK3, MXD1 and MYB appeared to negatively affect the proliferation in 22Rv1 and LNCaP parental cell lines but had minimal or no impact on the Abi-resistant cells. These results indicate that the parental cell lines depend more on these TFs to survive.

**Figure 5:**
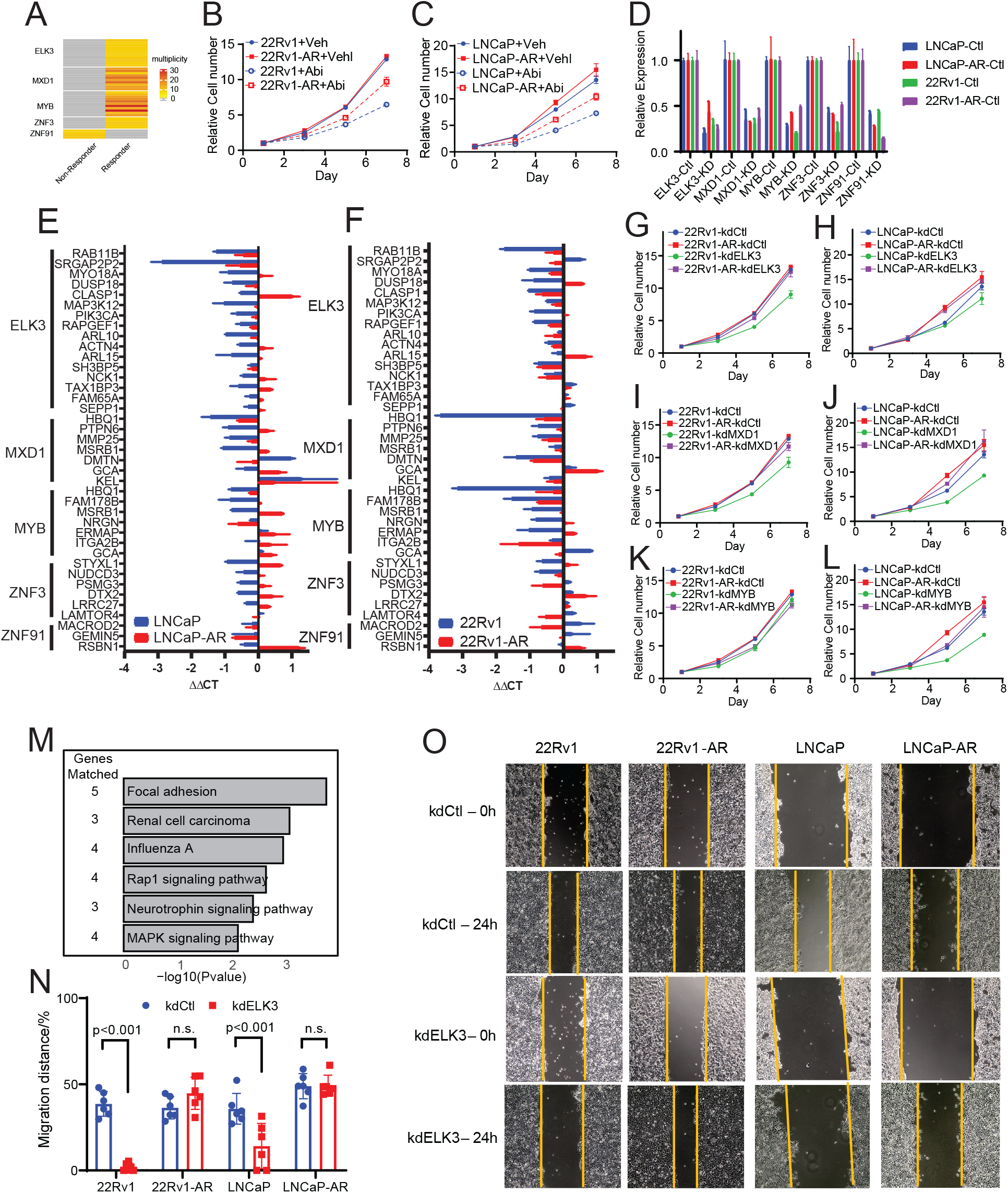
Experimental validation of regulons significantly different between responders and non-responders. **A:** Multiplicity of target genes identified within five selected regulons. **B-C:** Proliferation assay of 22Rv1 (**B**) and LNCaP (**C**): parental and Abi-resistant cell lines in the presence of vehicle or abiraterone, presented as mean *±* SD of 3 replicates. **D:** qRT-PCR measurement of knockdown efficiency of selected transcription factors, as mean *±* SD of 2 replicates. **E-F:** Quantification of expression of targets after knockdown of selected transcription factors using LNCaP (**E**) and 22Rv1 (**F**) and their respective Abi-resistant cell lines by qRT-PCR, as mean *±* SD of 2 replicates. **G-L:** Proliferation assay of 22Rv1 and 22Rv1 Abi-Resistant cell lines after knockdown of ELK3 (**G**), MXD1 (**I**) and MYB (**K**), presented as mean *±* SD of 3 replicates. Proliferation assay of LNCaP and LNCaP Abi-Resistant cell lines after knockdown of ELK3 (**H**), MXD1 (**J**) and MYB (**L**). **M:** KEGG Pathway analysis of ELK3 targeted genes. **N:** Representative picture of wound-healing assay of 22Rv1, LNCaP and their Abi-resistant cell lines after knockdown of ELK3. Pictures were taken at 0 and 24 hours after scratch. **O:** Quantification of wound healing assay. Data was presented as mean *±* SD of 6 replicates. Statistical difference was tested by paired two-sided Student’s *t*-test.

We also performed pathway analysis using the KEGG database to examine downstream functionalities affected by these TFs. We found that the targets of ELK3 were enriched in focal adhesion and Rap1 signaling pathway, which suggested a potential role of ELK3 in cell migration (Figure 5M). We thus performed wound-healing assay in ELK3 or non-targeting control siRNA transfected cells. We found that the migration was significantly inhibited in 22Rv1 and LNCaP parental cell lines, but not in the corresponding Abi-resistant cell lines when ELK3 was suppressed (Figure 5N-O).

In summary, we tested the regulatory effects of key TFs identified by TraRe on inferred targets in two models of Abi naïve and resistant cell lines obtaining differential response and cell proliferation, thus confirming our *in silico* findings.

### TraRe performance on simulated data benchmarks

In order to assess the performance of TraRe, we generated simulated data with 10 simulated regulatory modules. These simulated modules were created from randomly sampled regulatory programs (set of TFs driving the module’s expression) from the PROMOTE data, and adding multivariate Gaussian noise to generate expression patterns of target genes (Figure 6A, see Methods).

**Figure 6:**
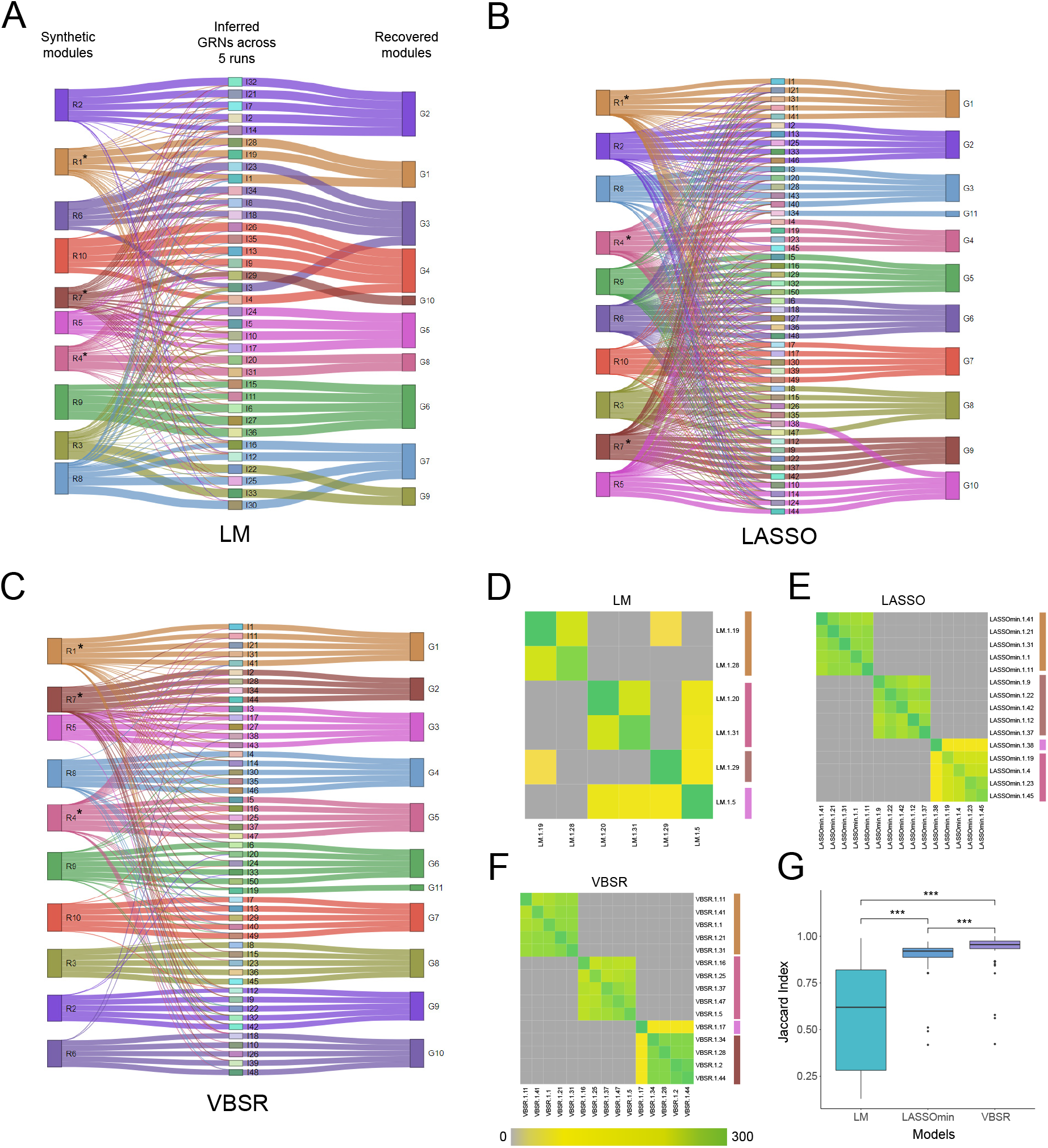
TraRe is able to recapitulate rewired regulatory modules on simulated data. **A-C:** Alluvial plots showing the flow of simulated regulatory modules to the corresponding inferred submodules and the cluster of these into regulatory modules by similarity for VBSR, Linear and LASSO models. **D-F:** Heatmaps showing clustered rewired regulatory modules from labeled-as-rewired simulated inferred submodules (simulated module 1, 4 and 7). **G:** Models boxplot and paired Wilcoxon test showing the distribution differences significance from the top 50 jaccard index scores.

Three well-known regression models were evaluated for their ability to discover the underlying simulated modules: i) Variational Bayes Spike Regression (VBSR)^13, 22, 24^, which is the one used by TraRe (see methods); ii) Least Absolute Shrinkage and Selection Operator (LASSO) regression, which has been used extensively to relate driver genes to their regulators^23, 30, 34^; and iii) Linear Regression Model (LM)^14, 52^, which is the baseline model. For each of these fitting models, TraRe was run 5 times using a random 80% of the samples to assess its generalization ability. To visualize the inference capabilities of TraRe for each model, we generated alluvial plots where the leftmost groups are the simulated regulatory modules, the middle column shows the inferred sub-modules across the different runs, and the column on the right shows the final inferred regulatory modules (Figure 6B-D). VBSR outperformed LM and LASSO, as TraRe was able to correctly infer and mark as rewired simulated sub-modules, and those were correctly clustered together (Figure 6B-D, Supplementary notes).

Further, to evaluate the capability of TraRe to infer rewired sub-modules under different regression models, three of the simulated regulatory modules were rewired by applying a whitening process to the gene expression data on half of the samples (see Methods). Therefore, the rewiring test was applied to the inferred sub-modules using a significance threshold of 0.01 (see Methods). The inference of rewired sub-modules was visualized via a heatmap, where a blockwise-diagonal heatmap is expected when the rewired mocules are properly discovered across runs (Figures 6E-G). From the three tested models, VBSR presented the largest similarity, as measured by the Jaccard index, to the simulated regulatory modules (Figure 6B, Supplementary notes).

Finally, we evaluated the behavior of VBSR, LASSO and LM over different noise levels, driver-assigning thresholds and number of samples. For each of them, we plotted the false positive rate (FPR) to true positive rate (TPR) curve. VBSR reported the largest TPR/FPR ratios on each of the scenarios (Supplementary notes and supplementary figures S3 to S6 on pages 17–20). Based on these findings, VBSR was selected as the default model for TraRe.

## DISCUSSION

We presented TraRe, a computational method to understand mechanistically altered regulatory dynamics through differential network analysis. We then applied TraRe to the transcriptomic data from the PROMOTE study to mechanistically understand how the regulatory differences in transcription networks may contribute to abiraterone response in mCRPC patients.

The analysis of differential networks can lead to a deeper understanding of network rewiring, elucidating molecular relationships associated with a characteristic of interest, such as disease progression or clinical treatments^53, 54^. Most of the approaches for differential network analysis have relied on different correlation-based metrics to measure the dependencies between pairs of gene nodes in the network^55, 56^. However, these methods are limited to marginal correlation networks (i.e., two nodes at a time) that are estimated separately using observations within each response group and generally do not consider relationships that are conserved across multiple groups. To address this issue, some methods separate group-specific conditional dependencies into global and group-specific components which have been shown to improve performance over other existing methods^57^. However, these methods were proposed for relatively small datasets (hundreds of genes at a time rather than thousands) and do not scale well to bigger datasets^58^. In addition, they do not consider the overall rewiring of individual GRNs and instead focus on pairwise gene-gene differential correlations.

The developed method TraRe, on the other hand, provides a robust and efficient methodology to perform differential network analysis, evidenced by handling large datasets of more than 20K genes. TraRe builds regulatory modules through an efficient module-based approach ^22^ and identifies key transcription factors via Variational Bayes Spike regression. To uncover potential transcriptional rewirings associated with a particular phenotype TraRe performs an efficient permutation test on the Frobenious norm of the difference of the estimated covariance matrices of each phenotype.

When applied to the PROMOTE mCRPC dataset, TraRe highlighted key regulatory modules associated with essential regulatory dynamics. These regulatory modules were also recapitulated when applying TraRe to the independent SU2C mCRPC study. More importantly, TraRe identified a regulatory module (P-M1) associated with immune response that was highly enriched in rewired regulons between Abiraterone R and NR. One of its TFs, TAL1, was strongly differentially expressed and showed an activating relationships with targets in both patient groups whereas the TF NFE2, although not significantly differentially expressed itself, led to strong differential expression of downstream target genes, demonstrating a mostly activating role. On the other hand, the TFs GLI1 and MXD1 were also not significantly differentially expressed but showed a significant loss of correlations with their targets in the NRs. These findings show the complexity of the regulatory landscape of metastatic cancer tissue and how TFs may have distinct roles in phenotype-specific network rewiring.

Leveraging TraRe’s inferred information, we identified and validated three transcription factors, ELK3, MYB and MXD1, whose expression were not significantly different between Abi R and NR, however, their regulatory networks were significantly disrupted in NR. The regulons driven by these TFs in R probably played important roles in tumorigenesis or progression in the Abi-sensitive samples. For example, we showed that knockdown of ETS family gene ELK3 suppressed the proliferation of Abi-sensitive prostate cancer cell lines and, more importantly, inhibited migration by suppressing the expression of genes related to focal adhesion and motility, including RAPGEF1^59^, ACTN4^60^, RAB11B^61^, MYO18A^62^, and PIK3CA^63^.

These genes have been previously linked to RAC1-PAK1 mediated E-cadherin stability, epithelial-mesenchymal-transition, FAK-induced invasion and metalloproteinase expression. This finding is consistent with Mao *et al.*^64^, who showed that silencing of ELK3 in prostate cancer cell lines induced S-M phase arrest, inhibited cell proliferation and migration. However, these regulations were not observed in Abi-resistant settings, probably due to the extensive reprogramming of regulation networks in the development of Abi-resistant phenotype. This may help explain why ETS family and ETS-fusion, despite their high prevalence in prostate cancer^65, 66^, are controversial in their prognostic values, especially in metastatic setting^67^.

Similarly, we found that knockdown of either MYB and MXD1 suppressed proliferation of Abi-sensitive cell lines. MYB is known to be overexpressed in prostate cancer cells. It interacts with AR and sustain AR activity under androgen-depleted condition^68^. MXD1 usually functions as antagonist of MYC-MAX signaling, but it was also reported to mediate HIF-1 *α*-induced PI3KAKT activation and chemoresistance in U2OS and MG-63 cells^69^. Interestingly, we found that MYB and MXD1 shared a number of genes in their regulons, such as DMTN and MSRB1, the prior of which regulates actin cytoskeletal organization^70^, while the latter responds to oxidative stress^71^. This crosstalk of regulons indicated potential co-regulations between the two transcription factors, which was not widely reported previously. Again, these signaling cascades have been disrupted in the development of Abi-resistance, probably due to the reprogramming of the transcriptional regulation networks. All these results emphasized the difficulties to therapeutically target transcription factors due to their dynamic regulatory adaption.

## AVAILABILITY

RNAseq data from PROMOTE and SU2C studies are available at the database of Genotypes and Phenotypes (dbGaP) with accession numbers phs001141.v1.p1 and phs000915.v2.p2, respectively. TraRe package is available at Bioconductor^72^ and GitHub repository ubioinformat/TraRe. Matrices and scripts used for the analysis are available at the ubioinformat/TraRe-materials GitHub repository.

## SUPPLEMENTARY DATA

Supplementary Data are available at NAR online

## Supplementary Material for

### S1 Supplementary Notes

#### S1.1 Model and Sample Selection Comparison

The most robust rewired regulatory module in the PROMOTE data, PM-1, was found using the 46 bone-only samples and Variational Bayes Spike Regression (VBSR) models to construct the regulatory submodules. We also investigated the effect on the results of selecting that particular regression model and the single metastatic site. For this analysis, we normalized 67 PROMOTE patient samples from all metastatic sites (41 responders and 26 non-responders) with conditional quantile normalization. We kept these expression values for only genes which had at least 32 reads from at least 2% of the samples from each class. For each configuration, submodules discovery was completed with 80% of the samples in 10 repeated runs. Submodules across runs were scored for rewiring, and rewired submodules were clustered using hierarchical clustering (Euclidean distance and ward linkage) on the similarity matrix of submodule log_10_ hypergeometric test *p*-values. This process for finding robust, rewired modules was run for three different types of regression models (VBSR, Linear (LM), and LASSO (LASSOmin)) and two different subsets of patient samples (41 bone-only and 67 all sample sites). For each of these 6 configurations, VBSR was used to find the regulatory network edges of the module formed from the cluster with the largest number of rewired submodules. In spite of the differences between these analyses and with the analysis in the main text, each configuration resulted in similar regulatory networks with a similar set of core regulators, CEBPE, GATA1, IKZF1, KLF1, NFE2, and MXD1 (see Supplementary Table S12). The most interesting variation between the resultant networks comes from the number of total driver regulators that contribute to the expression of the discovered submodules. We ultimately selected VBSR for submodule discovery as presented in the main results because it consistently returned modules with strong rewiring significance using a minimum number of driver regulators.

#### S1.2 Simulated data

##### S1.2.1 Alluvial plots analysis

When using VBSR as the fitting model for the module inference process, even though most of the inferred modules across runs were composed by a mixture of genes coming from different simulated modules (Figure 6B-middle column), the majority of the genes came from a unique simulated module while few came from another simulated module. More importantly, inferred modules were correctly grouped into a single module after the clustering step that integrates different runs (Figure 6B-leftmost column). Moreover, rewired modules were correctly inferred and labeled as rewired by TraRe, where rewired modules inferred across different runs were correctly clustered together (Figure 6C).

When LASSO was selected as the fitting model, the inferred GRNs contained a larger presence of drivers from other modules than when using VBSR which yielded a decrease on performance with respect to VBSR (Figure 6D). Nonetheless, rewired simulated modules were correctly identified and grouped together across runs (Figure 6F). Note that the similarity scores between rewired modules of different runs was generally lower than when using VBSR (Figure 6F).

LM, on the other hand, often inferred noisy modules containing large mixed proportions of genes coming from several simulated modules (Figure 6D). This behavior led to a single inferred GRN to be identified as rewired. Further, similarity between rewired GRNs belonging to different simulated modules emerged due to the mixed proportions of some inferred GRNs (Figure 6G).

##### S1.2.2 Parameter grid analysis

We assessed the behavior of the different models to changes on the noise level when generating the simulated modules. In this case, LM was only able to infer modules in a short range of noise (*σ* = [1.2, 1.6, 2]), as opposed to LASSO, which inferred modules on a larger range of noise variances at the expense of generally large false positive rates (FPR). VBSR, on the other hand, was the only model that ensured a high true positive rate (TPR) while controlling the FPR in a relatively large range of noise values (*σ* = 0.01 *−* 2.8), see Figure S4.

We then evaluated the different models by varying the internal threshold that each method uses to assign a driver gene to a target gene: the p-value cutoff in VBSR and LM, and the *λ* parameter in LASSO (see Methods). LASSO and LM obtained large FPRs even for small thresholds, while VBSR achieved the largest area under the receiver operating characteristic (ROC) curve (AUCs) on every Jaccard index threshold. (Figure S5).

We then verified the performance of the different models as a function of the number of available samples (i.e., patients) via ROC curves. VBSR achieved the largest AUCs above 28 samples. LASSO was the only model able to infer GRNs under 28 samples, maintaining similar TPR to FPR ratios of larger samples runs (Figure S6)

Finally, when measuring the TP, FP and FN for a fixed noise variance, number of samples and internal thresholds (see Methods), VBSR and LASSO models outperformed the LM in terms of the adjusted coefficient of determination (adjusted R-squared coefficient) and the recall score. On the other hand, LM and VBSR outperformed LASSO in precision (Figure S3).

Note that the evaluation metrics rely on matching inferred GRNs to the simulated ones. Thus, Jaccard indexes were used as thresholds to define at which extent we included or not a matched pair in the evaluation. Four different Jaccard thresholds were used: 0, where every match is included regardless of the number of overlapping genes (as long as these are larger than one); 0.5; 0.8; and max, where only one pair, the one with largest Jaccard index, is included for each simulated module.

### S2 Additional Tables

**Table S1:**
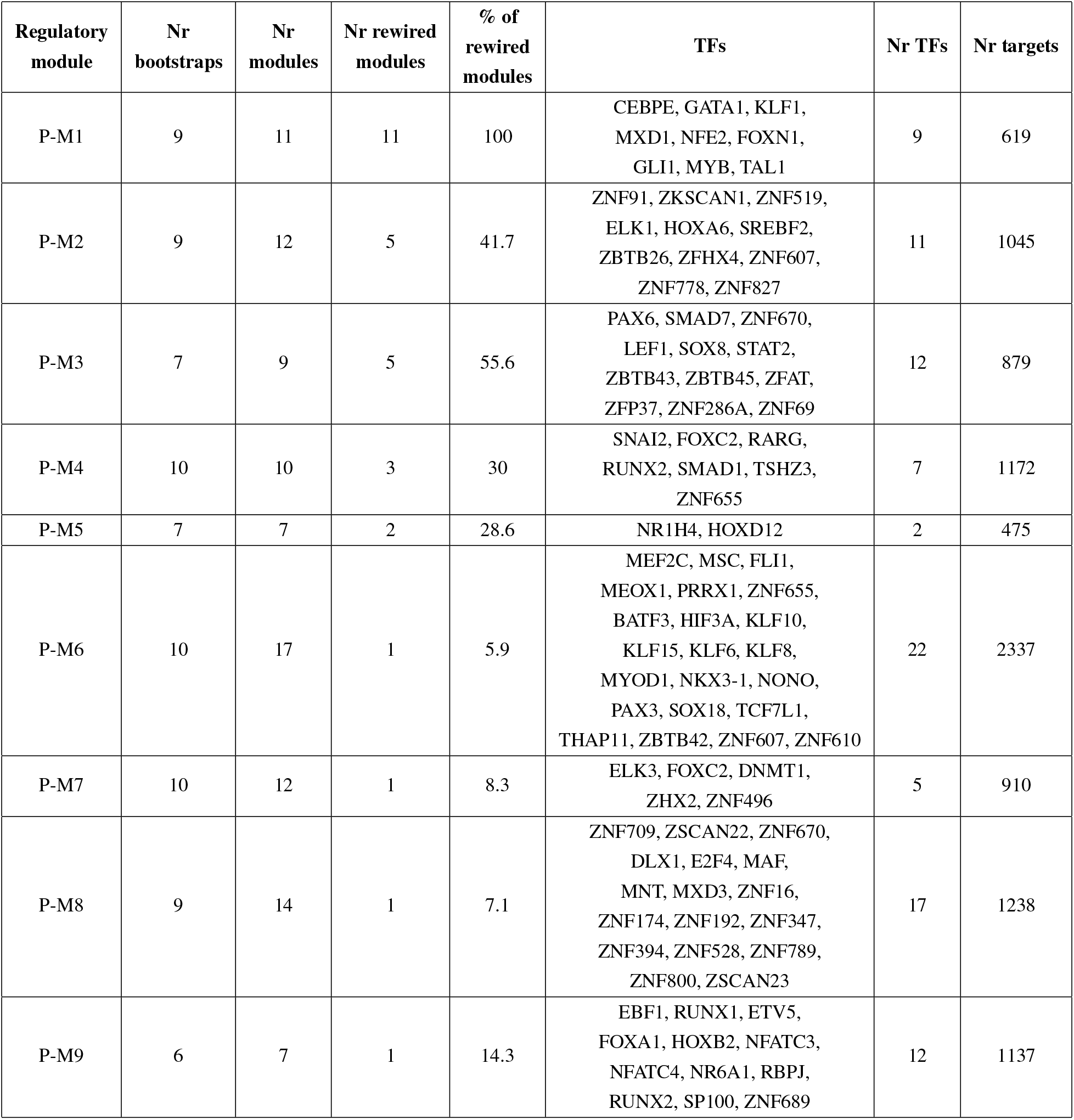
Summary table of regulatory modules generated with cAMARETTO software^30^ from all sub-modules obtained in a 10-bootstrap run with TraRe on PROMOTE dataset (Part 1: Rewired Regulatory Modules).

**Table S2:**
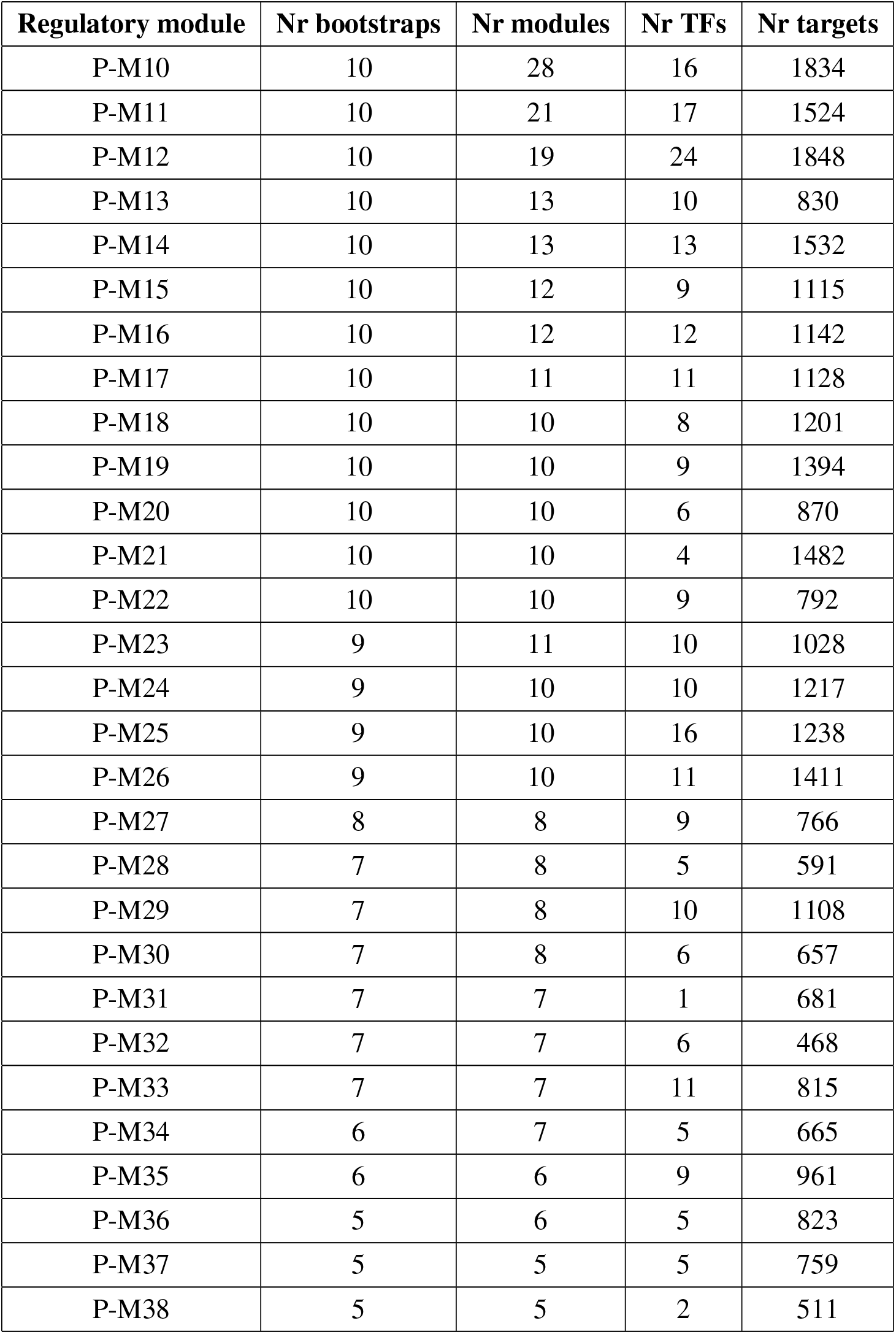
Summary table of regulatory modules generated with cAMARETTO software^30^ from all sub-modules obtained in a 10-bootstrap run with TraRe on PROMOTE dataset (Part 2: Non-rewired Regulatory Modules).

**Table S3:**
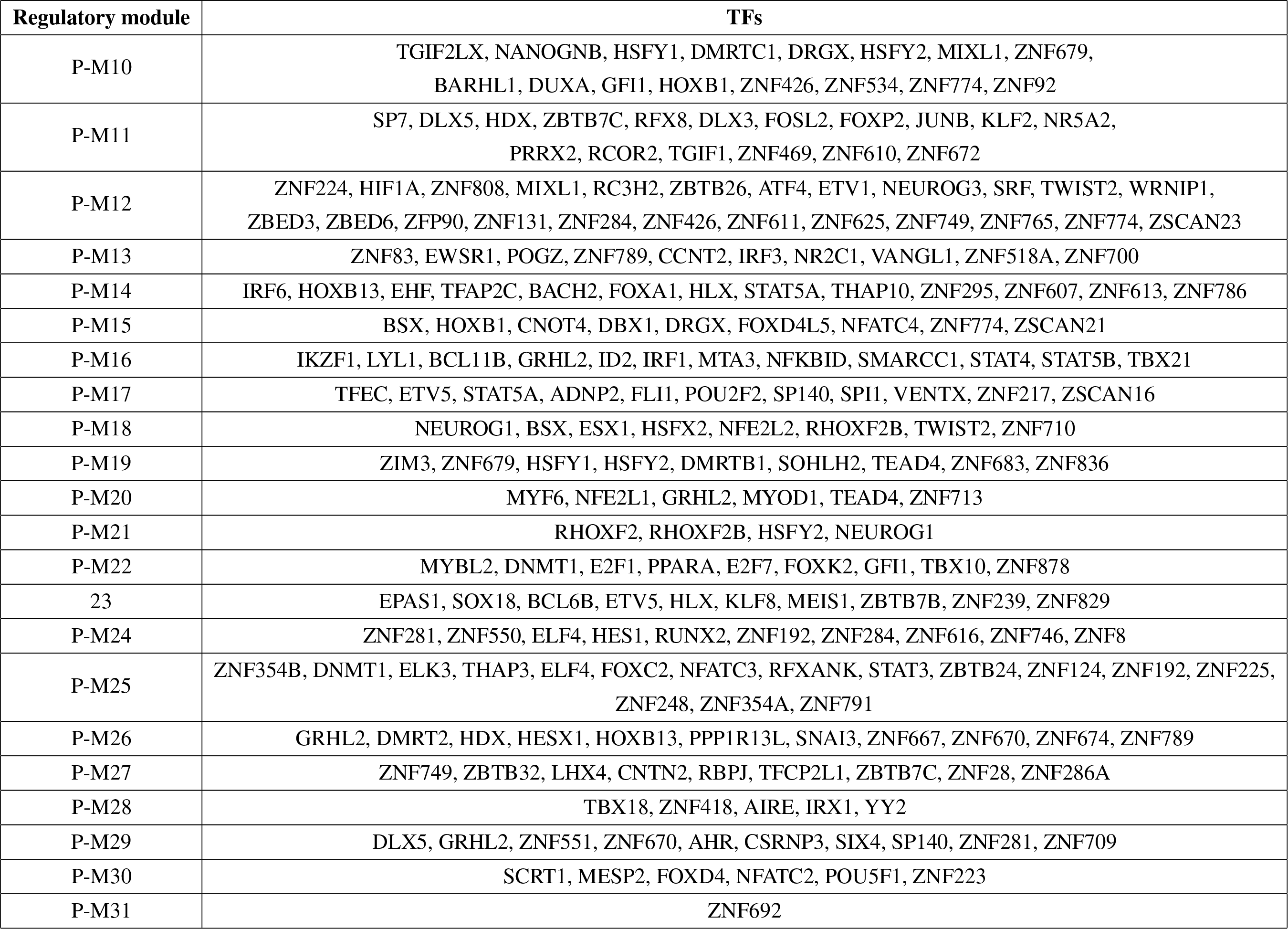
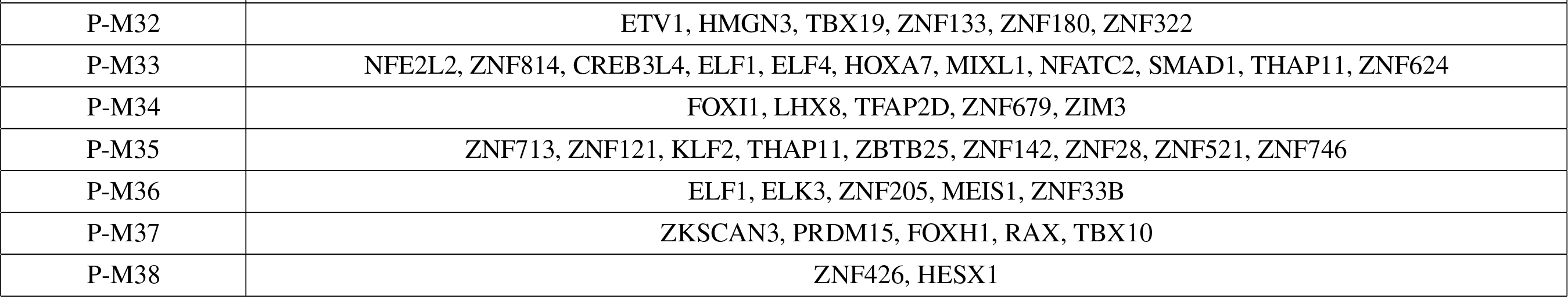
Summary table of regulatory modules generated with cAMARETTO software^30^ from all sub-modules obtained in a 10-bootstrap run with TraRe on PROMOTE dataset (Part 3: Non-rewired Regulatory Modules TF list).

**Table S4:**
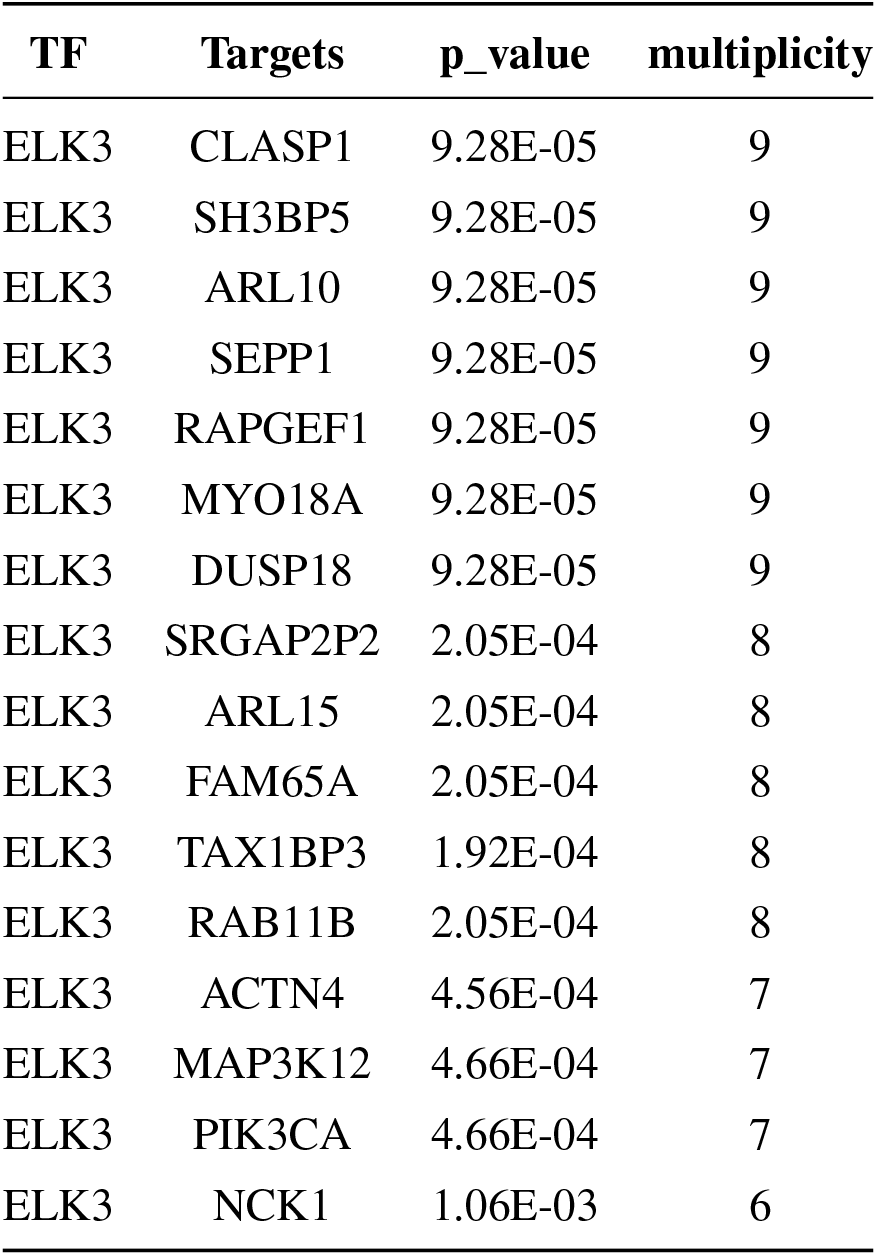
ELK3 regulon with selected targets for biological validation

**Table S5:**
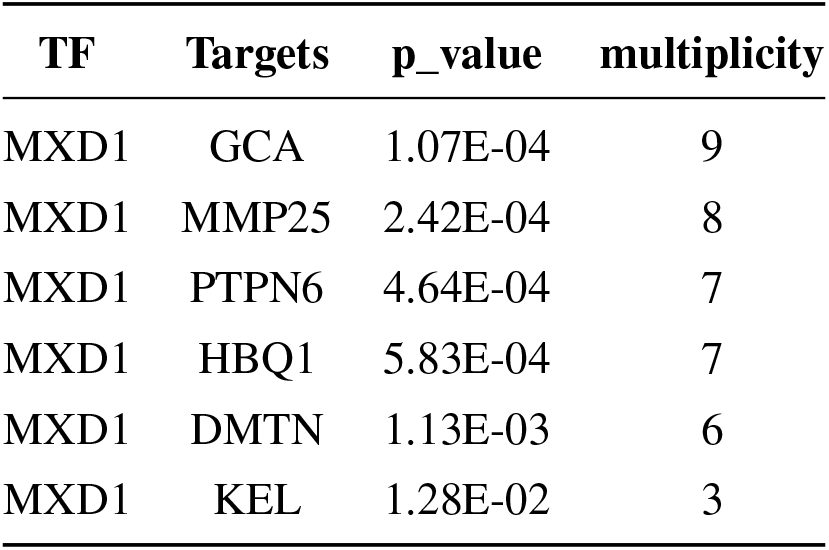
MXD1 regulon with selected targets for biological validation

**Table S6:**
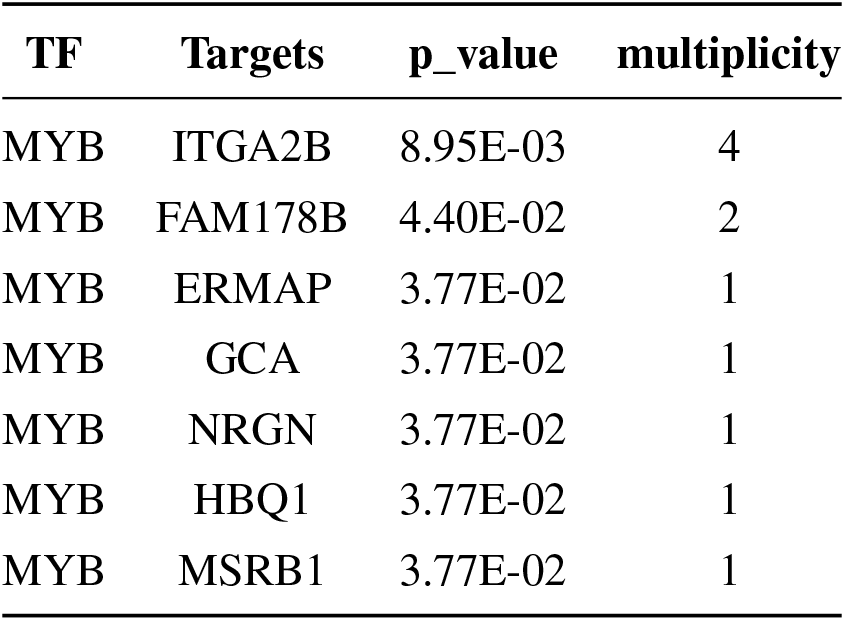
MYB regulon with selected targets for biological validation

**Table S7:**
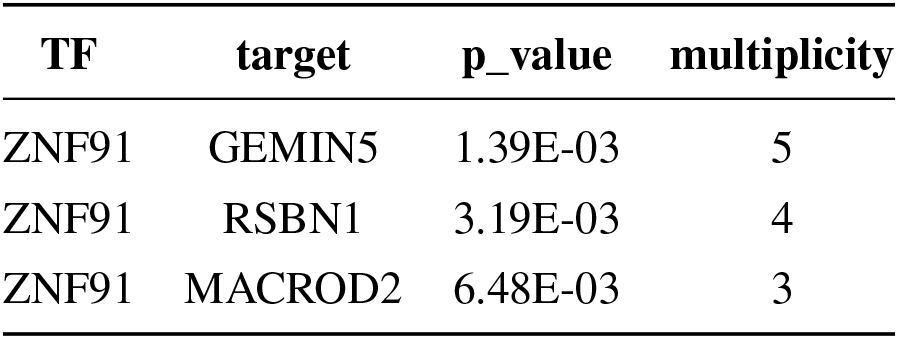
ZNF91 regulon with selected targets for biological validation

**Table S8:**
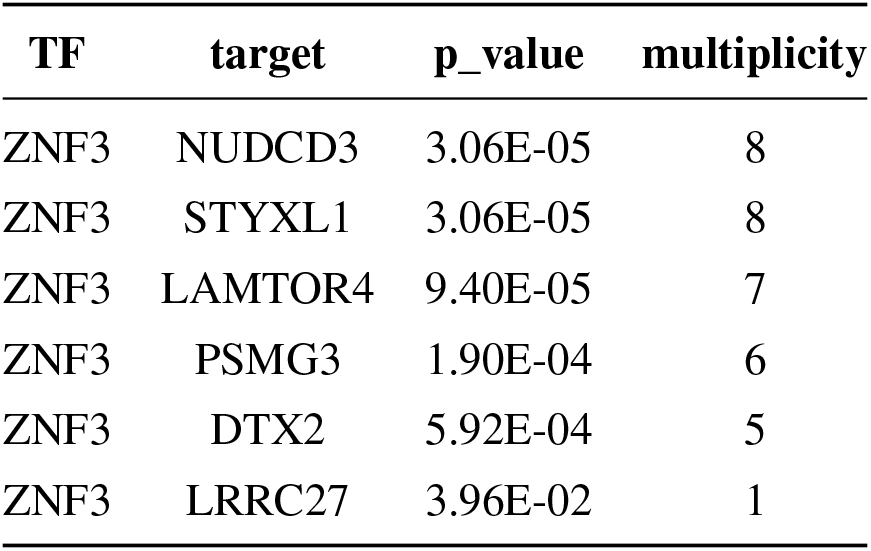
ZNF3 regulon with selected targets for biological validation

**Table S9:**
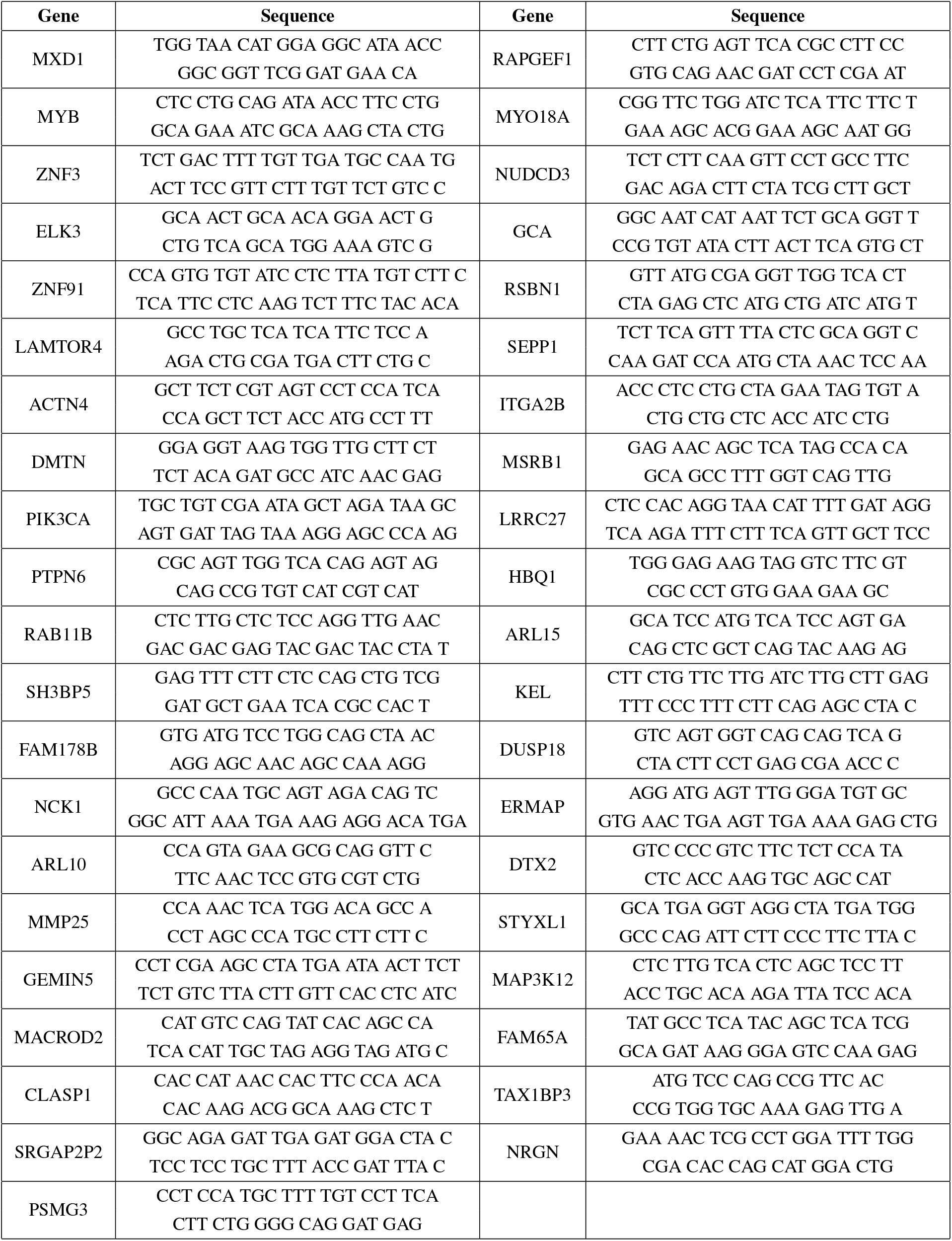
Primer sequences for qRT-PCR assays

**Table S10:**
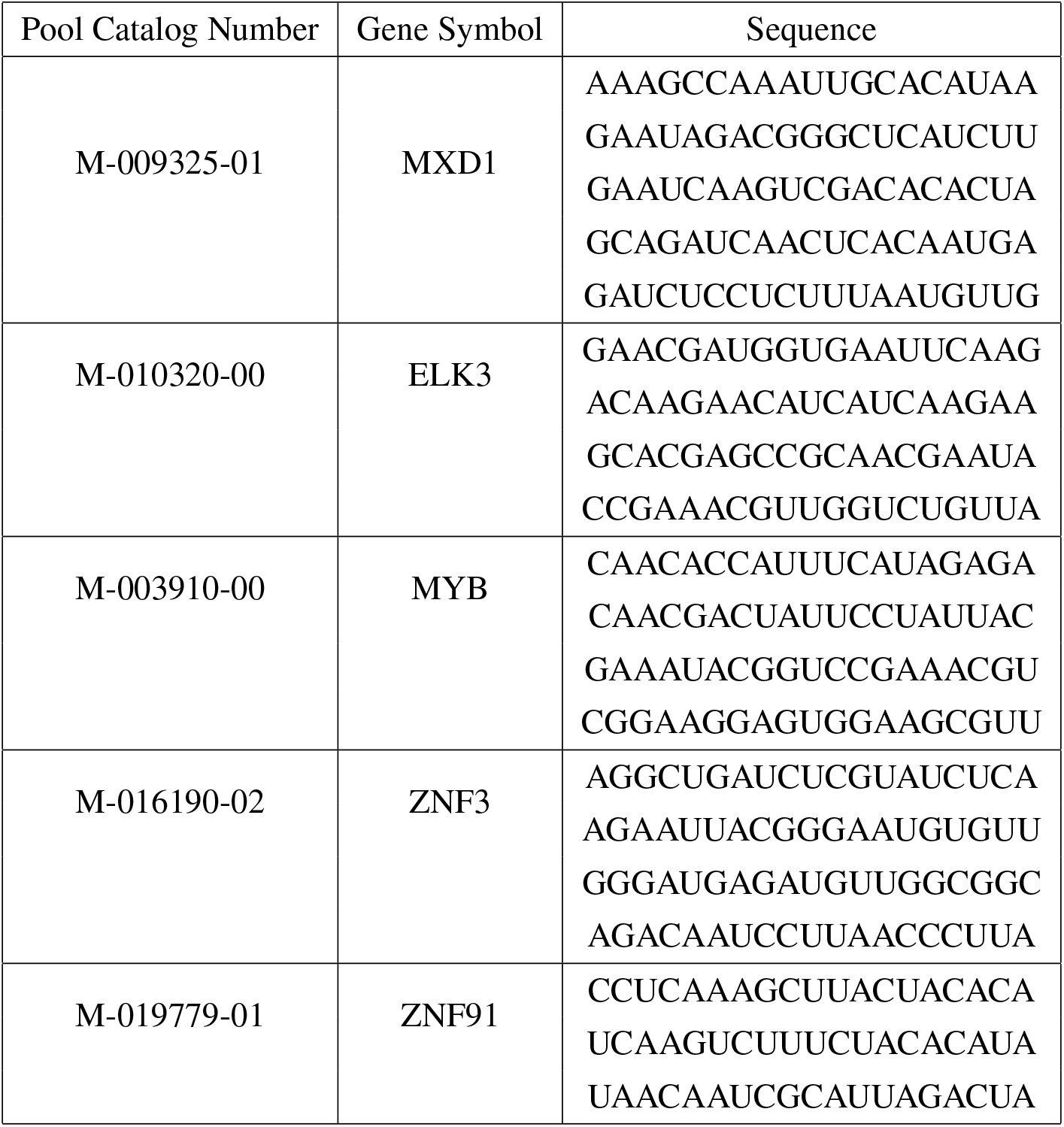
Sequences of siRNA for selected transcription factors.

**Table S11:** Enrichment of regulons with Gene Ontology Biological Processes. See Excel Supplementary Data.

**Table S12:**
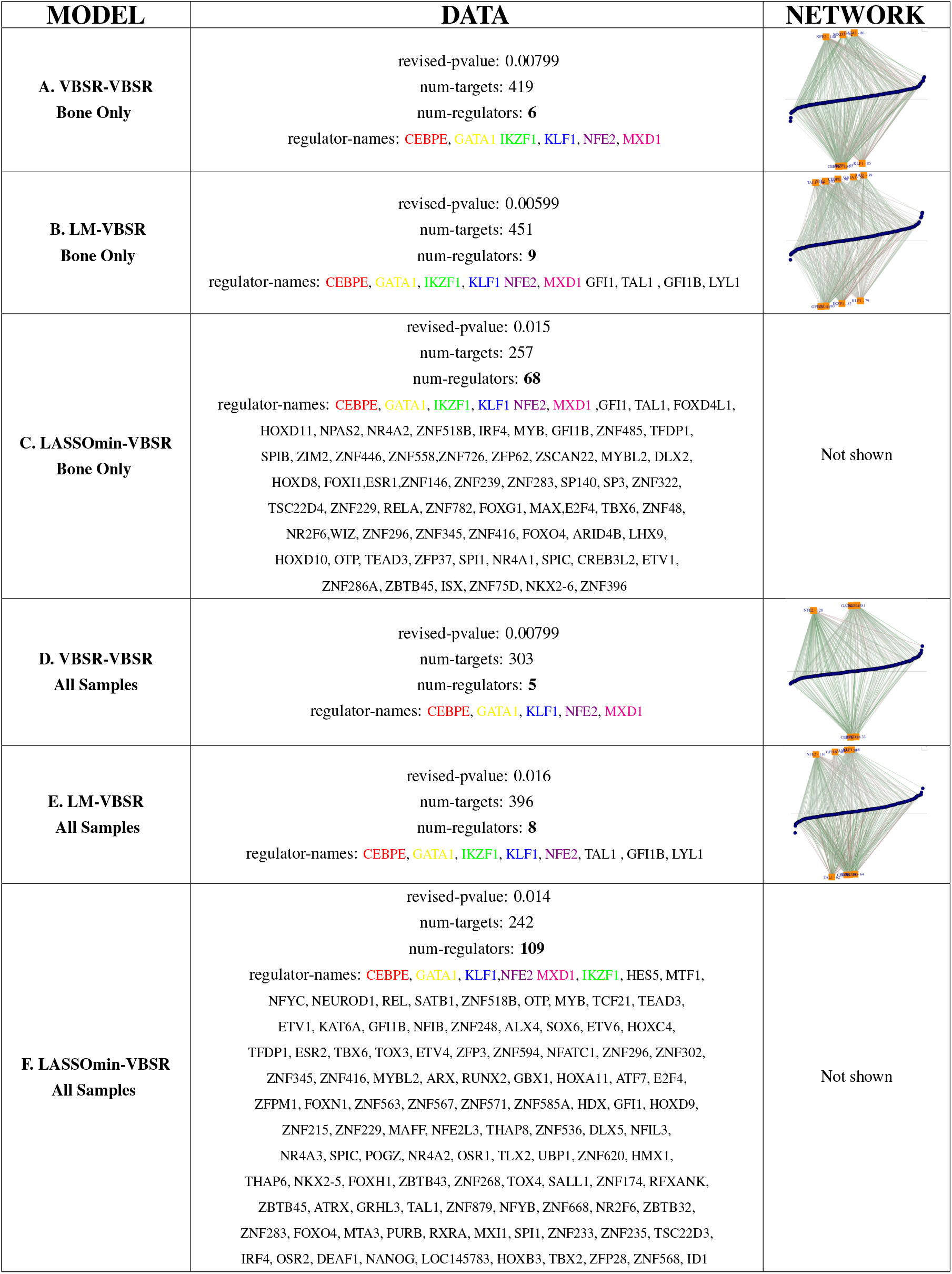
Consistent Discovery of P-M1. Identifying the most strongly rewired regulatory module from the 46 bone-only samples (**A-C**) resulted in similar regulatory networks, with the same core drivers (in colored text), regardless of the regression model used to construct the co-expressed modules. This finding also held for using the data from all 68 patient samples (**D-F**).

### S3 Additional Figures

**Figure S1:**
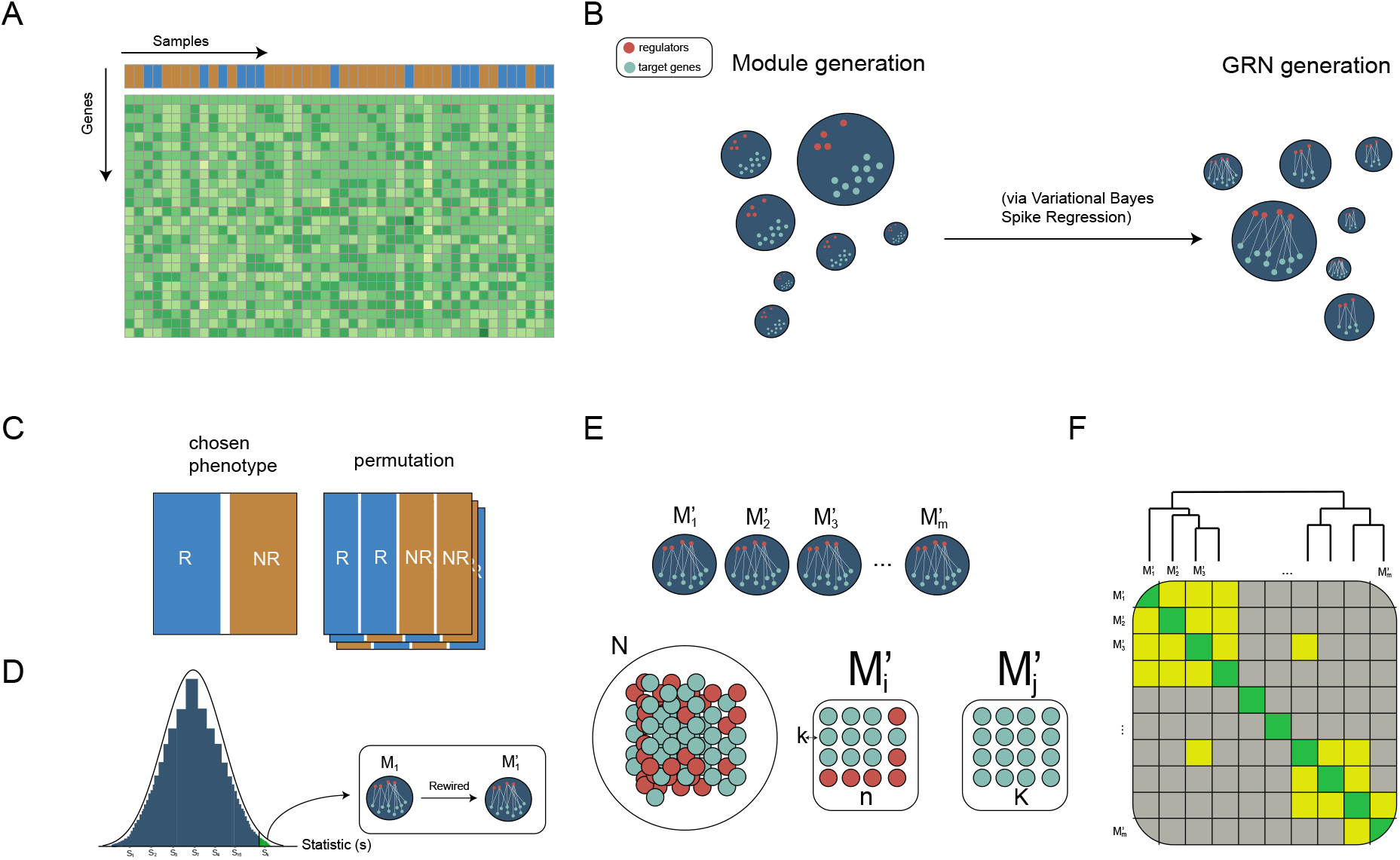
Workflow of TraRe. **A:** Example of a gene expression matrix (genes by samples). Color of samples indicates whether the patients are labeled as responders or non-responders. **B:** GRN inference process in two steps, module generation and GRN generation. **C:** GRN rewiring process. A permutation test on sample class labels is performed per module. **D:** A dissimilarity metric is evaluated for the true class labels and compared to a fixed amount of permutations. **E:** Robust rewired GRN inference process begins with Hypergeometric tests performed between rewired modules. **F:** Similar rewired modules are grouped via a hierarchical clustering yielding robust, regulatory modules.

**Figure S2:**
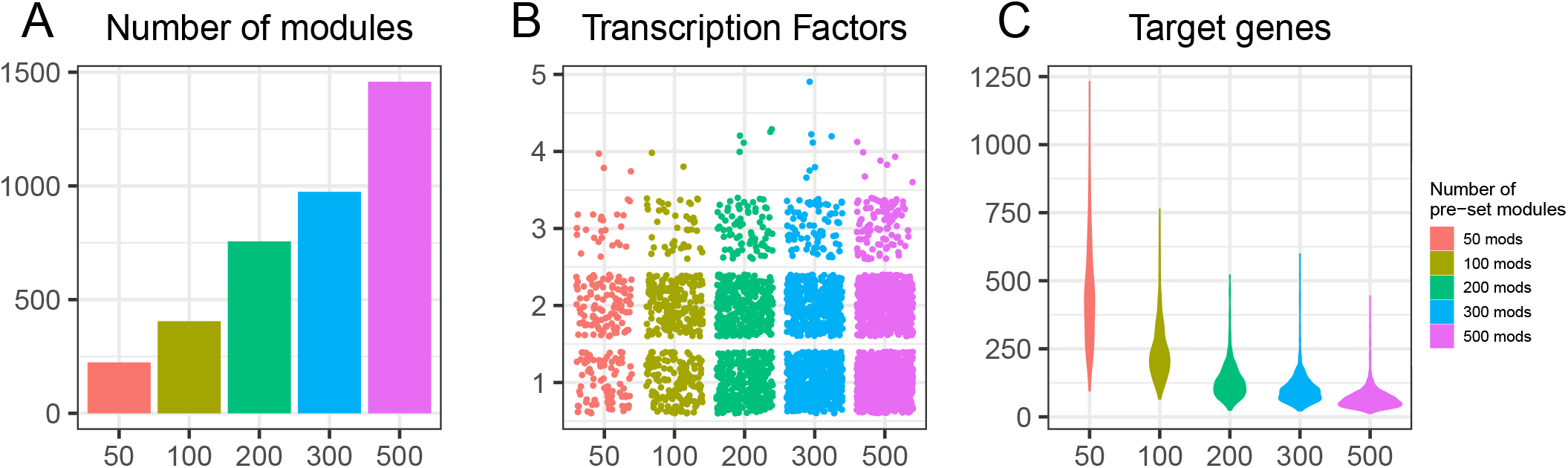
Variation of sub-module generation. We modified the number of the pre-set sub-modules to 50, 100, 200, 300 and 500 on PROMOTE data and run 5 bootstraps.Variation of sub-module generation. We modified the number of the pre-set sub-modules to 50,100,200,300 and 500 on PROMOTE data and run 5 bootstraps. We show the influence in the number of sub-modules generated (**A**), the number of transcription factors per generated sub-module (**B**) and the number of targets per generated sub-module (**C**).

**Figure S3:**
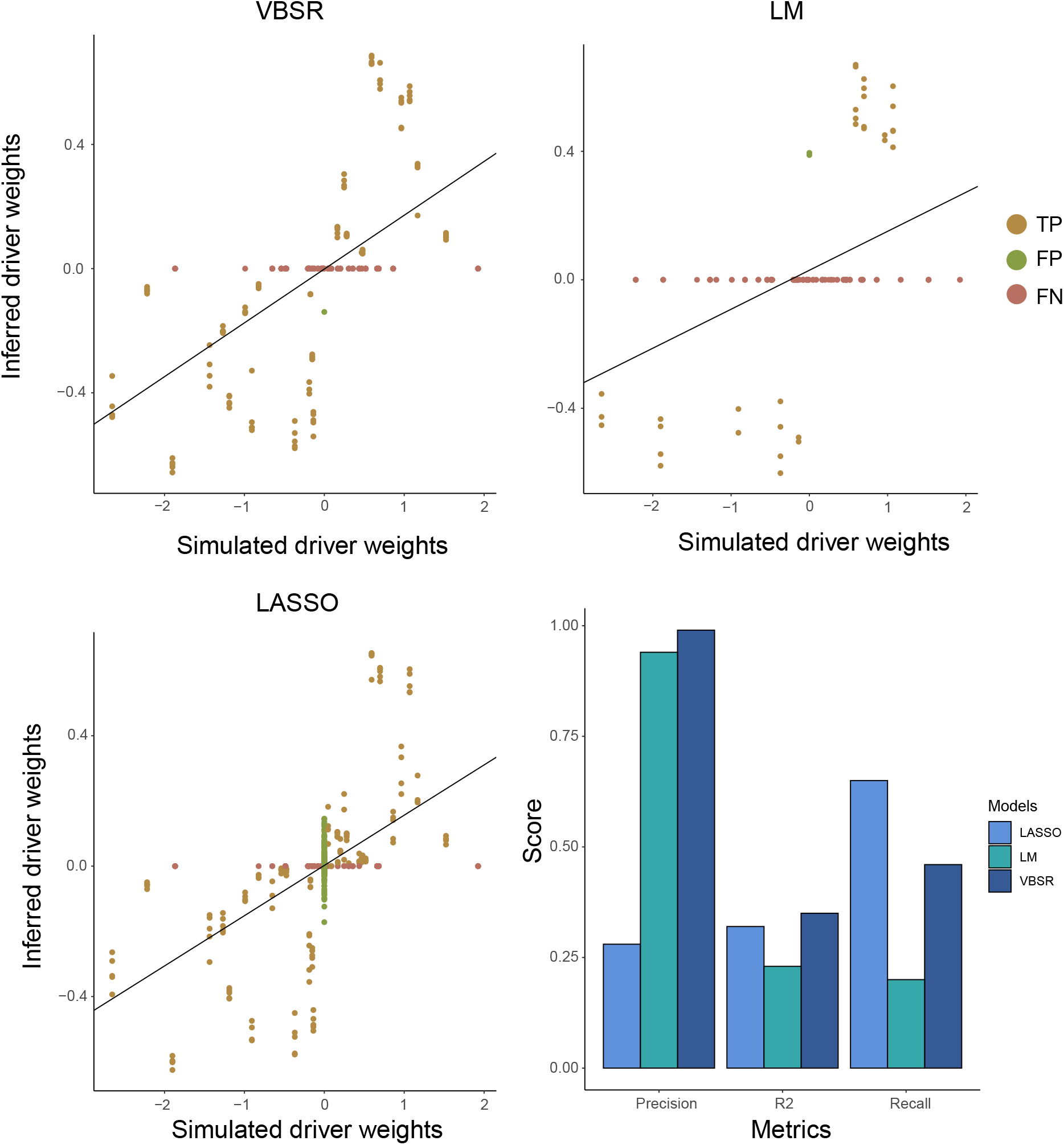
Classification metrics along VBSR,LM and LASSO. For a fixed Jaccard index, variance noise *σ*, number of samples and p-value thresholds, TP, FP and FN from simulated-inferred modules pairs are shown for each model. A linear model has been fitted to explain driver’s regulatory programs within inferred GRNs based on simulated regulatory programs. Moreover, *R*^2^ adjusted, precision and recall parameters from the fitted linear model are shown. VBSR obtained perfect precision, which was almost scored by LM, and also obtained the largest *R*^2^. Largest recall score was obtained by LASSO model, as it over-inferred drivers, which can be seen in the form of FPs at its precision score.

**Figure S4:**
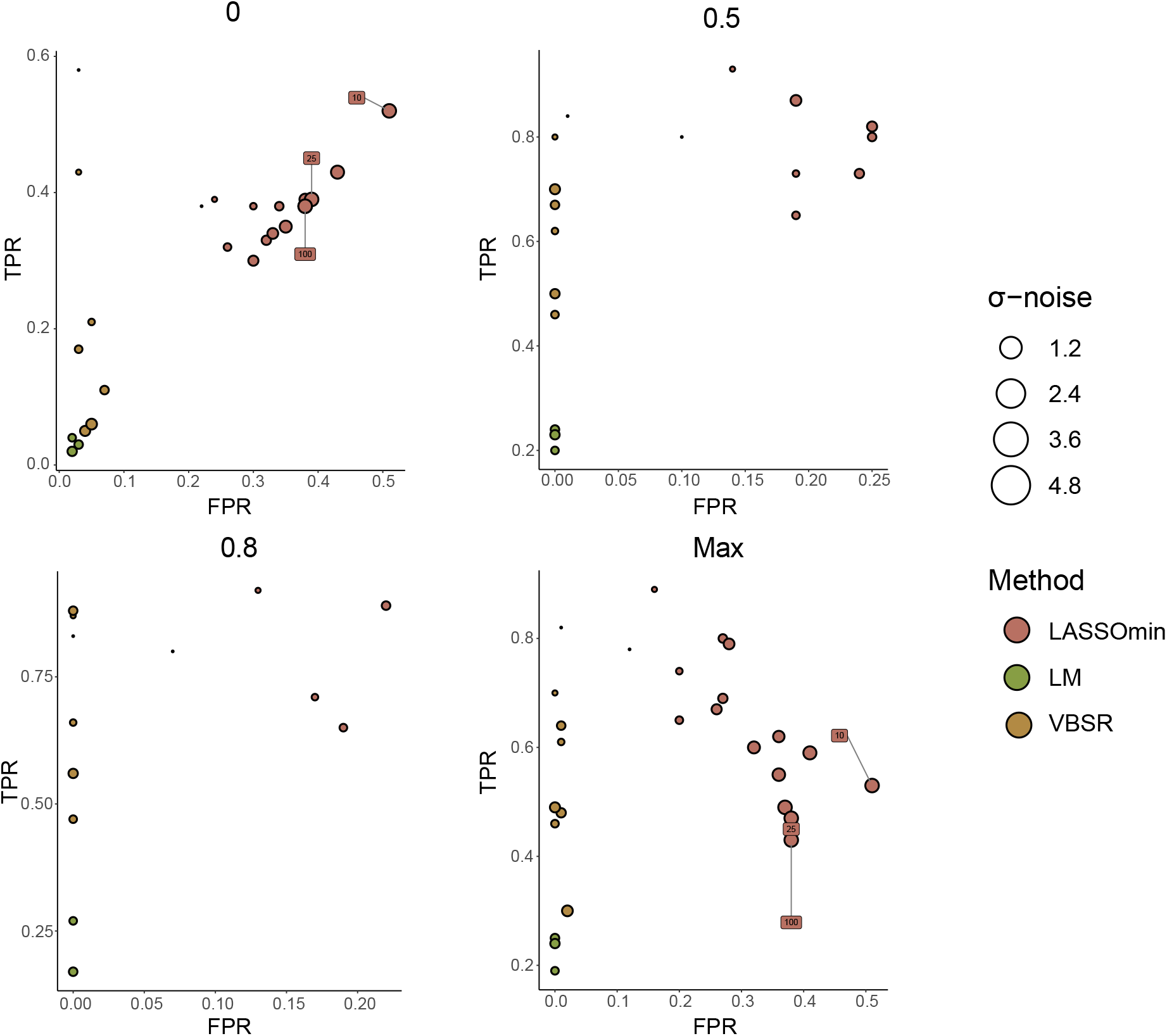
Discrete ROC’s curves comparing simulated and inferred modules as a function of the simulated modules variance noise *σ*. For each defined Jaccard index threshold and each value of *σ* = [0.01, 0.4, 0.8, 1.2, 1.6, 2, 2.4, 2.8, 3.2, 3.6, 4, 4.4, 4.8, 10, 25, 50, 100], TPR and FPR have been computed along VBSR, LASSO and LM. Default number of samples and p-value thresholds have been used. As the Jaccard index threshold decreases (more restrictive) the fitting models are less robust to noise variance. Also, VBSR achieved large TPR to FPR ratios in a wide range of noises (up to *σ*=2.8).

**Figure S5:**
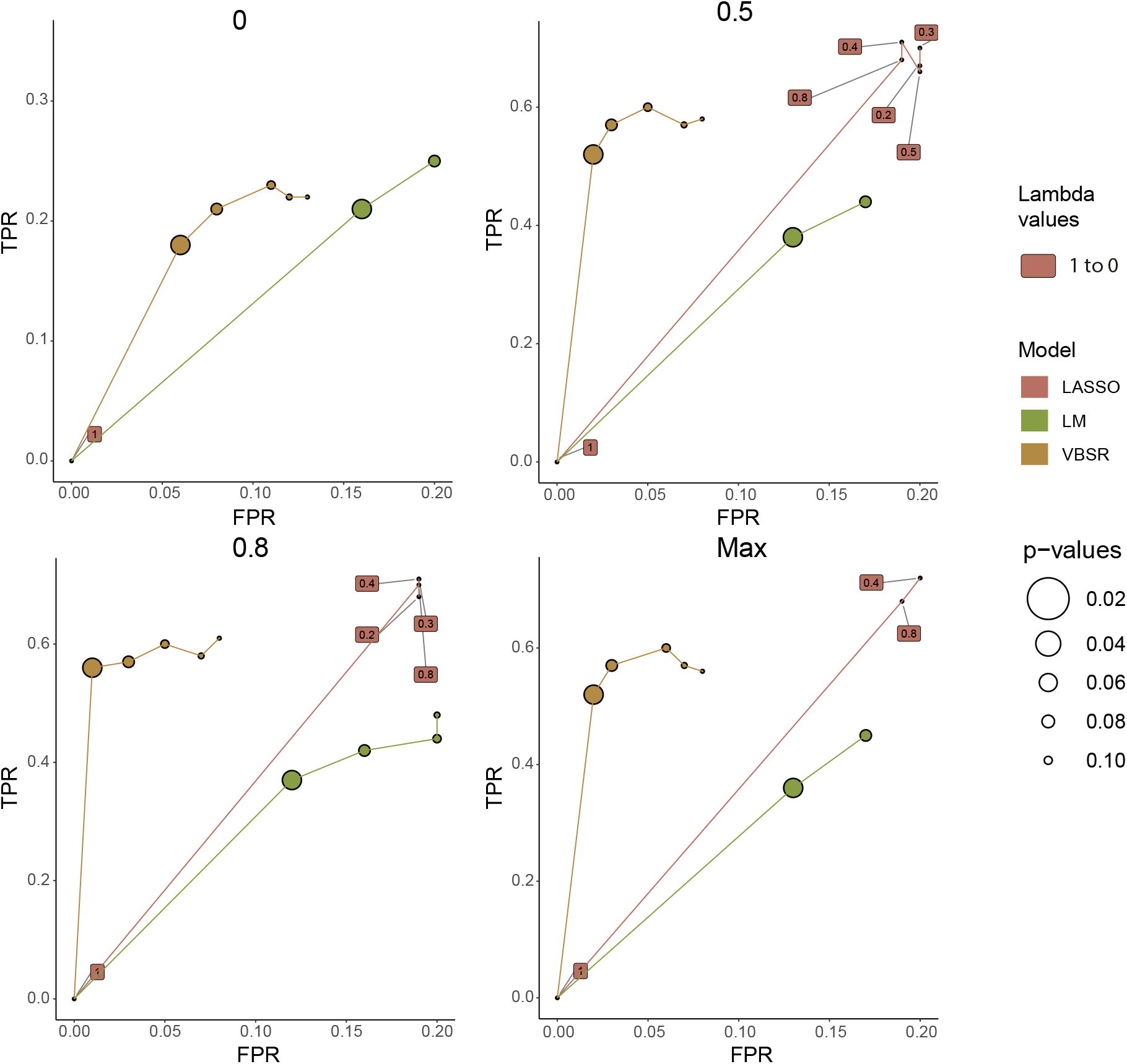
ROC’s curves as a function of p-value thresholds. TPR and FPR have been computed along a desired p-value’s range (0.02 to 0.14) for each Jaccard index threshold and fitting models VBSR, LASSO and LM. Noise *σ* and number of samples have been set to its default parameters. Note that, for the LASSO model, Lambda parameter has been modified as a metric of sensitivity, selecting Lambda’s deciles from LASSO cross validation model. (R’s *cv.glmnet* package). VBSR achieves the largest TPR to FPR ratio on every Jaccard index threshold.

**Figure S6:**
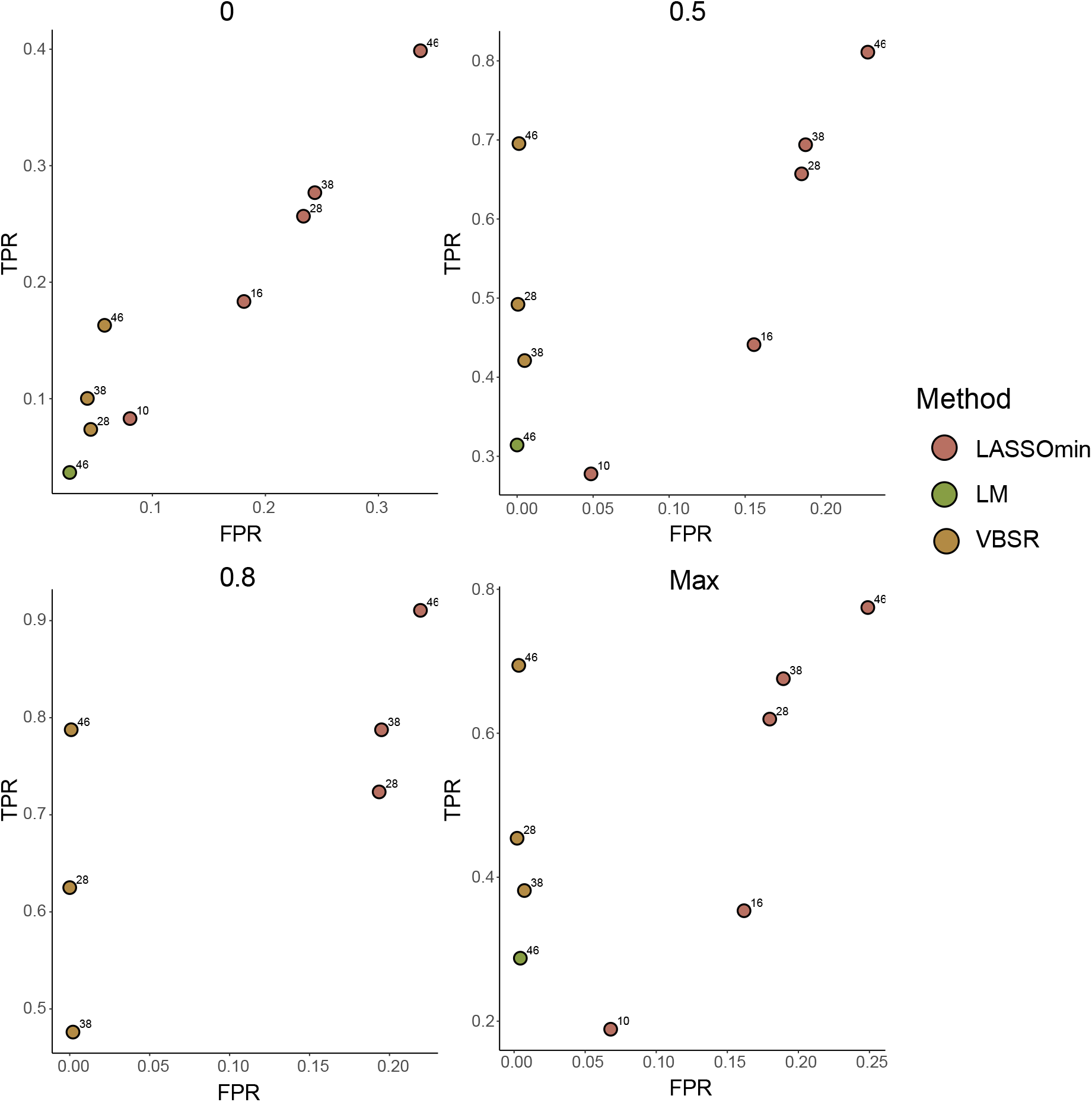
ROC’s curves a function of samples (patients). TPR and FPR have been computed for each Jaccard index threshold and fitting models VBSR, LM and LASSO, along 10, 16, 28, 38 and 46 samples. Noise *σ* and p-value’s threshold have been set to its default parameters (see Methods). Missing points when threshold = 0 are due to fitting errors from specific models. Missing points distinct to the aforementioned in the remaining thresholds are due to the filtering. Above 28 samples, VBSR achieved largest TPR to FPR ratio on every Jaccard index threshold. LASSO was the only model able to infer GRNs under 28 samples, maintaining similar TPR to FPR ratios of larger samples runs.

**Figure S7:**
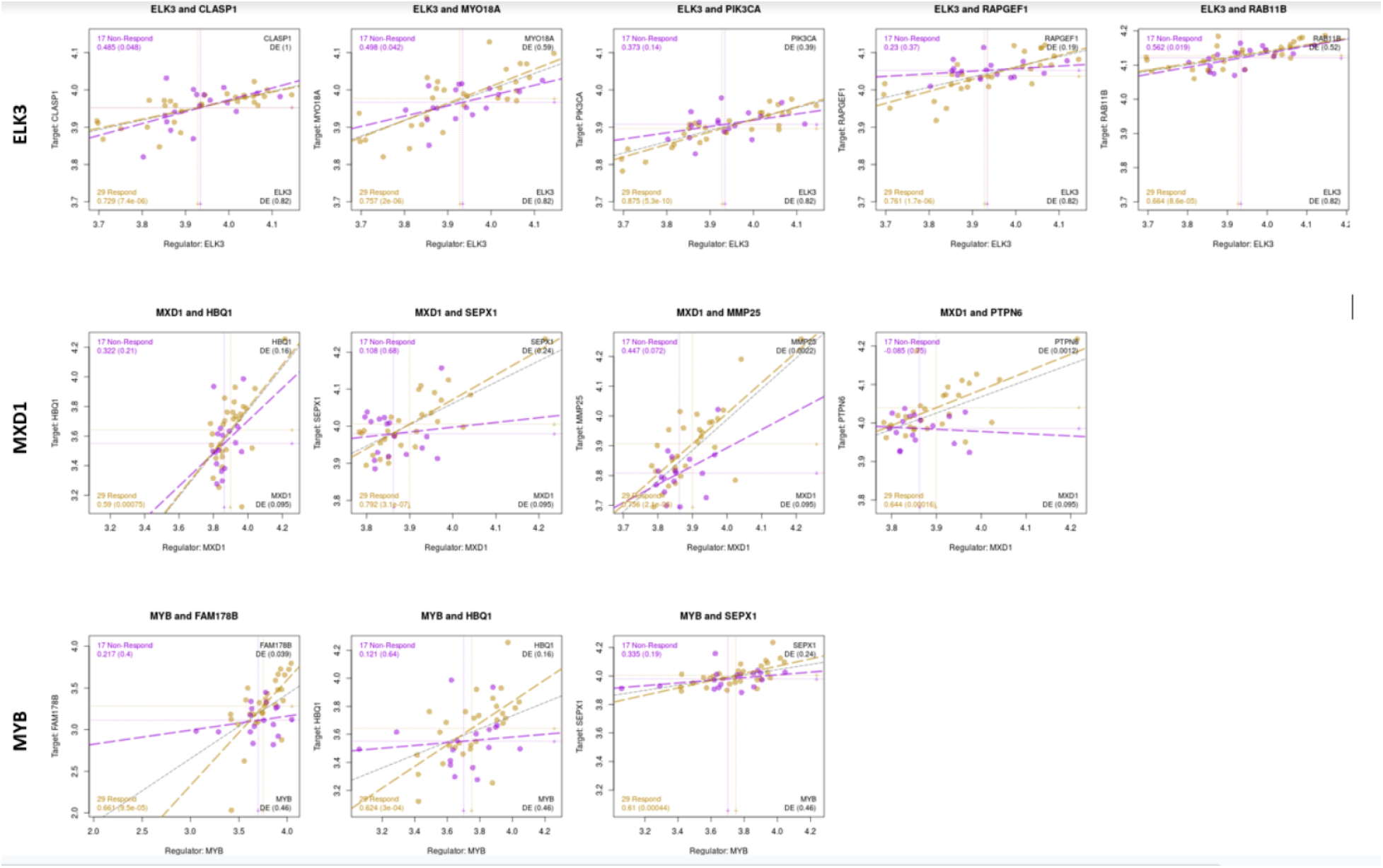
Expression Patterns in PROMOTE Data. For each of the reported experimental validations, we show the corresponding expression information from the original PROMOTE data. Points are plotted for responders (yellow) and non-responders (purple) with best fit lines drawn and correlations and their significance reported on the left hand side.

**Figure S8:**
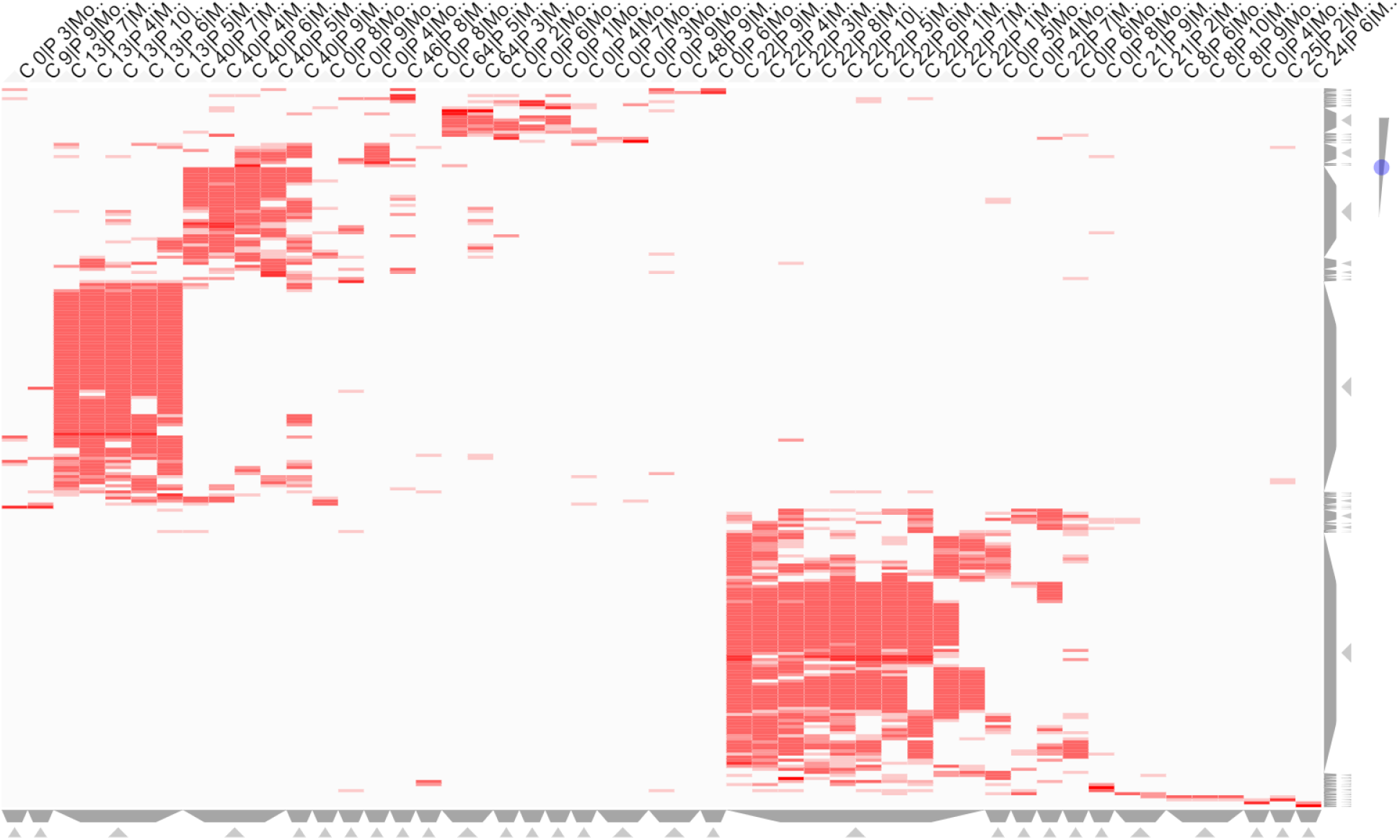
Heatmap showing the 51 rewired modules as columns labeled by their community membership (C_##, C_0 for not a member of any community) and their bootstrap run (P_##). The rows are gene targets that are in at least four of the rewired modules or regulator TFs in at least one. The red color of the cell indicates membership of the gene in the module (dark red indicates the gene is a regulator of that module). Figure generated with Clustergrammer^73^ web tool.

**Figure S9:**
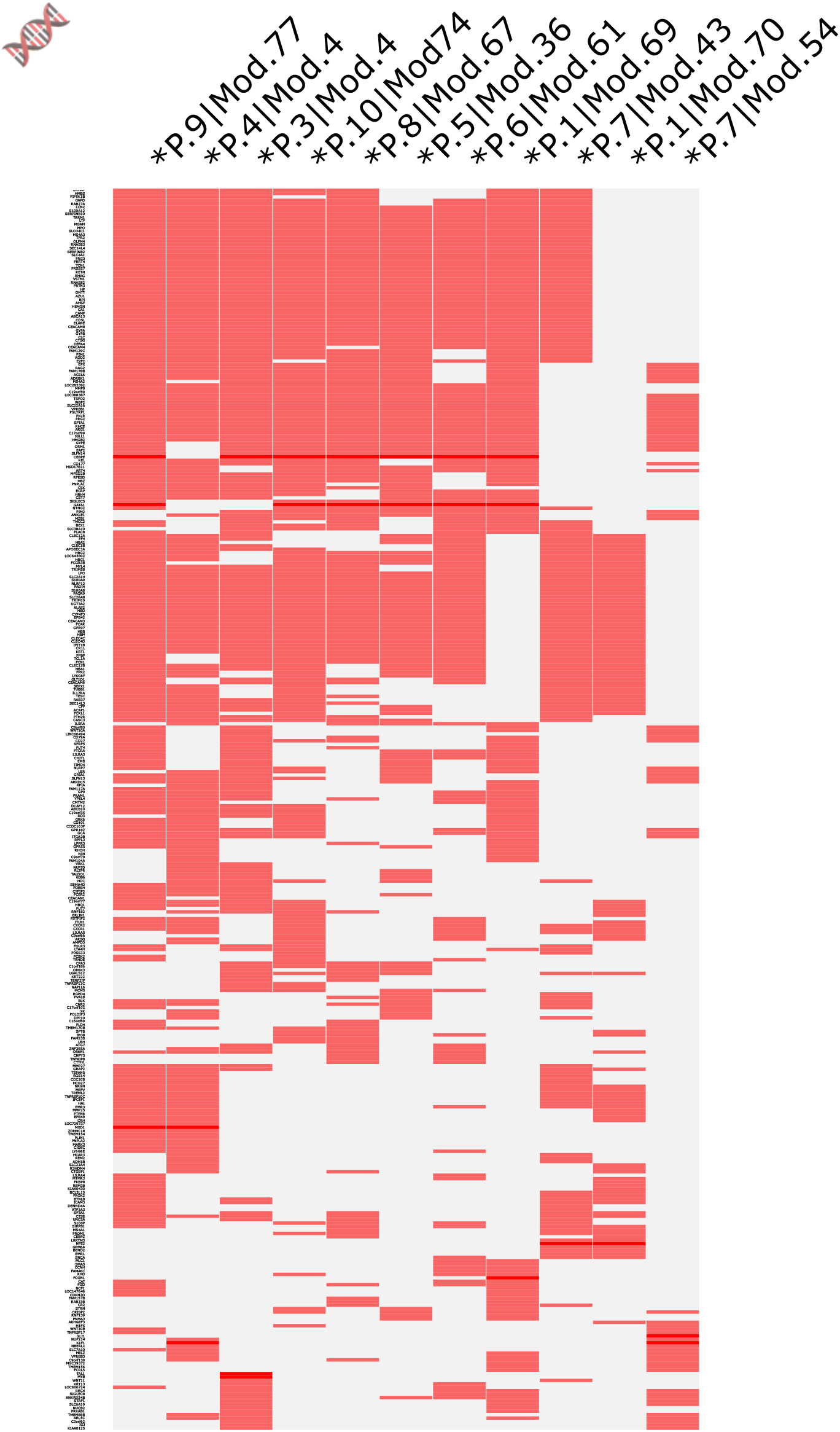
Heatmap showing the 11 modules of P-M1 as columns labeled by their bootstrap run (P.##) and within bootstrap submodule (Mod.##) and flagged with “*” if rewired. The rows are gene targets that are in at least two of the rewired modules or regulator TFs in at least one. The red color of the cell indicates membership of the gene in the module (dark red indicates the gene is a regulator of that module). Figure generated with Clustergrammer^73^ web tool.

**Figure S10:**
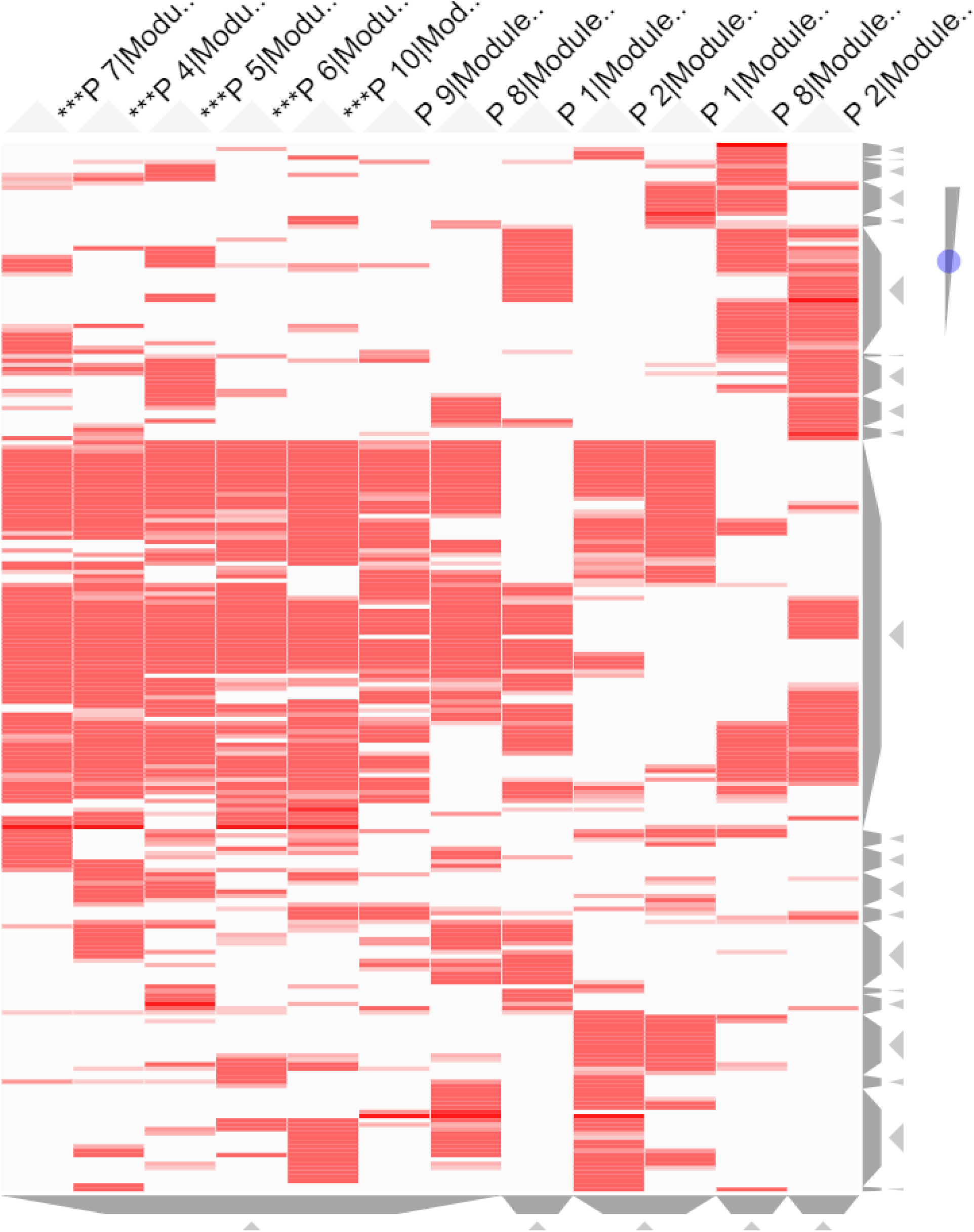
Heatmap showing the 12 modules of P-M2 as columns labeled by their bootstrap run (P_##) and flagged with “***” if rewired. The rows are gene targets that are in at least two of the rewired modules or regulator TFs in at least one. The red color of the cell indicates membership of the gene in the module (dark red indicates the gene is a regulator of that module). Figure generated with Clustergrammer^73^ web tool.

**Figure S11:**
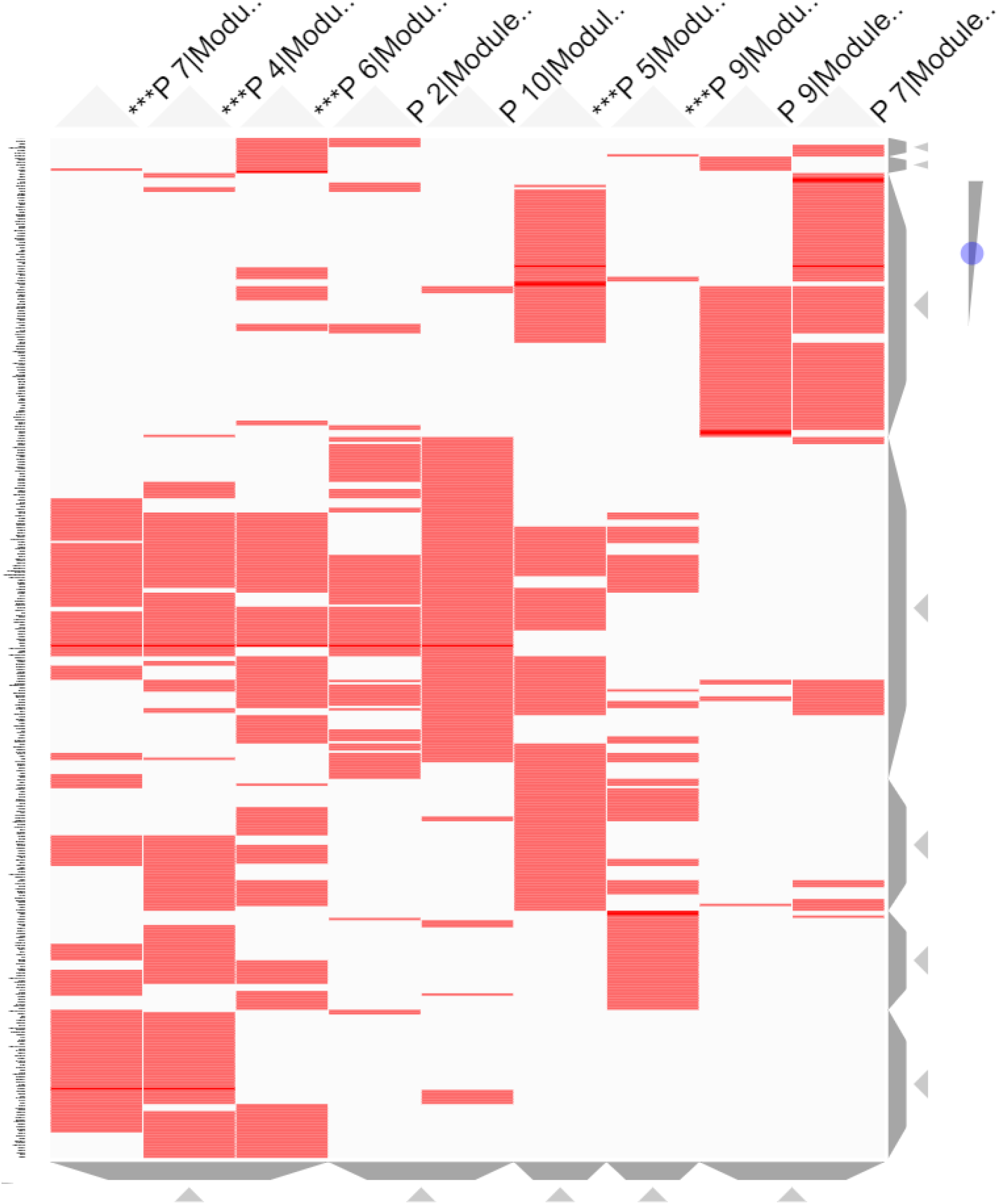
Heatmap showing the 9 modules of P-M3 as columns labeled by their bootstrap run (P_##) and flagged with “***” if rewired. The rows are gene targets that are in at least two of the rewired modules or regulator TFs in at least one. The red color of the cell indicates membership of the gene in the module (dark red indicates the gene is a regulator of that module). Figure generated with Clustergrammer^73^ web tool.

**Figure S12:**
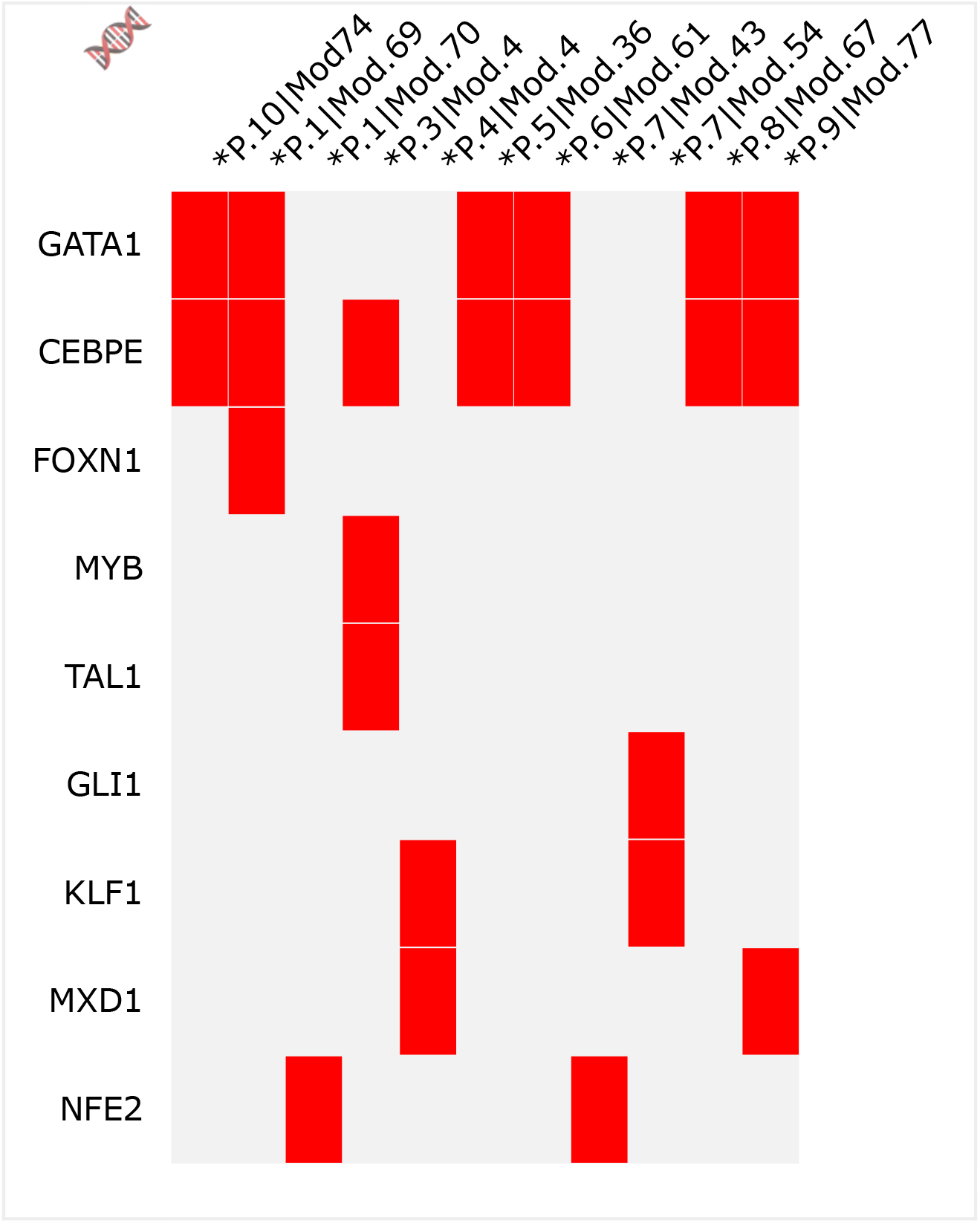
Regulators of P-M1 shown as rows of a membership heatmap with the 11 modules of P-M1 as columns labeled by their bootstrap run (P.##) and within bootstrap submodule (Mod.##) and flagged with “*” if rewired. Figure generated with Clustergrammer^73^ web tool.

## References

[1] Siegel, R. L., Miller, K. D., and Jemal, A. (2020) Cancer statistics, 2020. CA: A Cancer Journal for Clinicians, 70(1), 7–30.

[2] Litwin, M. S. and Tan, H.-J. (06, 2017) The Diagnosis and Treatment of Prostate Cancer: A Review. JAMA, 317(24), 2532–2542.

[3] James, N. D., De Bono, J. S., Spears, M. R., Clarke, N., Mason, M. D., Dearnaley, D. P., Ritchie, A. W., Russell, J. M., Gilson, C., Jones, R. J., et al. Adding abiraterone for men with high-risk prostate cancer (PCa) starting long-term androgen deprivation therapy (ADT): Survival results from STAMPEDE (NCT00268476).. (2017).

[4] Fizazi, K., Tran, N., Fein, L. E., Matsubara, N., Rodriguez Antolin, A., Alekseev, B. Y., Ozguroglu, M., Ye, D., Feyerabend, S., Protheroe, A., et al. LATITUDE: A phase III, double-blind, randomized trial of androgen deprivation therapy with abiraterone acetate plus prednisone or placebos in newly diagnosed high-risk metastatic hormone-naive prostate cancer. (2017).

[5] Petrylak, D. (2005) Therapeutic options in androgen-independent prostate cancer: building on docetaxel. BJU international, 96, 41–46.

[6] De Bono, J. S., Logothetis, C. J., Molina, A., Fizazi, K., North, S., Chu, L., Chi, K. N., Jones, R. J., Goodman Jr, O. B., Saad, F., et al. (2011) Abiraterone and increased survival in metastatic prostate cancer. New England Journal of Medicine, 364(21), 1995–2005.

[7] Scher, H. I., Fizazi, K., Saad, F., Taplin, M.-E., Sternberg, C. N., Miller, K., de Wit, R., Mulders, P., Chi, K. N., Shore, N. D., et al. (2012) Increased survival with enzalutamide in prostate cancer after chemotherapy. New England Journal of Medicine, 367(13), 1187–1197.

[8] Danila, D. C., Morris, M. J., De Bono, J. S., Ryan, C. J., Denmeade, S. R., Smith, M. R., Taplin, M.-E., Bubley, G. J., Kheoh, T., Haqq, C., et al. (2010) Phase II multicenter study of abiraterone acetate plus prednisone therapy in patients with docetaxel-treated castration-resistant prostate cancer. Journal of Clinical Oncology, 28(9), 1496.

[9] Sharma, N. L., Massie, C. E., Ramos-Montoya, A., Zecchini, V., Scott, H. E., Lamb, A. D., MacArthur, S., Stark, R., Warren, A. Y., Mills, I. G., et al. (2013) The androgen receptor induces a distinct transcriptional program in castration-resistant prostate cancer in man. Cancer cell, 23(1), 35–47.

[10] Augello, M. A., Liu, D., Deonarine, L. D., Robinson, B. D., Huang, D., Stelloo, S., Blattner, M., Doane, A. S., Wong, E. W., Chen, Y., et al. (2019) CHD1 loss alters AR binding at lineage-specific enhancers and modulates distinct transcriptional programs to drive prostate tumorigenesis. Cancer Cell, 35(4), 603–617.

[11] Wang, L., Dehm, S., Hillman, D., Sicotte, H., Tan, W., Gormley, M., Bhargava, V., Jimenez, R., Xie, F., Yin, P., et al. (2018) A prospective genome-wide study of prostate cancer metastases reveals association of wnt pathway activation and increased cell cycle proliferation with primary resistance to abiraterone acetate–prednisone. Annals of Oncology, 29(2), 352–360.

[12] Boehm, J. S. and Hahn, W. C. (2011) Towards systematic functional characterization of cancer genomes. Nature Reviews Genetics, 12(7), 487.

[13] Logsdon, B. A., Gentles, A. J., Miller, C. P., Blau, C. A., Becker, P. S., and Lee, S.-I. (2015) Sparse expression bases in cancer reveal tumor drivers. Nucleic acids research, 43(3), 1332–1344.

[14] Akavia, U. D., Litvin, O., Kim, J., Sanchez-Garcia, F., Kotliar, D., Causton, H. C., Pochanard, P., Mozes, E., Garraway, L. A., and Pe’er, D. (2010) An integrated approach to uncover drivers of cancer. Cell, 143(6), 1005–1017.

[15] Bandyopadhyay, S., Mehta, M., Kuo, D., Sung, M.-K., Chuang, R., Jaehnig, E. J., Bodenmiller, B., Licon, K., Copeland, W., Shales, M., et al. (2010) Rewiring of genetic networks in response to DNA damage. Science, 330(6009), 1385–1389.

[16] Califano, A. (2011) Rewiring makes the difference. Molecular systems biology, 7(1).

[17] Luscombe, N. M., Babu, M. M., Yu, H., Snyder, M., Teichmann, S. A., and Gerstein, M. (2004) Genomic analysis of regulatory network dynamics reveals large topological changes. Nature, 431(7006), 308.

[18] Rhinn, H., Fujita, R., Qiang, L., Cheng, R., Lee, J. H., and Abeliovich, A. (2013) Integrative genomics identifies APOE *ε*4 effectors in Alzheimer’s disease. Nature, 500(7460), 45.

[19] Taylor, I. W., Linding, R., Warde-Farley, D., Liu, Y., Pesquita, C., Faria, D., Bull, S., Pawson, T., Morris, Q., and Wrana, J. L. (2009) Dynamic modularity in protein interaction networks predicts breast cancer outcome. Nature biotechnology, 27(2), 199.

[20] De la Fuente, A. (2010) From ‘differential expression’to ‘differential networking’–identification of dysfunctional regulatory networks in diseases. Trends in genetics, 26(7), 326–333.

[21] Lai, Y., Wu, B., Chen, L., and Zhao, H. (2004) A statistical method for identifying differential gene–gene co-expression patterns. Bioinformatics, 20(17), 3146–3155.

[22] Hernaez, M., Blatti, C., and Gevaert, O. (2019) Comparison of single and module-based methods for modeling gene regulatory networks. Bioinformatics,.

[23] Gevaert, O., Van Vooren, S., and De Moor, B. (2007) A framework for elucidating regulatory networks based on prior information and expression data. Annals of the New York Academy of Sciences, 1115(1), 240–248.

[24] Logsdon, B. A., Carty, C. L., Reiner, A. P., Dai, J. Y., and Kooperberg, C. (2012) A novel variational Bayes multiple locus Z-statistic for genome-wide association studies with Bayesian model averaging. Bioinformatics, 28(13), 1738–1744.

[25] Haldane, J. (1940) The mean and variance of| chi 2, when used as a test of homogeneity, when expectations are small. Biometrika, 31(3/4), 346–355.

[26] Anscombe, F. J. (1956) On estimating binomial response relations. Biometrika, 43(3/4), 461–464.

[27] Abida, W., Cyrta, J., Heller, G., Prandi, D., Armenia, J., Coleman, I., Cieslik, M., Benelli, M., Robinson, D., Van Allen, E. M., Sboner, A., Fedrizzi, T., Mosquera, J. M., Robinson, B. D., De Sarkar, N., Kunju, L. P., Tomlins, S., Wu, Y. M., Nava Rodrigues, D., Loda, M., Gopalan, A., Reuter, V. E., Pritchard, C. C., Mateo, J., Bianchini, D., Miranda, S., Carreira, S., Rescigno, P., Filipenko, J., Vinson, J., Montgomery, R. B., Beltran, H., Heath, E. I., Scher, H. I., Kantoff, P. W., Taplin, M.-E., Schultz, N., deBono, J. S., Demichelis, F., Nelson, P. S., Rubin, M. A., Chinnaiyan, A. M., and Sawyers, C. L. (2019) Genomic correlates of clinical outcome in advanced prostate cancer. Proceedings of the National Academy of Sciences, 116(23), 11428–11436.

[28] Cerami, E., Gao, J., Dogrusoz, U., Gross, B. E., Sumer, S. O., Aksoy, B. A., Jacobsen, A., Byrne, C. J., Heuer, M. L., Larsson, E., et al. The cBio cancer genomics portal: an open platform for exploring multidimensional cancer genomics data. (2012).

[29] Human Protein Atlas 2020 v19.3. https://www.proteinatlas.org/search/protein_class:Transcription+factors.

[30] Champion, M., Brennan, K., Croonenborghs, T., Gentles, A. J., Pochet, N., and Gevaert, O. (2018) Module analysis captures pancancer genetically and epigenetically deregulated cancer driver genes for smoking and antiviral response. EBioMedicine, 27, 156–166.

[31] Newman, M. E. J. and Girvan, M. (Feb, 2004) Finding and evaluating community structure in networks. Phys. Rev. E, 69, 026113.

[32] Liberzon, A., Birger, C., Thorvaldsdóttir, H., Ghandi, M., Mesirov, J. P., and Tamayo, P. (2015) The molecular signatures database hallmark gene set collection. Cell systems, 1(6), 417–425.

[33] Gevaert, O., Villalobos, V., Sikic, B. I., and Plevritis, S. K. (8, 2013) Identification of ovarian cancer driver genes by using module network integration of multi-omics data. Interface Focus, 3.

[34] Manolakos, A., Ochoa, I., Venkat, K., Goldsmith, A. J., and Gevaert, O. (2014) CaMoDi: a new method for cancer module discovery. BMC Genomics, 15, S8.

[35] Shyamsunder, P., Shanmugasundaram, M., Mayakonda, A., Dakle, P., Teoh, W. W., Han, L., Kanojia, D., Lim, M. C., Fullwood, M., An, O., et al. (2019) Identification of a novel enhancer of CEBPE essential for granulocytic differentiation. *Blood*, The Journal of the American Society of Hematology, 133(23), 2507–2517.

[36] Evrard, M., Kwok, I. W., Chong, S. Z., Teng, K. W., Becht, E., Chen, J., Sieow, J. L., Penny, H. L., Ching, G. C., Devi, S., et al. (2018) Developmental analysis of bone marrow neutrophils reveals populations specialized in expansion, trafficking, and effector functions. Immunity, 48(2), 364–379.

[37] Gilles, L., Arslan, A. D., Marinaccio, C., Wen, Q. J., Arya, P., McNulty, M., Yang, Q., Zhao, J. C., Konstantinoff, K., Lasho, T., et al. (2017) Downregulation of GATA1 drives impaired hematopoiesis in primary myelofibrosis. The Journal of clinical investigation, 127(4), 1316–1320.

[38] Wang, X., Angelis, N., and Thein, S. L. (2018) MYB–A regulatory factor in hematopoiesis. Gene, 665, 6–17.

[39] Gautam, S., Fioravanti, J., Zhu, W., Le Gall, J. B., Brohawn, P., Lacey, N. E., Hu, J., Hocker, J. D., Hawk, N. V., Kapoor, V., et al. (2019) The transcription factor c-Myb regulates CD8+ T cell stemness and antitumor immunity. Nature immunology, 20(3), 337–349.

[40] Rosati, R., Patki, M., Chari, V., Dakshnamurthy, S., McFall, T., Saxton, J., Kidder, B. L., Shaw, P. E., and Ratnam, M. (2016) The amino-terminal domain of the androgen receptor co-opts extracellular signal-regulated kinase (ERK) docking sites in ELK1 protein to induce sustained gene activation that supports prostate cancer cell growth. Journal of Biological Chemistry, 291(50), 25983–25998.

[41] Li, X., Wu, J. B., Li, Q., Shigemura, K., Chung, L. W., and Huang, W.-C. (2016) SREBP-2 promotes stem cell-like properties and metastasis by transcriptional activation of c-Myc in prostate cancer. Oncotarget, 7(11), 12869.

[42] Thakur, N., Hamidi, A., Song, J., Itoh, S., Bergh, A., Heldin, C.-H., and Landström, M. (2020) Smad7 Enhances TGF-*β*-Induced Transcription of c-Jun and HDAC6 Promoting Invasion of Prostate Cancer Cells. Iscience, 23(9), 101470.

[43] Sun, H., Wang, H., Wang, X., Aoki, Y., Wang, X., Yang, Y., Cheng, X., Wang, Z., and Wang, X. (2020) Aurora-A/SOX8/FOXK1 signaling axis promotes chemoresistance via suppression of cell senescence and induction of glucose metabolism in ovarian cancer organoids and cells. Theranostics, 10(15), 6928.

[44] Xie, S.-L., Fan, S., Zhang, S.-Y., Chen, W.-X., Li, Q.-X., Pan, G.-K., Zhang, H.-Q., Wang, W.-W., Weng, B., Zhang, Z., et al. (2018) SOX8 regulates cancer stem-like properties and cisplatin-induced EMT in tongue squamous cell carcinoma by acting on the Wnt/*β*-catenin pathway. International journal of cancer, 142(6), 1252–1265.

[45] Fan, L., Lei, H., Zhang, S., Peng, Y., Fu, C., Shu, G., and Yin, G. (2020) Non-canonical signaling pathway of SNAI2 induces EMT in ovarian cancer cells by suppressing miR-222-3p transcription and upregulating PDCD10. Theranostics, 10(13), 5895.

[46] Alves, C. C., Carneiro, F., Hoefler, H., and Becker, K.-F. (2009) Role of the epithelial-mesenchymal transition regulator Slug in primary human cancers. Front Biosci, 14(1), 3035–50.

[47] Kim, S., Yao, J., Suyama, K., Qian, X., Qian, B.-Z., Bandyopadhyay, S., Loudig, O., De Leon-Rodriguez, C., Zhou, Z. N., Segall, J., et al. (2014) Slug promotes survival during metastasis through suppression of Puma-mediated apoptosis. Cancer research, 74(14), 3695–3706.

[48] Li, W., Shen, M., Jiang, Y.-Z., Zhang, R., Zheng, H., Wei, Y., Shao, Z.-M., and Kang, Y. (2020) Deubiquitinase USP20 promotes breast cancer metastasis by stabilizing SNAI2. Genes & development, 34(19-20), 1310–1315.

[49] Helgeson, B. E., Tomlins, S. A., Shah, N., Laxman, B., Cao, Q., Prensner, J. R., Cao, X., Singla, N., Montie, J. E., Varambally, S., et al. (2008) Characterization of TMPRSS2: ETV5 and SLC45A3: ETV5 gene fusions in prostate cancer. Cancer research, 68(1), 73–80.

[50] Szklarczyk, D., Gable, A. L., Lyon, D., Junge, A., Wyder, S., Huerta-Cepas, J., Simonovic, M., Doncheva, N. T., Morris, J. H., Bork, P., Jensen, L. J., and Mering, C. v. (11, 2018) STRING v11: protein–protein association networks with increased coverage, supporting functional discovery in genome-wide experimental datasets. Nucleic Acids Research, 47(D1), D607–D613.

[51] Vagapova, E., Spirin, P., Lebedev, T., and Prassolov, V. (2018) The role of TAL1 in hematopoiesis and leukemogenesis. Acta Naturae, 10(1 (36)), 15–23.

[52] Langfelder, P. and Horvath, S. (2008) WGCNA: an R package for weighted correlation network analysis. BMC bioinformatics, 9(1), 1–13.

[53] Alvarez, M. J., Shen, Y., Giorgi, F. M., Lachmann, A., Ding, B. B., Ye, B. H., and Califano, A. (2016) Functional characterization of somatic mutations in cancer using network-based inference of protein activity. Nature genetics, 48(8), 838.

[54] Califano, A. and Alvarez, M. J. (2017) The recurrent architecture of tumour initiation, progression and drug sensitivity. Nature reviews Cancer, 17(2), 116.

[55] Hudson, N. J., Reverter, A., and Dalrymple, B. P. (2009) A differential wiring analysis of expression data correctly identifies the gene containing the causal mutation. PLoS computational biology, 5(5), e1000382.

[56] Reverter, A., Ingham, A., Lehnert, S. A., Tan, S.-H., Wang, Y., Ratnakumar, A., and Dalrymple, B. P. (2006) Simultaneous identification of differential gene expression and connectivity in inflammation, adipogenesis and cancer. Bioinformatics, 22(19), 2396–2404.

[57] Ha, M. J., Baladandayuthapani, V., and Do, K.-A. (2015) DINGO: differential network analysis in genomics. Bioinformatics, 31(21), 3413–3420.

[58] Yu, D., Lim, J., Wang, X., Liang, F., and Xiao, G. (2017) Enhanced construction of gene regulatory networks using hub gene information. BMC bioinformatics, 18(1), 186.

[59] Ohba, Y., Ikuta, K., Ogura, A., Matsuda, J., Mochizuki, N., Nagashima, K., Kurokawa, K., Mayer, B. J., Maki, K., Miyazaki, J.-i., et al. (2001) Requirement for C3G-dependent Rap1 activation for cell adhesion and embryogenesis. The EMBO journal, 20(13), 3333–3341.

[60] Nakatsuji, H., Nishimura, N., Yamamura, R., Kanayama, H.-o., and Sasaki, T. (2008) Involvement of actinin-4 in the recruitment of JRAB/MICAL-L2 to cell-cell junctions and the formation of functional tight junctions. Molecular and cellular biology, 28(10), 3324–3335.

[61] Erasmus, J. C., Smolarczyk, K., Brezovjakova, H., Mohd-Naim, N. F., Lozano, E., Matter, K., and Braga, V. M. (2021) Rac1-PAK1 regulation of Rab11 cycling promotes junction destabilization. Journal of Cell Biology, 220(6), e202002114.

[62] Liu, J., Ren, G., Li, K., Liu, Z., Wang, Y., Chen, T., Mu, W., Yang, X., Li, X., Shi, A., et al. (2021) The Smad4-MYO18A-PP1A complex regulates *β*-catenin phosphorylation and pemigatinib resistance by inhibiting PAK1 in cholangiocarcinoma. Cell Death & Differentiation, pp. 1–14.

[63] by Metastatic, D. I. (2006) Role of Focal Adhesion Kinase and Phosphatidylinositol 3. Cancer Res, 66(16), 8091–9.

[64] Mao, Y., Li, W., Hua, B., Gu, X., Pan, W., Chen, Q., Xu, B., Wang, Z., and Lu, C. (2020) Silencing of ELK3 induces SM phase arrest and apoptosis and upregulates SERPINE1 expression reducing migration in prostate cancer cells. BioMed Research International, 2020.

[65] Kar, A. and Gutierrez-Hartmann, A. (2013) Molecular mechanisms of ETS transcription factor-mediated tumorigenesis. Critical reviews in biochemistry and molecular biology, 48(6), 522–543.

[66] Abeshouse, A., Ahn, J., Akbani, R., Ally, A., Amin, S., Andry, C. D., Annala, M., Aprikian, A., Armenia, J., Arora, A., et al. (2015) The molecular taxonomy of primary prostate cancer. Cell, 163(4), 1011–1025.

[67] Clark, J. P. and Cooper, C. S. (2009) ETS gene fusions in prostate cancer. Nature Reviews Urology, 6(8), 429–439.

[68] Drabsch, Y., Ramsay, R. G., and Gonda, T. J. (2010) MYB suppresses differentiation and apoptosis of human breast cancer cells. Breast Cancer Research, 12(4), 1–17.

[69] Zheng, D., Wu, W., Dong, N., Jiang, X., Xu, J., Zhan, X., Zhang, Z., and Hu, Z. (2017) Mxd1 mediates hypoxia-induced cisplatin resistance in osteosarcoma cells by repression of the PTEN tumor suppressor gene. Molecular Carcinogenesis, 56(10), 2234–2244.

[70] Ye, Y.-P., Jiao, H.-L., Wang, S.-Y., Xiao, Z.-Y., Zhang, D., Qiu, J.-F., Zhang, L.-J., Zhao, Y.-L., Li, T.-T., Liao, W.-T., et al. (2018) Hypermethylation of DMTN promotes the metastasis of colorectal cancer cells by regulating the actin cytoskeleton through Rac1 signaling activation. Journal of Experimental & Clinical Cancer Research, 37(1), 1–14.

[71] Novoselov, S. V., Kim, H.-Y., Hua, D., Lee, B. C., Astle, C. M., Harrison, D. E., Friguet, B., Moustafa, M. E., Carlson, B. A., Hatfield, D. L., et al. (2010) Regulation of selenoproteins and methionine sulfoxide reductases A and B1 by age, calorie restriction, and dietary selenium in mice. Antioxidants & redox signaling, 12(7), 829–838.

[72] De La Fuente Cedeño, J., Hernaez, M., and Blatti, C. TraRe: Transcriptional Rewiring (2021) R package version 1.2.0.

[73] Fernandez, N. F., Gundersen, G. W., Rahman, A., Grimes, M. L., Rikova, K., Hornbeck, P., and Ma’ayan, A. (2017) Clustergrammer, a web-based heatmap visualization and analysis tool for high-dimensional biological data. Scientific data, 4(1), 1–12.

