## Supplementary material for "Identification of transcriptional network disruptions in drug-resistant prostate cancer with TraRe": Suplementary Data

C. Blatti, J. de la Fuente, H. Gao, I. Marin, Z. Chen, S. D. Zhao,
W. Tan, R. Weinshilbaum, K. R. Kalari, L. Wang, M. Hernaez

### S2 Additional Tables

| Regulatory module | Nr bootstraps | Nr modules | Nr rewired modules | % of rewired modules | TFs | Nr TFs | Nr targets |
|---|---|---|---|---|---|---|---|
| P-M1 | 9 | 11 | 11 | 100 | CEBPE, GATA1, KLF1, MXD1, NFE2, FOXN1, GLI1, MYB, TAL1 | 9 | 619 |
| P-M2 | 9 | 12 | 5 | 41.7 | ZNF91, ZKSCAN1, ZNF519, ELK1, HOXA6, SREBF2, ZBTB26, ZFHX4, ZNF607, ZNF778, ZNF827 | 11 | 1045 |
| P-M3 | 7 | 9 | 5 | 55.6 | PAX6, SMAD7, ZNF670, LEF1, SOX8, STAT2, ZBTB43, ZBTB45, ZFAT, ZFP37, ZNF286A, ZNF69 | 12 | 879 |
| P-M4 | 10 | 10 | 3 | 30 | SNAI2, FOXC2, RARG, RUNX2, SMAD1, TSHZ3, ZNF655 | 7 | 1172 |
| P-M5 | 7 | 7 | 2 | 28.6 | NR1H4, HOXD12 | 2 | 475 |
| P-M6 | 10 | 17 | 1 | 5.9 | MEF2C, MSC, FLI1, MEOX1, PRRX1, ZNF655, BATF3, HIF3A, KLF10, KLF15, KLF6, KLF8, MYOD1, NKX3-1, NONO, PAX3, SOX18, TCF7L1, THAP11, ZBTB42, ZNF607, ZNF610 | 22 | 2337 |
| P-M7 | 10 | 12 | 1 | 8.3 | ELK3, FOXC2, DNMT1, ZHX2, ZNF496 | 5 | 910 |
| P-M8 | 9 | 14 | 1 | 7.1 | ZNF709, ZSCAN22, ZNF670, DLX1, E2F4, MAF, MNT, MXD3, ZNF16, ZNF174, ZNF192, ZNF347, ZNF394, ZNF528, ZNF789, ZNF800, ZSCAN23 | 17 | 1238 |
| P-M9 | 6 | 7 | 1 | 14.3 | EBF1, RUNX1, ETV5, FOXA1, HOXB2, NFATC3, NFATC4, NR6A1, RBPJ, RUNX2, SP100, ZNF689 | 12 | 1137 |

**Table S1:** Summary table of regulatory modules generated with cAMARETTO software[30] from all sub-modules obtained in a 10-bootstrap run with TraRe on PROMOTE dataset (Part 1: Rewired Regulatory Modules).

| Regulatory module | Nr bootstraps | Nr modules | Nr TFs | Nr targets |
|---|---|---|---|---|
| P-M10 | 10 | 28 | 16 | 1834 |
| P-M11 | 10 | 21 | 17 | 1524 |
| P-M12 | 10 | 19 | 24 | 1848 |
| P-M13 | 10 | 13 | 10 | 830 |
| P-M14 | 10 | 13 | 13 | 1532 |
| P-M15 | 10 | 12 | 9 | 1115 |
| P-M16 | 10 | 12 | 12 | 1142 |
| P-M17 | 10 | 11 | 11 | 1128 |
| P-M18 | 10 | 10 | 8 | 1201 |
| P-M19 | 10 | 10 | 9 | 1394 |
| P-M20 | 10 | 10 | 6 | 870 |
| P-M21 | 10 | 10 | 4 | 1482 |
| P-M22 | 10 | 10 | 9 | 792 |
| P-M23 | 9 | 11 | 10 | 1028 |
| P-M24 | 9 | 10 | 10 | 1217 |
| P-M25 | 9 | 10 | 16 | 1238 |
| P-M26 | 9 | 10 | 11 | 1411 |
| P-M27 | 8 | 8 | 9 | 766 |
| P-M28 | 7 | 8 | 5 | 591 |
| P-M29 | 7 | 8 | 10 | 1108 |
| P-M30 | 7 | 8 | 6 | 657 |
| P-M31 | 7 | 7 | 1 | 681 |
| P-M32 | 7 | 7 | 6 | 468 |
| P-M33 | 7 | 7 | 11 | 815 |
| P-M34 | 6 | 7 | 5 | 665 |
| P-M35 | 6 | 6 | 9 | 961 |
| P-M36 | 5 | 6 | 5 | 823 |
| P-M37 | 5 | 5 | 5 | 759 |
| P-M38 | 5 | 5 | 2 | 511 |

**Table S2:** Summary table of regulatory modules generated with cAMARETTO software[30] from all sub-modules obtained in a 10-bootstrap run with TraRe on PROMOTE dataset (Part 2: Non-rewired Regulatory Modules).

| Regulatory module | TFs |
|---|---|
| P-M10 | TGIF2LX, NANOGNB, HSFY1, DMRTC1, DRGX, HSFY2, MIXL1, ZNF679, BARHL1, DUXA, GFI1, HOXB1, ZNF426, ZNF534, ZNF774, ZNF92 |
| P-M11 | SP7, DLX5, HDX, ZBTB7C, RFX8, DLX3, FOSL2, FOXP2, JUNB, KLF2, NR5A2, PRRX2, RCOR2, TGIF1, ZNF469, ZNF610, ZNF672 |
| P-M12 | ZNF224, HIF1A, ZNF808, MIXL1, RC3H2, ZBTB26, ATF4, ETV1, NEUROG3, SRF, TWIST2, WRNIP1, ZBED3, ZBED6, ZFP90, ZNF131, ZNF284, ZNF426, ZNF611, ZNF625, ZNF749, ZNF765, ZNF774, ZSCAN23 |
| P-M13 | ZNF83, EWSR1, POGZ, ZNF789, CCNT2, IRF3, NR2C1, VANGL1, ZNF518A, ZNF700 |
| P-M14 | IRF6, HOXB13, EHF, TFAP2C, BACH2, FOXA1, HLX, STAT5A, THAP10, ZNF295, ZNF607, ZNF613, ZNF786 |
| P-M15 | BSX, HOXB1, CNOT4, DBX1, DRGX, FOXD4L5, NFATC4, ZNF774, ZSCAN21 |
| P-M16 | IKZF1, LYL1, BCL11B, GRHL2, ID2, IRF1, MTA3, NFKBID, SMARCC1, STAT4, STAT5B, TBX21 |
| P-M17 | TFEC, ETV5, STAT5A, ADNP2, FLI1, POU2F2, SP140, SPI1, VENTX, ZNF217, ZSCAN16 |
| P-M18 | NEUROG1, BSX, ESX1, HSFX2, NFE2L2, RHOXF2B, TWIST2, ZNF710 |
| P-M19 | ZIM3, ZNF679, HSFY1, HSFY2, DMRTB1, SOHLH2, TEAD4, ZNF683, ZNF836 |
| P-M20 | MYF6, NFE2L1, GRHL2, MYOD1, TEAD4, ZNF713 |
| P-M21 | RHOXF2, RHOXF2B, HSFY2, NEUROG1 |
| P-M22 | MYBL2, DNMT1, E2F1, PPARA, E2F7, FOXK2, GFI1, TBX10, ZNF878 |
| 23 | EPAS1, SOX18, BCL6B, ETV5, HLX, KLF8, MEIS1, ZBTB7B, ZNF239, ZNF829 |
| P-M24 | ZNF281, ZNF550, ELF4, HES1, RUNX2, ZNF192, ZNF284, ZNF616, ZNF746, ZNF8 |
| P-M25 | ZNF354B, DNMT1, ELK3, THAP3, ELF4, FOXC2, NFATC3, RFXANK, STAT3, ZBTB24, ZNF124, ZNF192, ZNF225, ZNF248, ZNF354A, ZNF791 |
| P-M26 | GRHL2, DMRT2, HDX, HESX1, HOXB13, PPP1R13L, SNAI3, ZNF667, ZNF670, ZNF674, ZNF789 |
| P-M27 | ZNF749, ZBTB32, LHX4, CNTN2, RBPJ, TFCP2L1, ZBTB7C, ZNF28, ZNF286A |
| P-M28 | TBX18, ZNF418, AIRE, IRX1, YY2 |
| P-M29 | DLX5, GRHL2, ZNF551, ZNF670, AHR, CSRNP3, SIX4, SP140, ZNF281, ZNF709 |
| P-M30 | SCRT1, MESP2, FOXD4, NFATC2, POU5F1, ZNF223 |
| P-M31 | ZNF692 |

| **Regulatory module** | **TFs** |
|---|---|
| P-M32 | ETV1, HMGN3, TBX19, ZNF133, ZNF180, ZNF322 |
| P-M33 | NFE2L2, ZNF814, CREB3L4, ELF1, ELF4, HOXA7, MIXL1, NFATC2, SMAD1, THAP11, ZNF624 |
| P-M34 | FOXI1, LHX8, TFAP2D, ZNF679, ZIM3 |
| P-M35 | ZNF713, ZNF121, KLF2, THAP11, ZBTB25, ZNF142, ZNF28, ZNF521, ZNF746 |
| P-M36 | ELF1, ELK3, ZNF205, MEIS1, ZNF33B |
| P-M37 | ZKSCAN3, PRDM15, FOXH1, RAX, TBX10 |
| P-M38 | ZNF426, HESX1 |

**Table S3:** Summary table of regulatory modules generated with cAMARETTO software[30] from all sub-modules obtained in a 10-bootstrap run with TraRe on PROMOTE dataset (Part 3: Non-rewired Regulatory Modules TF list).

| TF | Targets | p_value | multiplicity |
|---|---|---|---|
| ELK3 | CLASP1 | 9.28E-05 | 9 |
| ELK3 | SH3BP5 | 9.28E-05 | 9 |
| ELK3 | ARL10 | 9.28E-05 | 9 |
| ELK3 | SEPP1 | 9.28E-05 | 9 |
| ELK3 | RAPGEF1 | 9.28E-05 | 9 |
| ELK3 | MYO18A | 9.28E-05 | 9 |
| ELK3 | DUSP18 | 9.28E-05 | 9 |
| ELK3 | SRGAP2P2 | 2.05E-04 | 8 |
| ELK3 | ARL15 | 2.05E-04 | 8 |
| ELK3 | FAM65A | 2.05E-04 | 8 |
| ELK3 | TAX1BP3 | 1.92E-04 | 8 |
| ELK3 | RAB11B | 2.05E-04 | 8 |
| ELK3 | ACTN4 | 4.56E-04 | 7 |
| ELK3 | MAP3K12 | 4.66E-04 | 7 |
| ELK3 | PIK3CA | 4.66E-04 | 7 |
| ELK3 | NCK1 | 1.06E-03 | 6 |

**Table S4:** ELK3 regulon with selected targets for biological validation

| TF | Targets | p_value | multiplicity |
|---|---|---|---|
| MXD1 | GCA | 1.07E-04 | 9 |
| MXD1 | MMP25 | 2.42E-04 | 8 |
| MXD1 | PTPN6 | 4.64E-04 | 7 |
| MXD1 | HBQ1 | 5.83E-04 | 7 |
| MXD1 | DMTN | 1.13E-03 | 6 |
| MXD1 | KEL | 1.28E-02 | 3 |

**Table S5:** MXD1 regulon with selected targets for biological validation

| TF | Targets | p_value | multiplicity |
|---|---|---|---|
| MYB | ITGA2B | 8.95E-03 | 4 |
| MYB | FAM178B | 4.40E-02 | 2 |
| MYB | ERMAP | 3.77E-02 | 1 |
| MYB | GCA | 3.77E-02 | 1 |
| MYB | NRGN | 3.77E-02 | 1 |
| MYB | HBQ1 | 3.77E-02 | 1 |
| MYB | MSRB1 | 3.77E-02 | 1 |

**Table S6:** MYB regulon with selected targets for biological validation

| TF | target | p_value | multiplicity |
|---|---|---|---|
| ZNF91 | GEMIN5 | 1.39E-03 | 5 |
| ZNF91 | RSBN1 | 3.19E-03 | 4 |
| ZNF91 | MACROD2 | 6.48E-03 | 3 |

**Table S7:** ZNF91 regulon with selected targets for biological validation

| TF | target | p_value | multiplicity |
|---|---|---|---|
| ZNF3 | NUDCD3 | 3.06E-05 | 8 |
| ZNF3 | STYXL1 | 3.06E-05 | 8 |
| ZNF3 | LAMTOR4 | 9.40E-05 | 7 |
| ZNF3 | PSMG3 | 1.90E-04 | 6 |
| ZNF3 | DTX2 | 5.92E-04 | 5 |
| ZNF3 | LRRC27 | 3.96E-02 | 1 |

**Table S8:** ZNF3 regulon with selected targets for biological validation

| Gene | Sequence | Gene | Sequence |
|---|---|---|---|
| MXD1 | TGG TAA CAT GGA GGC ATA ACC<br>GGC GGT TCG GAT GAA CA | RAPGEF1 | CTT CTG AGT TCA CGC CTT CC<br>GTG CAG AAC GAT CCT CGA AT |
| MYB | CTC CTG CAG ATA ACC TTC CTG<br>GCA GAA ATC GCA AAG CTA CTG | MYO18A | CGG TTC TGG ATC TCA TTC TTC T<br>GAA AGC ACG GAA AGC AAT GG |
| ZNF3 | TCT GAC TTT TGT TGA TGC CAA TG<br>ACT TCC GTT CTT TGT TCT GTC C | NUDCD3 | TCT CTT CAA GTT CCT GCC TTC<br>GAC AGA CTT CTA TCG CTT GCT |
| ELK3 | GCA ACT GCA ACA GGA ACT G<br>CTG TCA GCA TGG AAA GTC G | GCA | GGC AAT CAT AAT TCT GCA GGT T<br>CCG TGT ATA CTT ACT TCA GTG CT |
| ZNF91 | CCA GTG TGT ATC CTC TTA TGT CTT C<br>TCA TTC CTC AAG TCT TTC TAC ACA | RSBN1 | GTT ATG CGA GGT TGG TCA CT<br>CTA GAG CTC ATG CTG ATC ATG T |
| LAMTOR4 | GCC TGC TCA TCA TTC TCC A<br>AGA CTG CGA TGA CTT CTG C | SEPP1 | TCT TCA GTT TTA CTC GCA GGT C<br>CAA GAT CCA ATG CTA AAC TCC AA |
| ACTN4 | GCT TCT CGT AGT CCT CCA TCA<br>CCA GCT TCT ACC ATG CCT TT | ITGA2B | ACC CTC CTG CTA GAA TAG TGT A<br>CTG CTG CTC ACC ATC CTG |
| DMTN | GGA GGT AAG TGG TTG CTT CT<br>TCT ACA GAT GCC ATC AAC GAG | MSRB1 | GAG AAC AGC TCA TAG CCA CA<br>GCA GCC TTT GGT CAG TTG |
| PIK3CA | TGC TGT CGA ATA GCT AGA TAA GC<br>AGT GAT TAG TAA AGG AGC CCA AG | LRRC27 | CTC CAC AGG TAA CAT TTT GAT AGG<br>TCA AGA TTT CTT TCA GTT GCT TCC |
| PTPN6 | CGC AGT TGG TCA CAG AGT AG<br>CAG CCG TGT CAT CGT CAT | HBQ1 | TGG GAG AAG TAG GTC TTC GT<br>CGC CCT GTG GAA GAA GC |
| RAB11B | CTC TTG CTC TCC AGG TTG AAC<br>GAC GAC GAG TAC GAC TAC CTA T | ARL15 | GCA TCC ATG TCA TCC AGT GA<br>CAG CTC GCT CAG TAC AAG AG |
| SH3BP5 | GAG TTT CTT CTC CAG CTG TCG<br>GAT GCT GAA TCA CGC CAC T | KEL | CTT CTG TTC TTG ATC TTG CTT GAG<br>TTT CCC TTT CTT CAG AGC CTA C |
| FAM178B | GTG ATG TCC TGG CAG CTA AC<br>AGG AGC AAC AGC CAA AGG | DUSP18 | GTC AGT GGT CAG CAG TCA G<br>CTA CTT CCT GAG CGA ACC C |
| NCK1 | GCC CAA TGC AGT AGA CAG TC<br>GGC ATT AAA TGA AAG AGG ACA TGA | ERMAP | AGG ATG AGT TTG GGA TGT GC<br>GTG AAC TGA AGT TGA AAA GAG CTG |
| ARL10 | CCA GTA GAA GCG CAG GTT C<br>TTC AAC TCC GTG CGT CTG | DTX2 | GTC CCC GTC TTC TCT CCA TA<br>CTC ACC AAG TGC AGC CAT |
| MMP25 | CCA AAC TCA TGG ACA GCC A<br>CCT AGC CCA TGC CTT CTT C | STYXL1 | GCA TGA GGT AGG CTA TGA TGG<br>GCC CAG ATT CTT CCC TTC TTA C |
| GEMIN5 | CCT CGA AGC CTA TGA ATA ACT TCT<br>TCT GTC TTA CTT GTT CAC CTC ATC | MAP3K12 | CTC TTG TCA CTC AGC TCC TT<br>ACC TGC ACA AGA TTA TCC ACA |
| MACROD2 | CAT GTC CAG TAT CAC AGC CA<br>TCA CAT TGC TAG AGG TAG ATG C | FAM65A | TAT GCC TCA TAC AGC TCA TCG<br>GCA GAT AAG GGA GTC CAA GAG |
| CLASP1 | CAC CAT AAC CAC TTC CCA ACA<br>CAC AAG ACG GCA AAG CTC T | TAX1BP3 | ATG TCC CAG CCG TTC AC<br>CCG TGG TGC AAA GAG TTG A |
| SRGAP2P2 | GGC AGA GAT TGA GAT GGA CTA C<br>TCC TCC TGC TTT ACC GAT TTA C | NRGN | GAA AAC TCG CCT GGA TTT TGG<br>CGA CAC CAG CAT GGA CTG |
| PSMG3 | CCT CCA TGC TTT TGT CCT TCA<br>CTT CTG GGG CAG GAT GAG | | |

**Table S9:** Primer sequences for qRT-PCR assays

| Pool Catalog Number | Gene Symbol | Sequence |
|---|---|---|
| M-009325-01 | MXD1 | AAAGCCAAAUUGCACAUAA<br>GAAUAGACGGGCUCAUCUU<br>GAAUCAAGUCGACACACUA<br>GCAGAUCAACUCACAAUGA<br>GAUCUCCUCUUUAAUGUUG |
| M-010320-00 | ELK3 | GAACGAUGGUGAAUUCAAG<br>ACAAGAACAUCAUCAAGAA<br>GCACGAGCCGCAACGAAUA<br>CCGAAACGUUGGUCUGUUA |
| M-003910-00 | MYB | CAACACCAUUUCAUAGAGA<br>CAACGACUAUUCCUAUUAC<br>GAAAUACGGUCCGAAACGU<br>CGGAAGGAGUGGAAGCGUU |
| M-016190-02 | ZNF3 | AGGCUGAUCUCGUAUCUCA<br>AGAAUUACGGGAAUGUGUU<br>GGGAUGAGAUGUUGGCGGC<br>AGACAAUCCUUAACCCUUA |
| M-019779-01 | ZNF91 | CCUCAAAGCUUACUACACA<br>UCAAGUCUUUCUACACAUA<br>UAACAAUCGCAUUAGACUA |

**Table S10:** Sequences of siRNA for selected transcription factors.

**Table S11:** Enrichment of regulons with Gene Ontology Biological Processes. See Excel Supplementary Data.

| MODEL | DATA | NETWORK |
| --- | --- | --- |
| **A. VBSR-VBSR**<br>**Bone Only** | revised-pvalue: 0.00799<br>num-targets: 419<br>num-regulators: **6**<br>regulator-names: CEBPE, GATA1 IKZF1, KLF1, NFE2, MXD1 | 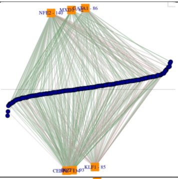 |
| **B. LM-VBSR**<br>**Bone Only** | revised-pvalue: 0.00599<br>num-targets: 451<br>num-regulators: **9**<br>regulator-names: CEBPE, GATA1, IKZF1, KLF1 NFE2, MXD1 GFI1, TAL1 , GFI1B, LYL1 | 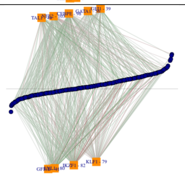 |
| **C. LASSOmin-VBSR**<br>**Bone Only** | revised-pvalue: 0.015<br>num-targets: 257<br>num-regulators: **68**<br>regulator-names: CEBPE, GATA1, IKZF1, KLF1 NFE2, MXD1 ,GFI1, TAL1, FOXD4L1, HOXD11, NPAS2, NR4A2, ZNF518B, IRF4, MYB, GFI1B, ZNF485, TFDP1, SPIB, ZIM2, ZNF446, ZNF558,ZNF726, ZFP62, ZSCAN22, MYBL2, DLX2, HOXD8, FOXI1,ESR1,ZNF146, ZNF239, ZNF283, SP140, SP3, ZNF322, TSC22D4, ZNF229, RELA, ZNF782, FOXG1, MAX,E2F4, TBX6, ZNF48, NR2F6,WIZ, ZNF296, ZNF345, ZNF416, FOXO4, ARID4B, LHX9, HOXD10, OTP, TEAD3, ZFP37, SPI1, NR4A1, SPIC, CREB3L2, ETV1, ZNF286A, ZBTB45, ISX, ZNF75D, NKX2-6, ZNF396 | Not shown |
| **D. VBSR-VBSR**<br>**All Samples** | revised-pvalue: 0.00799<br>num-targets: 303<br>num-regulators: **5**<br>regulator-names: CEBPE, GATA1, KLF1, NFE2, MXD1 | 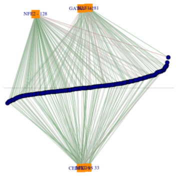 |
| **E. LM-VBSR**<br>**All Samples** | revised-pvalue: 0.016<br>num-targets: 396<br>num-regulators: **8**<br>regulator-names: CEBPE, GATA1, IKZF1, KLF1, NFE2, TAL1 , GFI1B, LYL1 | 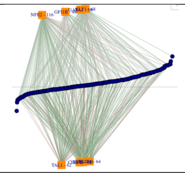 |
| **F. LASSOmin-VBSR**<br>**All Samples** | revised-pvalue: 0.014<br>num-targets: 242<br>num-regulators: **109**<br>regulator-names: CEBPE, GATA1, KLF1,NFE2 MXD1, IKZF1, HES5, MTF1, NFYC, NEUROD1, REL, SATB1, ZNF518B, OTP, MYB, TCF21, TEAD3, ETV1, KAT6A, GFI1B, NFIB, ZNF248, ALX4, SOX6, ETV6, HOXC4, TFDP1, ESR2, TBX6, TOX3, ETV4, ZFP3, ZNF594, NFATC1, ZNF296, ZNF302, ZNF345, ZNF416, MYBL2, ARX, RUNX2, GBX1, HOXA11, ATF7, E2F4, ZFPM1, FOXN1, ZNF563, ZNF567, ZNF571, ZNF585A, HDX, GFI1, HOXD9, ZNF215, ZNF229, MAFF, NFE2L3, THAP8, ZNF536, DLX5, NFIL3, NR4A3, SPIC, POGZ, NR4A2, OSR1, TLX2, UBP1, ZNF620, HMX1, THAP6, NKX2-5, FOXH1, ZBTB43, ZNF268, TOX4, SALL1, ZNF174, RFXANK, ZBTB45, ATRX, GRHL3, TAL1, ZNF879, NFYB, ZNF668, NR2F6, ZBTB32, ZNF283, FOXO4, MTA3, PURB, RXRA, MXI1, SPI1, ZNF233, ZNF235, TSC22D3, IRF4, OSR2, DEAF1, NANOG, LOC145783, HOXB3, TBX2, ZFP28, ZNF568, ID1 | Not shown |

**Table S12: Consistent Discovery of P-M1**: Identifying the most strongly rewired regulatory module from the 46 bone-only samples (**A-C**) resulted in similar regulatory networks, with the same core drivers (in colored text), regardless of the regression model used to construct the co-expressed modules. This finding also held for using the data from all 68 patient samples (**D-F**).

### S3 Additional Figures

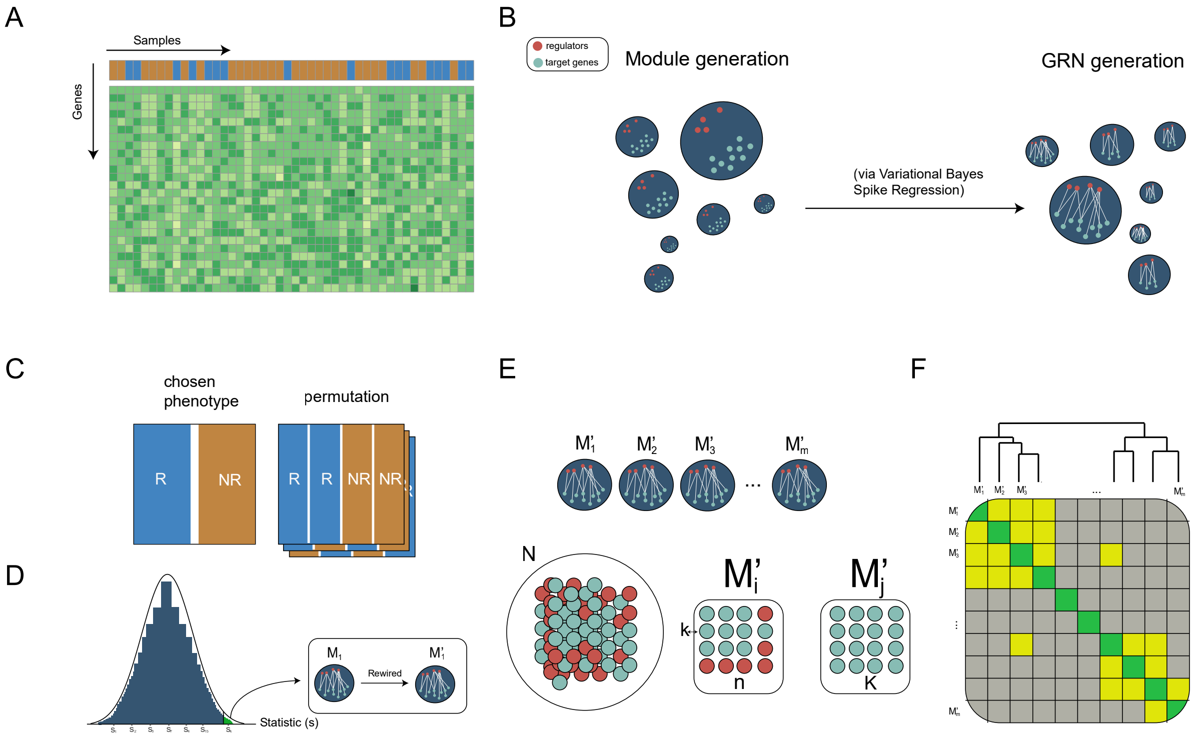

**Figure S1: Workflow of TraRe. A:** Example of a gene expression matrix (genes by samples). Color of samples indicates whether the patients are labeled as responders or non-responders. **B:** GRN inference process in two steps, module generation and GRN generation. **C:** GRN rewiring process. A permutation test on sample class labels is performed per module. **D:** A dissimilarity metric is evaluated for the true class labels and compared to a fixed amount of permutations. **E:** Robust rewired GRN inference process begins with Hypergeometric tests performed between rewired modules. **F:** Similar rewired modules are grouped via a hierarchical clustering yielding robust, regulatory modules.

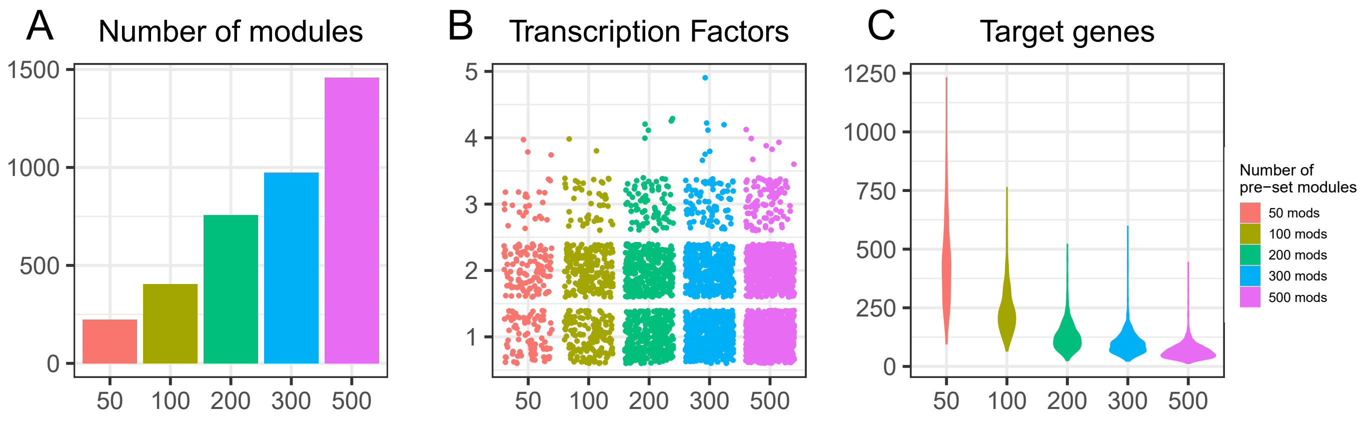

**Figure S2: Variation of sub-module generation**. We modified the number of the pre-set sub-modules to 50, 100, 200, 300 and 500 on PROMOTE data and run 5 bootstraps.Variation of sub-module generation. We modified the number of the pre-set sub-modules to 50,100,200,300 and 500 on PROMOTE data and run 5 bootstraps. We show the influence in the number of sub-modules generated (**A**), the number of transcription factors per generated sub-module (**B**) and the number of targets per generated sub-module (**C**).

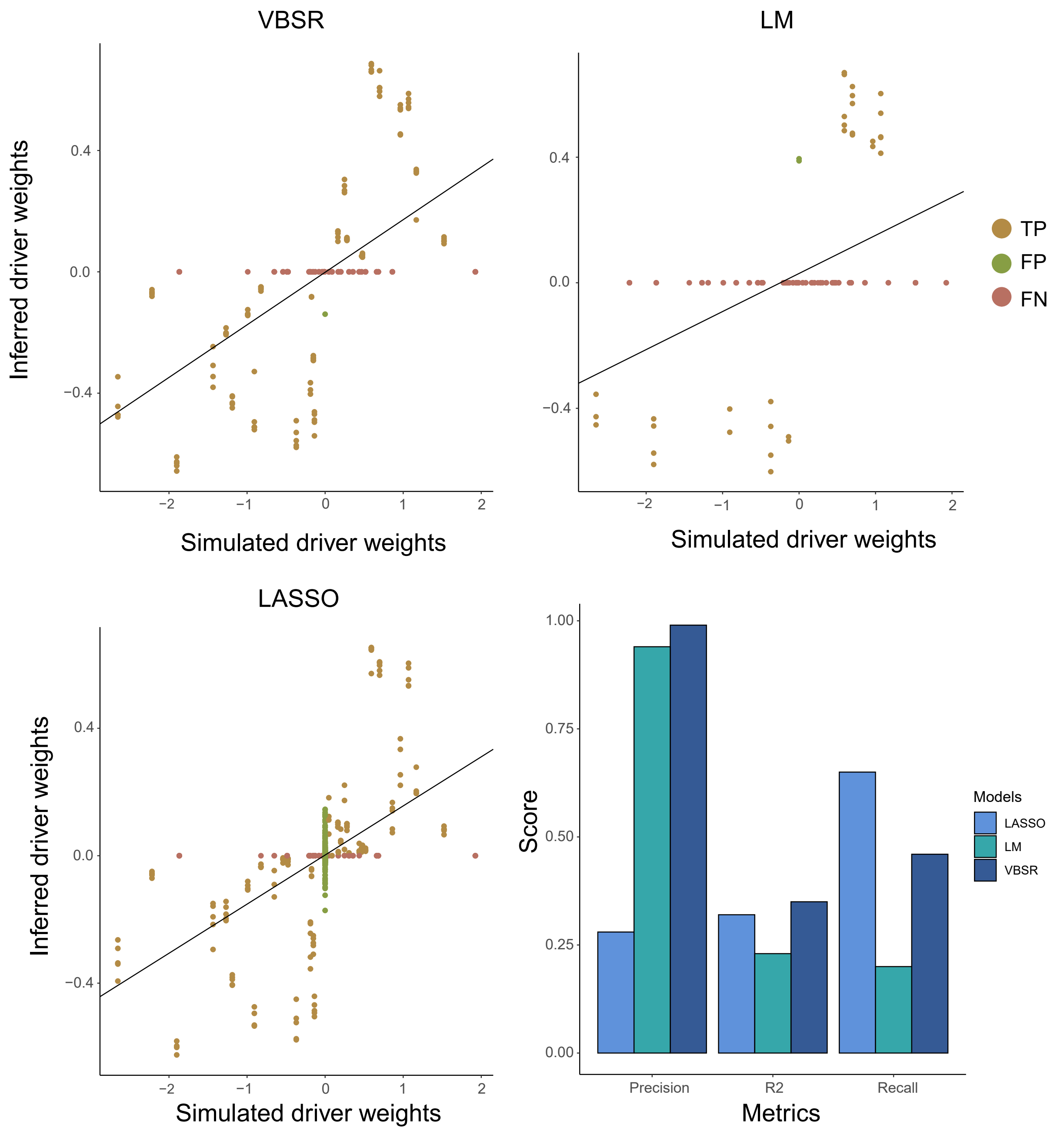

**Figure S3: Classification metrics along VBSR,LM and LASSO**. For a fixed Jaccard index, variance noise $\sigma$, number of samples and p-value thresholds, TP, FP and FN from simulated-inferred modules pairs are shown for each model. A linear model has been fitted to explain driver's regulatory programs within inferred GRNs based on simulated regulatory programs. Moreover, $R^2$ adjusted, precision and recall parameters from the fitted linear model are shown. VBSR obtained perfect precision, which was almost scored by LM, and also obtained the largest $R^2$. Largest recall score was obtained by LASSO model, as it over-inferred drivers, which can be seen in the form of FPs at its precision score.

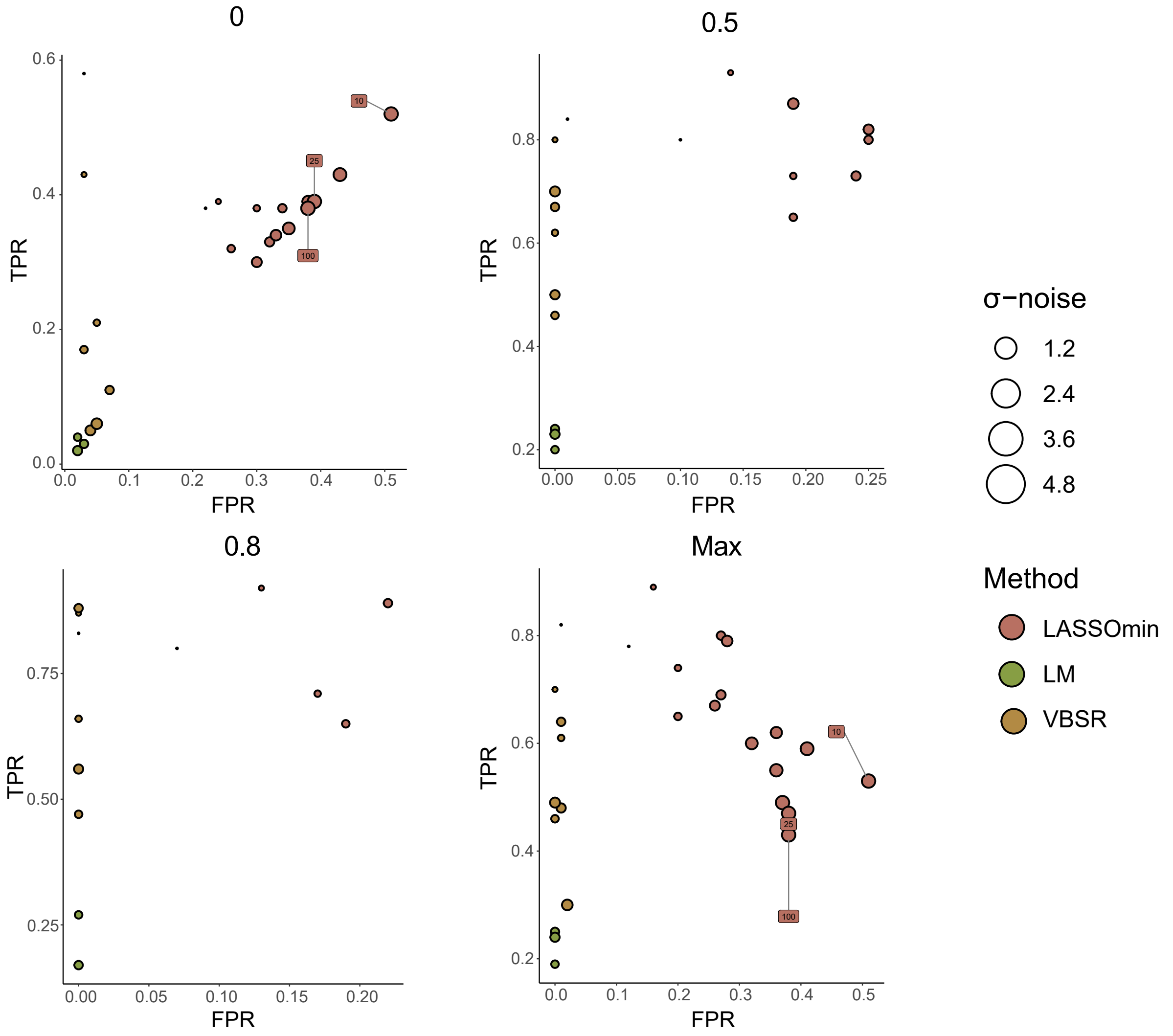

**Figure S4: Discrete ROC's curves comparing simulated and inferred modules as a function of the simulated modules variance noise $\sigma$.** For each defined Jaccard index threshold and each value of $\sigma$ = [0.01, 0.4, 0.8, 1.2, 1.6, 2, 2.4, 2.8, 3.2, 3.6, 4, 4.4, 4.8, 10, 25, 50, 100], TPR and FPR have been computed along VBSR, LASSO and LM. Default number of samples and p-value thresholds have been used. As the Jaccard index threshold decreases (more restrictive) the fitting models are less robust to noise variance. Also, VBSR achieved large TPR to FPR ratios in a wide range of noises (up to $\sigma$=2.8).

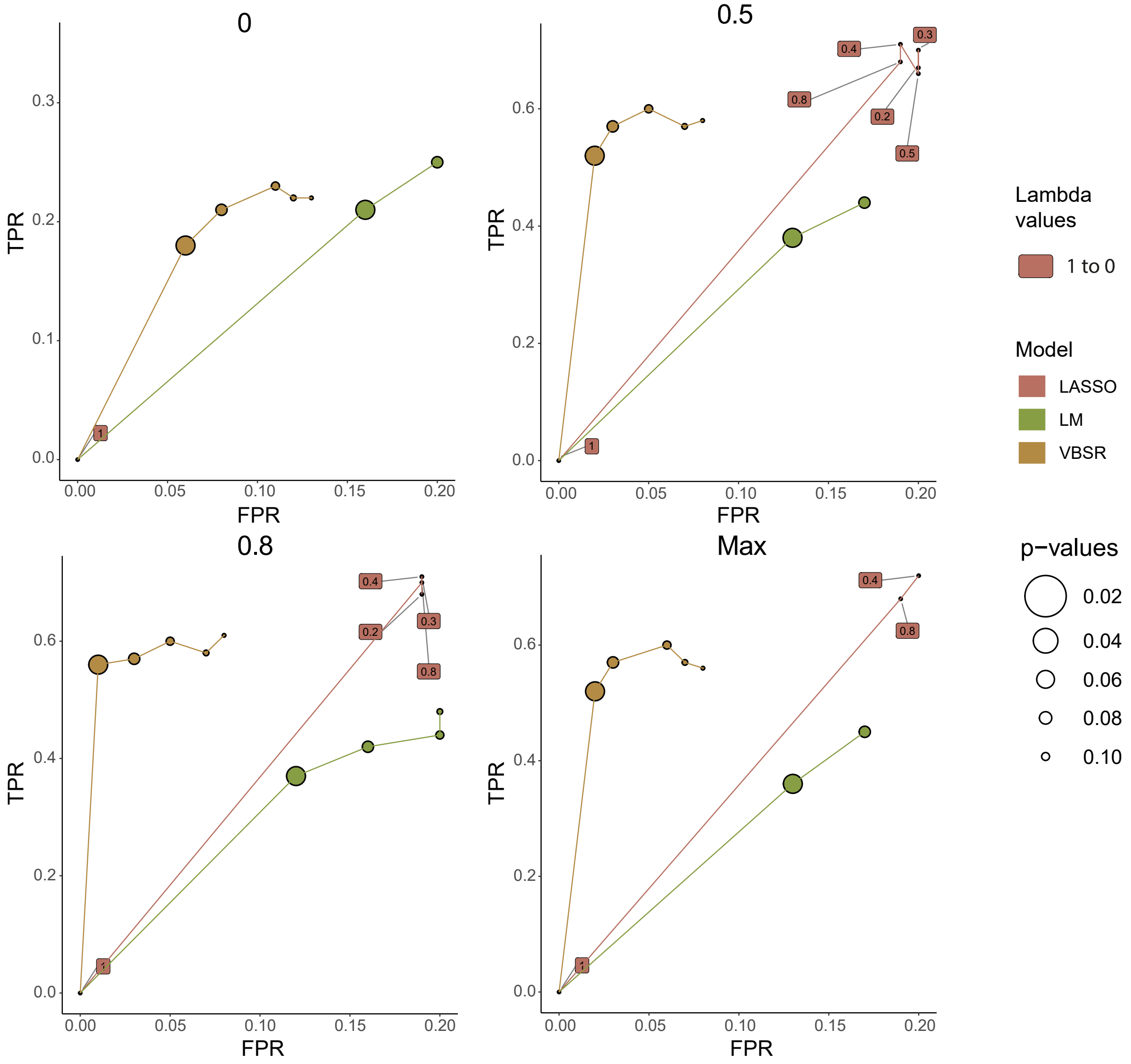

**Figure S5: ROC's curves as a function of p-value thresholds**. TPR and FPR have been computed along a desired p-value's range (0.02 to 0.14) for each Jaccard index threshold and fitting models VBSR, LASSO and LM. Noise $\sigma$ and number of samples have been set to its default parameters. Note that, for the LASSO model, Lambda parameter has been modified as a metric of sensitivity, selecting Lambda's deciles from LASSO cross validation model. (R's *cv.glmnet* package). VBSR achieves the largest TPR to FPR ratio on every Jaccard index threshold.

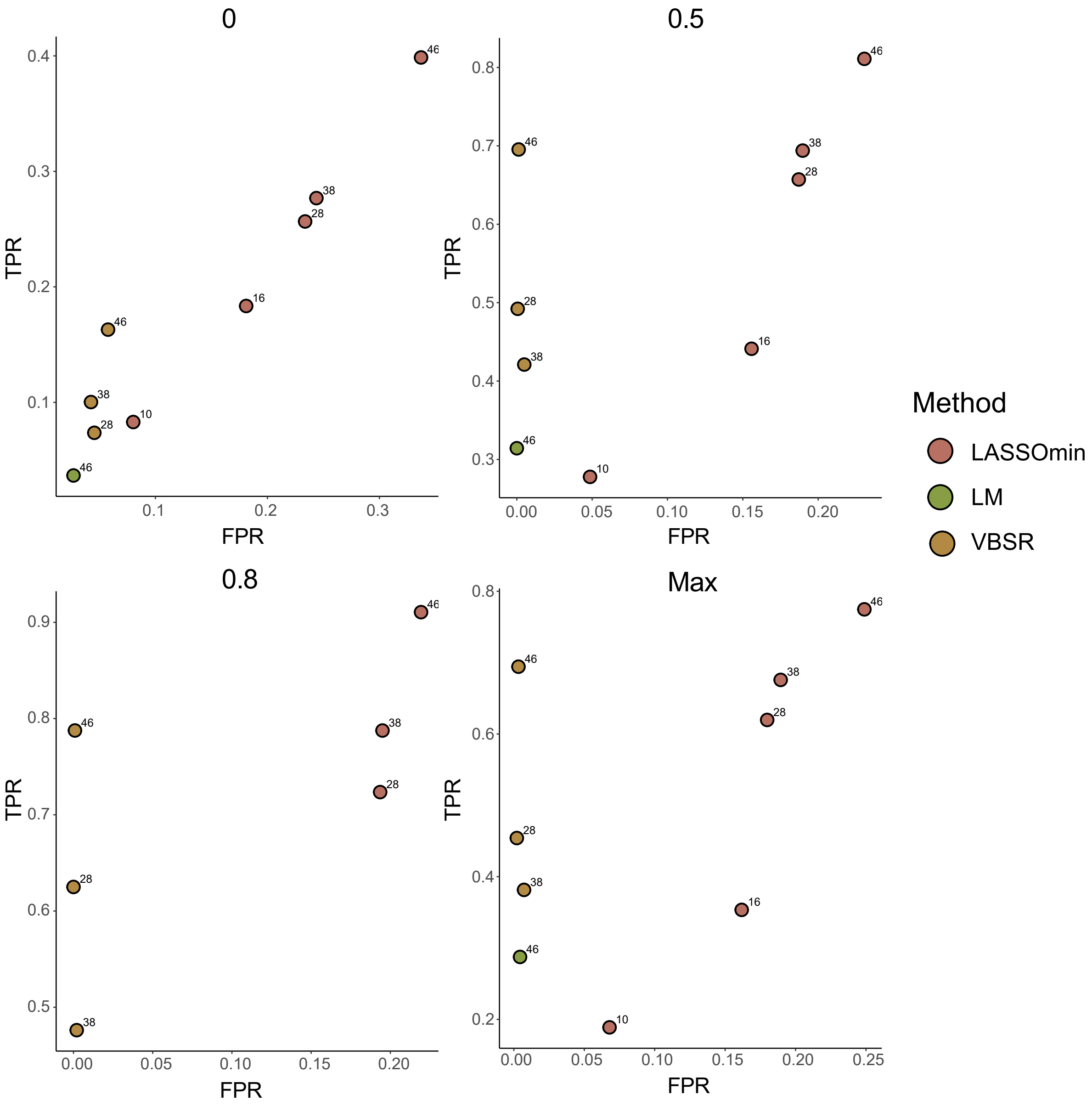

**Figure S6: ROC's curves a function of samples (patients)**. TPR and FPR have been computed for each Jaccard index threshold and fitting models VBSR, LM and LASSO, along 10, 16, 28, 38 and 46 samples. Noise $\sigma$ and p-value's threshold have been set to its default parameters (see Methods). Missing points when threshold = 0 are due to fitting errors from specific models. Missing points distinct to the aforementioned in the remaining thresholds are due to the filtering. Above 28 samples, VBSR achieved largest TPR to FPR ratio on every Jaccard index threshold. LASSO was the only model able to infer GRNs under 28 samples, maintaining similar TPR to FPR ratios of larger samples runs.

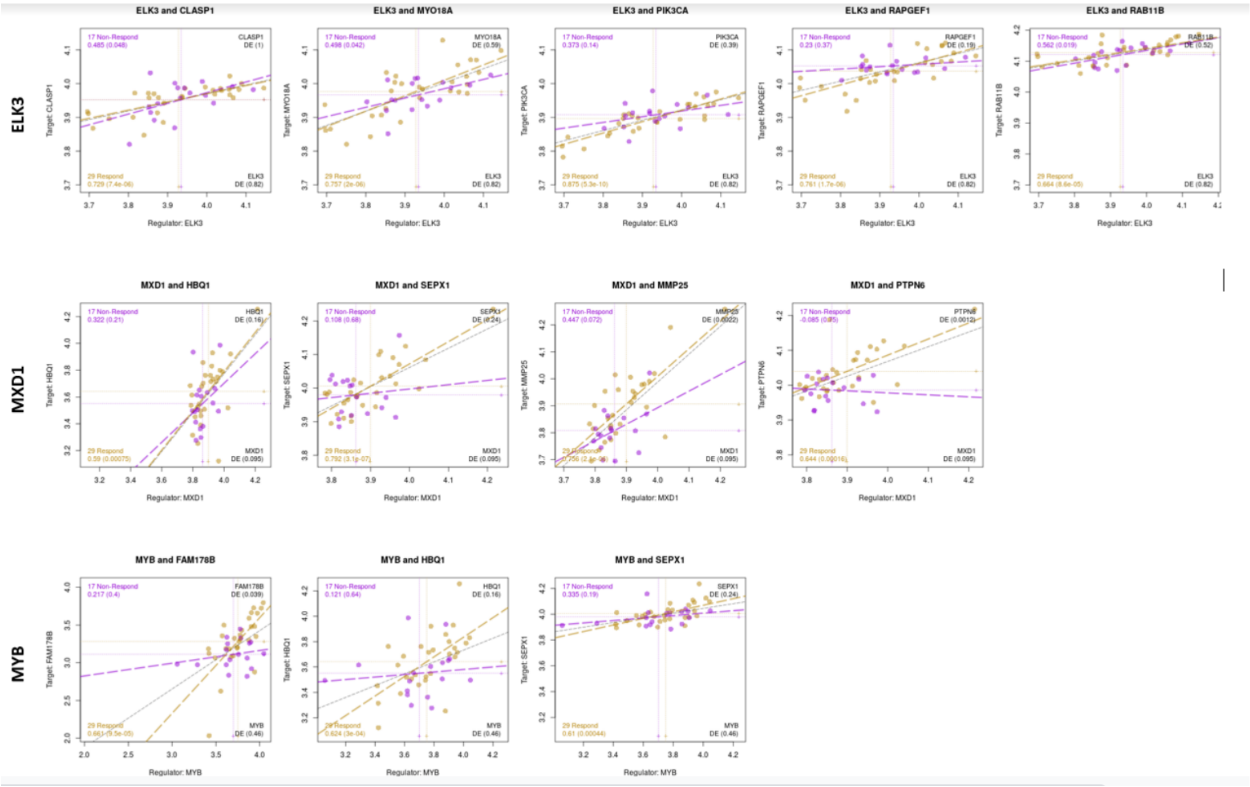

**Figure S7: Expression Patterns in PROMOTE Data.** For each of the reported experimental validations, we show the corresponding expression information from the original PROMOTE data. Points are plotted for responders (yellow) and non-responders (purple) with best fit lines drawn and correlations and their significance reported on the left hand side.

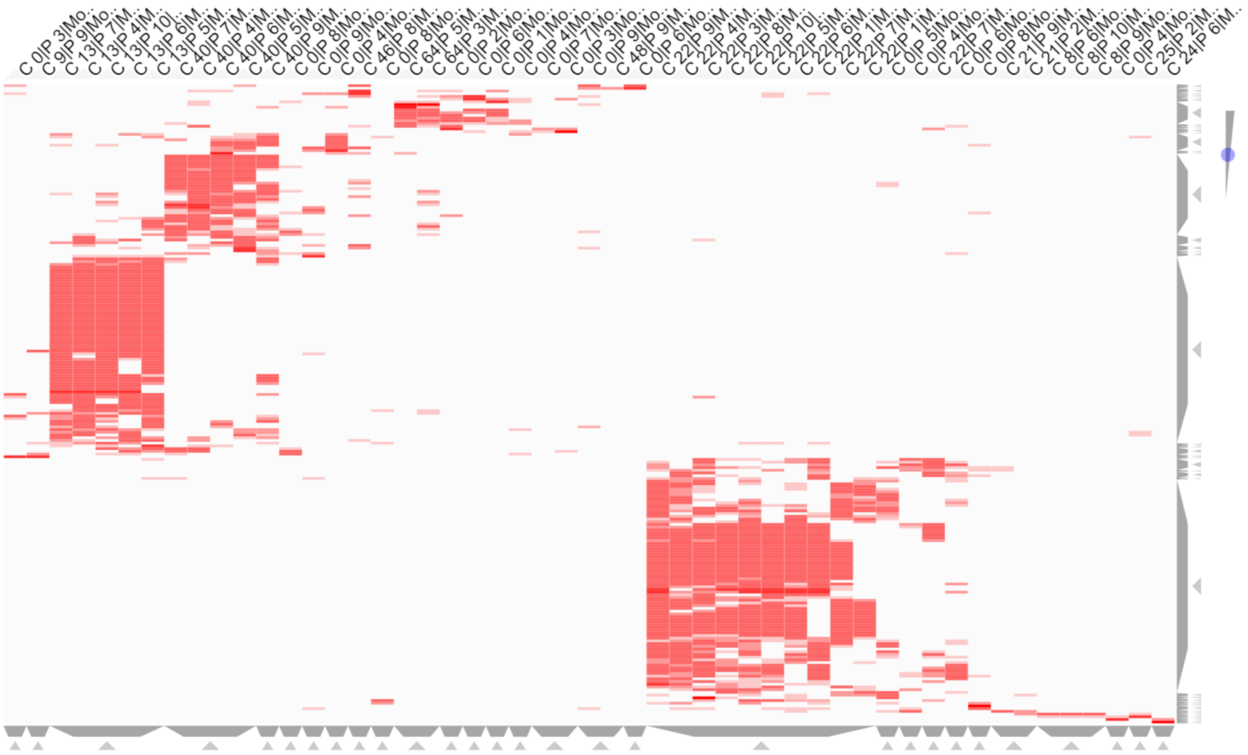

**Figure S8:** Heatmap showing the 51 rewired modules as columns labeled by their community membership (C_##, C_0 for not a member of any community) and their bootstrap run (P_##). The rows are gene targets that are in at least four of the rewired modules or regulator TFs in at least one. The red color of the cell indicates membership of the gene in the module (dark red indicates the gene is a regulator of that module). Figure generated with Clustergrammer[73] web tool.

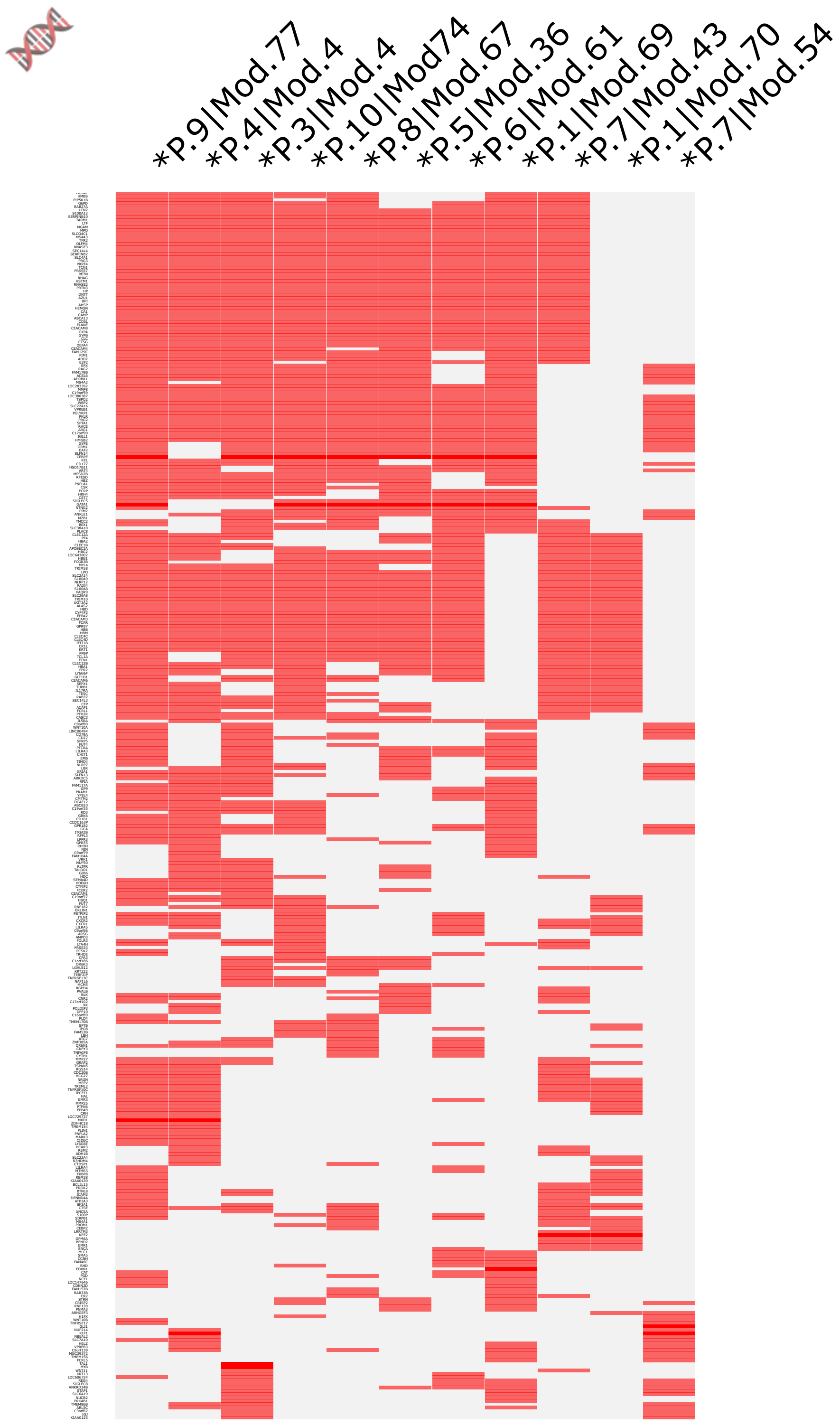

**Figure S9:** Heatmap showing the 11 modules of P-M1 as columns labeled by their bootstrap run (P.##) and within bootstrap submodule (Mod.##) and flagged with "*" if rewired. The rows are gene targets that are in at least two of the rewired modules or regulator TFs in at least one. The red color of the cell indicates membership of the gene in the module (dark red indicates the gene is a regulator of that module). Figure generated with Clustergrammer[73] web tool.

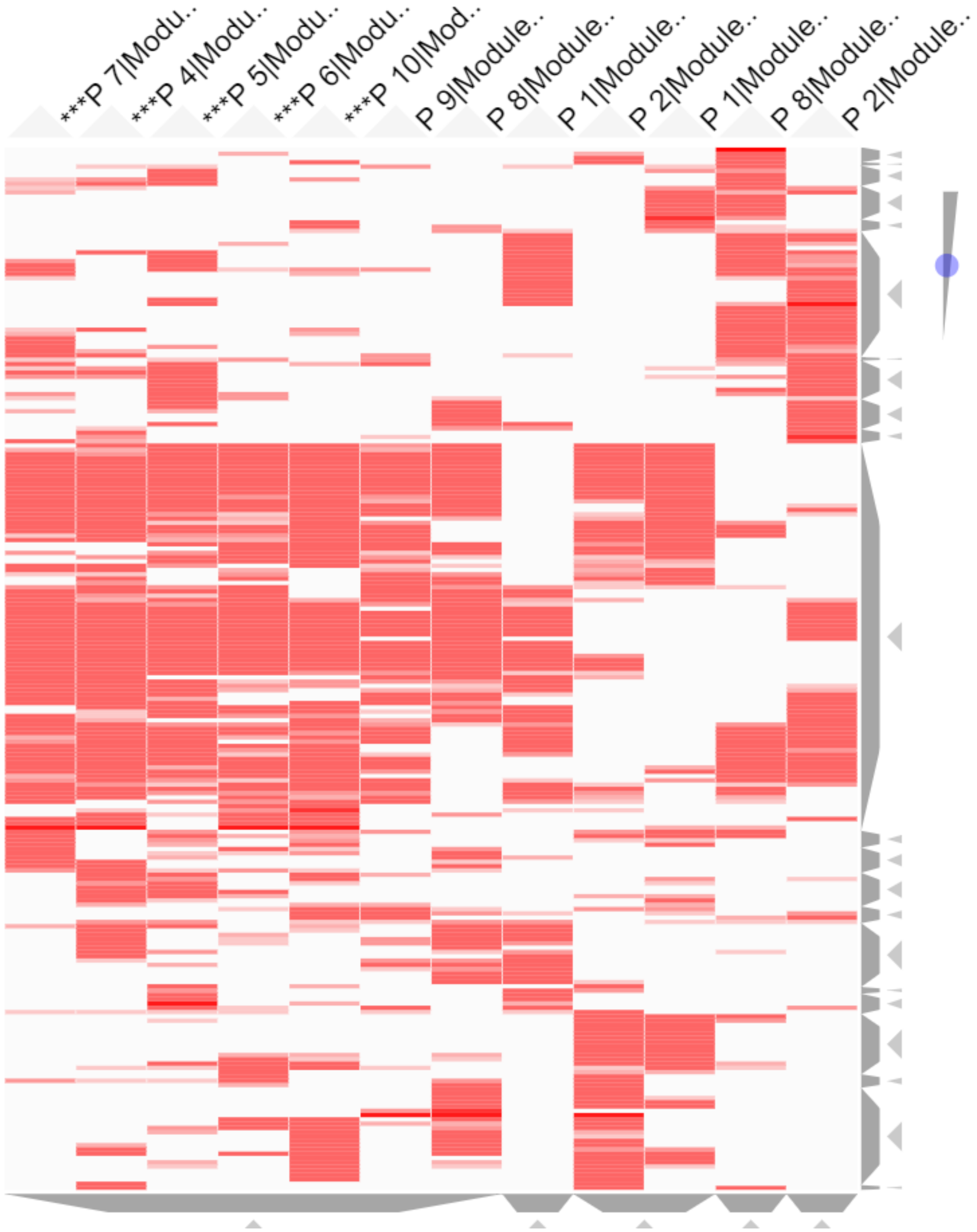

**Figure S10:** Heatmap showing the 12 modules of P-M2 as columns labeled by their bootstrap run (P_##) and flagged with "***" if rewired. The rows are gene targets that are in at least two of the rewired modules or regulator TFs in at least one. The red color of the cell indicates membership of the gene in the module (dark red indicates the gene is a regulator of that module). Figure generated with Clustergrammer[73] web tool.

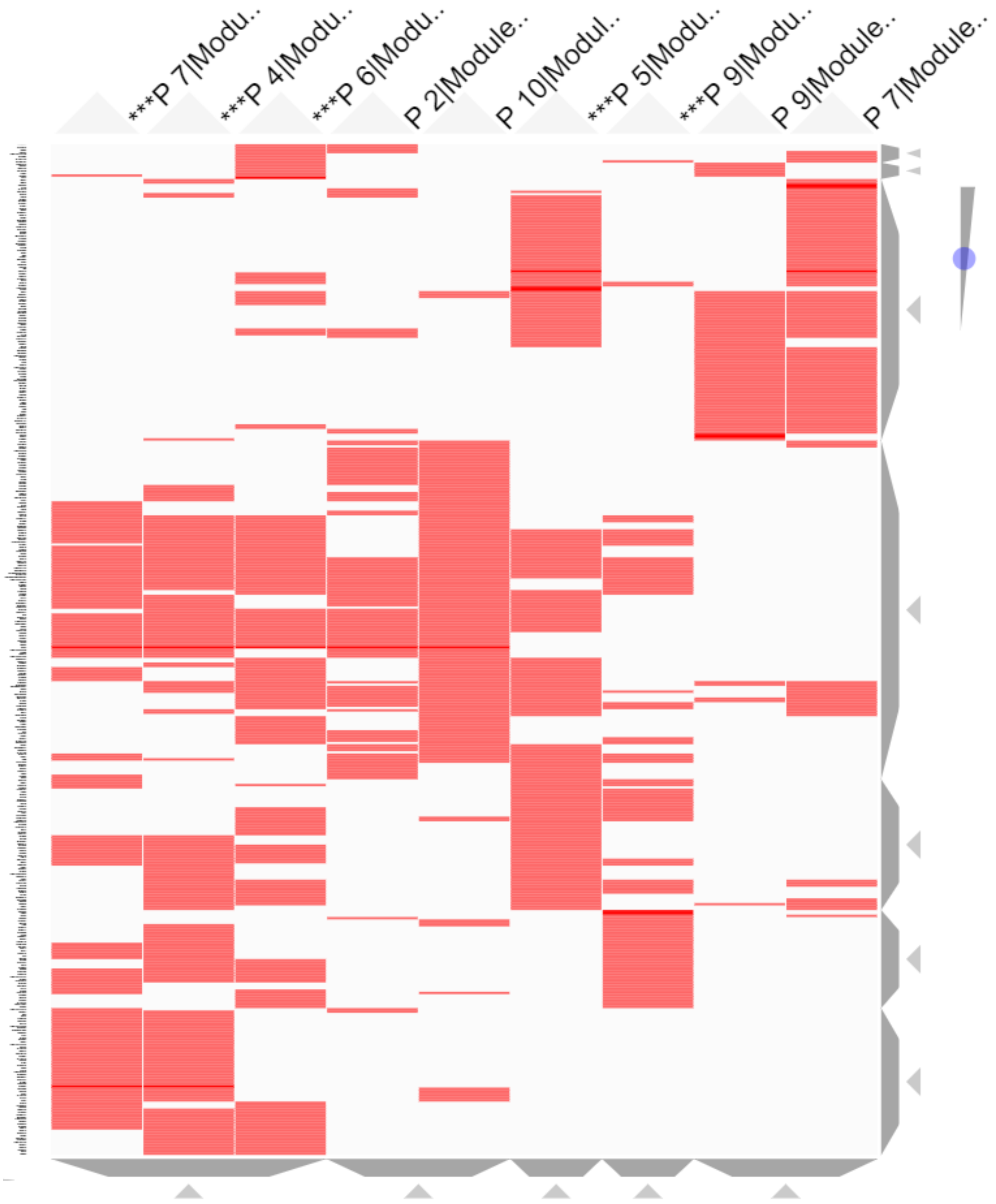

**Figure S11:** Heatmap showing the 9 modules of P-M3 as columns labeled by their bootstrap run (P_##) and flagged with "***" if rewired. The rows are gene targets that are in at least two of the rewired modules or regulator TFs in at least one. The red color of the cell indicates membership of the gene in the module (dark red indicates the gene is a regulator of that module). Figure generated with Clustergrammer[73] web tool.

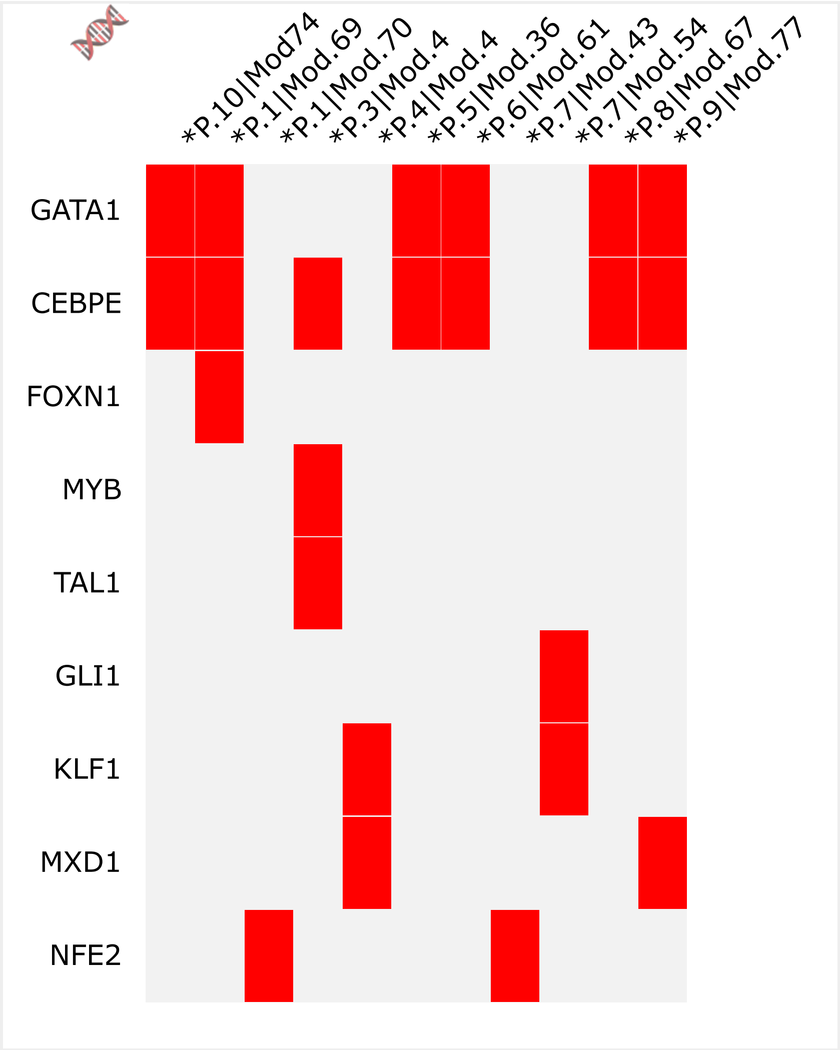

**Figure S12:** Regulators of P-M1 shown as rows of a membership heatmap with the 11 modules of P-M1 as columns labeled by their bootstrap run (P.##) and within bootstrap submodule (Mod.##) and flagged with "*" if rewired. Figure generated with Clustergrammer[73] web tool.
